# stVCR: Spatiotemporal dynamics of single cells

**DOI:** 10.1101/2024.06.02.596937

**Authors:** Qiangwei Peng, Peijie Zhou, Tiejun Li

## Abstract

Time-series spatial transcriptome data with single-cell resolution provide an opportunity to study cell differentiation, proliferation and migration in physical space over time. Due to the destructive nature of sequencing, reconstruction of spatiotemporal dynamics from data remains challenging. Especially, the inference of migration in physical space remains a difficult task, as samples obtained at different temporal snapshots might not be in the same coordinate system due to the difference of biological replicates. Here we developed stVCR, a generative deep learning model, which integrates the dynamical optimal transport (OT) with the unbalanced setting, the density matching invariant to rigid body transformations as well as priors to model known biology and preserve spatial structure. stVCR achieves the end-to-end simultaneous reconstruction of continuous cell differentiation, proliferation, physical space migration, and spatial coordinates alignment from spatial transcriptome snapshots. In addition, stVCR allows the interpretable study of complex interactions between cell phenotype transition, spatial migration and proliferation. Through benchmarking on both simulation data and real datasets, we validated the effectiveness and robustness of stVCR and demonstrated its advantages over static OT or linear interpolation methods. We applied stVCR to dissect spatiotemporal dynamics underlying axolotl brain regeneration and 3D Drosophila embryo development.

## Introduction

The development of a fertilized egg into a complete embryo is a highly complex and important process in biology [1–4]. This process involves intricate interactions between the dynamic regulation of gene expression, cell differentiation, cell division, apoptosis, as well as cell migration within physical space [5, 6].

The advent of spatial transcriptome (ST) technology has allowed obtaining both gene expression data and spatial coordinates [7–11]. As technology advances, spatial resolution has reached the single-cell or even subcellular level, exemplified by methods such as Stereo-seq [10] and 10x Visium HD [11]. However, due to the destructive nature of sequencing, ST data can only provide snapshots rather than a continuous trajectory. If ST sequencing technology is likened to an ultra-wide-angle camera, it can take pictures of living organisms but lacks video recording capability. Especially, when sequencing at multiple time points during embryonic development, the resulting time-series ST data often come from different biological replicates, therefore yielding multiple unpaired snapshots.

Recovering cells’ dynamic trajectories from single-cell sequencing data or ST data is a challenging task. RNA velocity [12] utilizes unspliced/spliced RNA to infer the developmental direction of each cell. This inspired a series of subsequent works using unspliced/spliced RNA to more accurately infer RNA velocity [13–18]. These methods suffer from scale invariance due to the lack of temporal information [19]. Metabolic labeling scRNA-seq introduces temporal information into the data by distinguishing new/old RNA [20–27]. Dynamo [28] designed parameter inference methods for metabolic labeling scRNA-seq data based on steady-state assumptions and deterministic models, and Storm [29] extended it to be independent of steady-state assumptions and stochastic models. Time-series scRNA-seq data introduces temporal information into the data in another way. Waddington-OT [30] pioneered the use of optimal transport (OT) for modeling time-series scRNA-seq data, finding the optimal mapping in cells at two adjacent time points. However, it approximates cell proliferation by using growth hallmark genes, which largely depends on the choice of database.

Targeted for time-series scRNA data, various extensions of OT haven been formulated, with a comprehensive review recently presented in [52]. TrajectoryNet [31, 32] combines dynamical OT [33] and normalizing flows to infer continuous trajectories of cells and takes cell proliferation into account through a separated static OT. MIOFlow [34] follows the geometry by operating in the latent space of a geodesic autoencoder. OT-CFM [35] is a minibatch OT-based simulation-free method for training continuous normalizing flows (CNFs), which enhances stability, training efficiency and inference speed. Some work considers stochastic cellular dynamics, such as AM [37], WLF [38], FBSDE [39] and PI-SDE [40]. SF2M [36] further extends the inference of stochastic dynamics in simulation-free setups. To account for cell proliferation and death effects, UDSB [41] infers unbalanced Schrödinger bridges under the given prior knowledge. TIGON [42] uses dynamical unbalanced OT [43] to reconstruct dynamic trajectories and population growth simultaneously. DeepRUOT [44] extends even further to the stochastic unbalanced case. In addition, CellOT [45] uses OT to model single-cell perturbation responses. Since the usual scRNA-seq data do not include the spatial coordinates of the cells, these methods have limitations Since the usual scRNA-seq data do not include the spatial coordinates of the cells, these methods have limitations in modeling cell migration in physical space directly.

The availability of time-series ST data has made it possible to study how cells migrate in physical space along with their state transition dynamics. PASTE [46] uses fused Gromov-Wasserstein(GW) OT [47] to align 2D adjacent tissue ST slices to reconstruct the 3D structure of the tissue. Moscot [48] uses a similar fused GW-OT to find the optimal mapping of cells between slices at two adjacent time points, incorporating penalties for unbalanced and entropy regularization and employing a low-rank OT [49] to accommodate larger data sizes. Spateo [50] aligns the spatial coordinates of two adjacent time points by optimal mapping to obtain cell migration velocity and then learns a vector field of continuous spatial coordinates. However, it does not fully address the interplay between gene expression and spatial location, and processes such as cell division and apoptosis. DeST-OT [51] considers how to model cell proliferation in the static OT setting, in particular ST data. In addition, TopoVelo [53] uses spatial coordinates to model cellular neighborhoods when inferring usual RNA velocity based on unspliced/spliced RNA, and designs post-processing steps to infer cell migration velocity. STT [54] characterizes multistability in space by integrating unspliced/spliced RNA and ST through a multiscale dynamical model.

Reconstructing dynamical trajectories of cell differentiation, proliferation, and migration in physical space simultaneously for time-series ST data is a challenging task. Especially to quantify the migration in physical space, improper treatment might introduce pseudo movements of cells as the cell coordinates obtained at different temporal snapshots are not in same coordinate system. Analogous to recovering a video from multiple photos, we aim to reconstruct the entire cellular developmental dynamics from multiple unpaired ST snapshots, thus obtaining a continuous spatiotemporal developmental trajectory. To achieve this goal, we developed an algorithm called <u>s</u>patio-<u>t</u>emporal <u>V</u>ideo <u>C</u>assette <u>R</u>ecorder (stVCR), which is a dynamical optimal transport algorithm for resolving the issue of alignment of ST section data and unbalanced populations at different snapshots, and incorporation of biological structure priors in an integrative manner. As the result, stVCR reconstructs the spatiotemporal dynamical process for the considered system from multiple ST snapshots. Furthermore, stVCR also reveals the complex regulatory mechanisms behind the overall cellular dynamics, including how gene expression and spatial location affect each other, and how they affect cell proliferation.

## Results

### Overview of stVCR

In stVCR, we develop a new dynamical OT formulation with special treatments for different aspects of data for more biologically meaningful spatiotemporal matching task (Fig. 1A). Specifically, for gene expression, we use the Wasserstein OT to model the temporal coupling of distributions (Fig. 1A Left and Methods). For spatial coordinates of cells, since rotations and translations may be involved to prevent a direct comparison of cell coordinates at different instants, we use the rigidbody transformation invariant OT to make the spatial alignment in time (Fig. 1A Middle and Methods). For the number of cells, due to the cell division and apoptosis, we use the unbalanced OT to model the unbalanced populations (Fig. 1A Right and Methods). Additionally, stVCR optionally takes known cell type transition prior (Fig. 1B Left) as well as the spatial structure preserving prior for specific cell types (or organs) (Fig. 1B) to produce more biologically meaningful results (Methods). We unified all three necessary modules and two optional modules, allowing us to study how a population of cells changes in gene expression, how they migrate in physical space, and how they divide and apoptose over time (Figure 1A,B,C and Methods). We take the spatial coordinates of cells at the first time point *t* = *t*_0_ as the reference coordinates system, and the state of considered cell group at time *t* is described by a time-dependent distribution *ρ*_*t*_(***x, q***), where *ρ*_*t*_ depends on the spatial coordinates 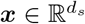 (*d*_*s*_ = 2 or 3) of the cells in the reference coordinates system and the gene expression variable 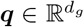 after dimensionality reduction. stVCR finds the optimal rigid rotation matrix *R*_*k*_ and translation vector ***r***_*k*_ for the coordinate system at time point *t* = *t*_*k*_ except *t*_0_, and uses the transport-with-growth PDE

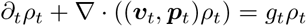

to interpolate the empirical density 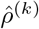 and cell number *n*_*k*_ of the snapshot spatial transcriptomic data at *t* = *t*_*k*_ after rigid body transformation (Fig. 1C), where ***v***_*t*_, ***p***_*t*_, and *g*_*t*_ are parameterized to be learned by neural networks. Biologically, ***v***_*t*_ = d***x****/*d*t* describes the migration velocity of cells in physical space, ***p***_*t*_ = d***q****/*d*t* describes the RNA velocity of cells in gene expression space, and *g*_*t*_ *∈* ℝ describes cell proliferation (Fig. 1C, D).

**Fig. 1:**
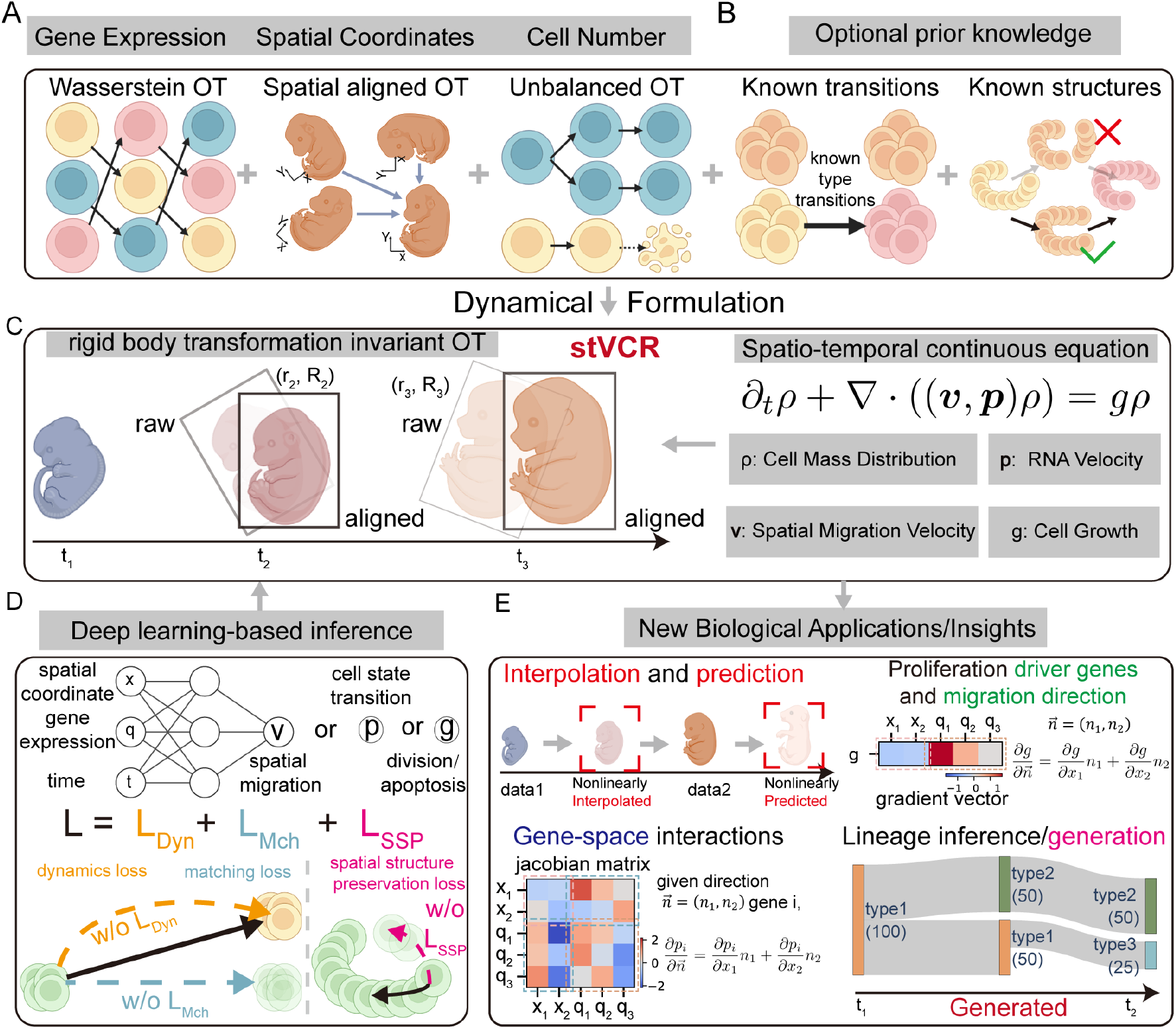
Overview of stVCR. **A**. stVCR adopts dynamical OT framework yet with special treatments for different types of data in the spatial transcriptome. Specifically we use Wasserstein OT for gene expression data (Left), rigid body transformation invariant OT for spatial coordinates (Middle), and unbalanced OT for cell number change due to cell division and apoptosis (Right). **B**. stVCR can optionally model prior knowledge, including biological priors for known type transitions (Left) and spatial structure preserving priors (Right). **C**. stVCR unifies the three necessary modules and two optional modules into a dynamical OT. The input spatial transcriptome snapshots are described as distributions *ρ*^(*k*)^, and the permissible rotations and translations are characterized by (*R*_*k*_, ***r***_*k*_) at *t* = *t*_*k*_. The modeling density *ρ*_*t*_ is governed by a partial differential equation involving spatial velocity ***v***, RNA velocity ***p***, and growth rate *g*. **D**. stVCR solves the problem in **B** based on deep learning. ***v***_*t*_(***x, q***), ***p***_*t*_(***x, q***) and *g*_*t*_(***x, q***) are modeled by three neural networks. The loss function includes three parts: dynamics loss, matching loss and spatial structure preserving loss. **E**. stVCR can perform downstream analyses, including interpolation and prediction (Top left), studying cell-specific gene-gene, gene-space, space-space interactions (Bottom left), exploring the cell-specific effects of gene expression and spatial variations of growth rates (Top right) and inferring temporal cell-type developmental lineages (Bottom right).

To train the spatiotemporal dynamics, stVCR parameterizes the 2D rotation matrix by using rotation angle (or Euler angle for 3D case), and simultaneously finds the optimal rigid body transformations and parameterized dynamics by minimizing a total loss composed of dynamics loss, matching loss and optional spatial structure preserving loss

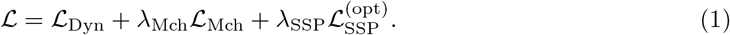

The dynamics loss *ℒ*_Dyn_ further contains three parts

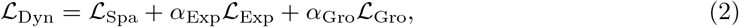

promoting the least consumption of kinetic energy of spatial migration and gene expression change, and growth energy, respectively. The matching loss *ℒ*_Mch_ promotes the cell dynamics to match the aligned ST data as well as possible at observation time points, and the *optional* spatial structure preserving loss 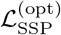 promotes a stable spatial structure for the the user-specified organ or cell type by encouraging adjacent cells to have similar spatial velocities, thereby preventing arbitrary deformations (Fig. 1D and Methods). The training process involves OT optimization and integrating ODEs represented by neural networks, which we solve using the POT [55] and torchdiffeq packages [56], respectively.

Once we have completed the entire training process to obtain the optimal rigid-body transformation and parameterized dynamics, we can first apply the optimal rigid-body transformation to align the spatial coordinates of cells at different time points to the reference coordinate system, and then perform a series of downstream analyses (Fig. 1E and Methods). (1) Interpolation and prediction. We evolve forward or backward from the nearest observations to the interested time point (between observations or in the future) based on learned cellular dynamics (Fig. 1E Top left and Methods). (2) Gene-space interactions. We study cell-specific gene-gene, gene-space and space-space interactions by calculating the Jacobian matrices of learned spatial migration dynamics and gene expression dynamics and further calculating the directional derivatives along the cell migration direction of interest (Fig. 1E Bottom left and Methods). (3) Proliferation driver genes and migration direction. We study the effects of genes and migration on growth by calculating the gradient of the learned growth dynamics and further calculating the similar directional derivatives (Fig. 1E Top right and Methods). (4) Lineage inference/generation. For originally annotated data, we can infer temporal developmental lineages by learning a time-dependent classifier to annotate unobserved cells generated by interpolation or prediction (Fig. 1E Bottom right and Methods).

### Benchmark experiments for accuracy, scalability and robustness of stVCR

To demonstrate the necessity of aligning the spatial coordinates of different temporal snapshots into the same coordinate system, and benchmark the ability of the stVCR to recover spatiotemporal dynamics and reveal key regulatory mechanisms, we generated the simulated dataset of gene circuits and two spatial dimensions (Fig. 2A, B and Methods). The three genes are named *Red, Green* and *Blue* genes. There are regulatory relationships between different genes and different spatial coordinates, in addition, gene expression and cell migration also affect cell proliferation (Fig. 2A). *Red* and *Green* genes form a toggle switch circuit and they have opposite effects on growth. In addition, the difference in spatial location makes them unequal in status (Fig. 2A, B and Methods).

**Fig. 2:**
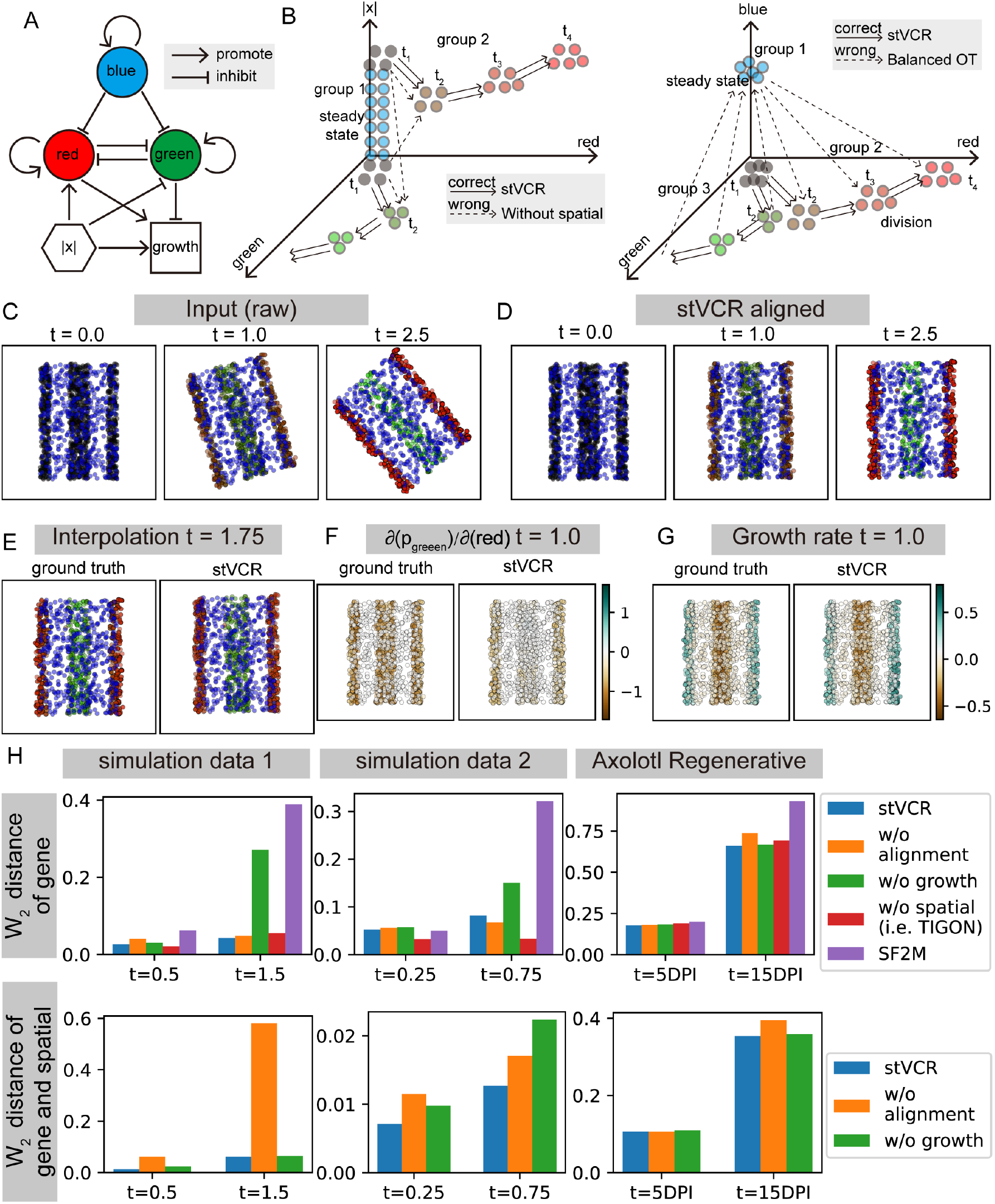
Benchmark of stVCR. **A**. Regulation relationship diagram to generate simulation data. **B**. Dynamic diagram of cell evolution over time. Left: Dynamics of the second group and third group of cells over time in *r* (red gene expression), *g* (green gene expression) and |*x*| (absolute values of spatial coordinates) dimension. Right: Dynamics of all cells over time in *r, g* and *b* (blue gene expression) dimension. **C**. Input data to stVCR at *t* = 0.0, 1.0 and 2.5. The color was determined by the expression of three genes *Red, Green* and *Blue*. **D**. The aligned results at *t* = 0.0, 1.0 and 2.5 of cells at *t* = 0.0 using stVCR. **E**. Results of stVCR interpolation at *t* = 1.75 and comparison with ground truth. Left: ground truth; Right: stVCR. **F**. Derivative of *Green* gene velocity with respect to *Red* gene at *t* = 1.0 of true dynamics and learned dynamics. **G**. Same as **F**, but for growth rates of cells. **H**. The results of the leave-one-time-point-out benchmark test. From left to right, the panels correspond to the first simulated data, the second simulated data and the axolotl brain regeneration data. The first row shows the Wasserstein distance in gene expression space. The second row shows the Wasserstein distance that includes both gene expressions and spatial coordinates.

In the simulated dataset, there are three groups of cells at the initial moment (Fig. 2 B). The first group highly expressed gene *Blue* and was in steady state (Fig. 2 B). The second and third groups had similar low gene expression at the initial moment, but their different spatial locations determined their different fates of transition and growth (Fig. 2B). Without spatial information, it is not possible to distinguish between the second and third group at the initial moment, which would lead to erroneous trajectories (Fig. 2B Left and Fig. S5 last row). If cell proliferation is ignored, it will lead to incorrect cell trajectories of the small number group to the large number group (Fig. 2B Right and Fig. S6 last row). The input data totaled 6 time points, and we rotated the spatial coordinates by different angles to simulate the possible rotation of tissues by spatial transcriptome sequencing (Fig. 2C and Fig. S1A).

To illustrate the ability of stVCR to align the spatial coordinates of different temporal snapshots and reconstruct the entire spatiotemporal dynamics, we took the data from the first time point and evolved them according to the learned dynamics, demonstrating consistency with real dynamics (Supplementary Video 1; Fig. 2D and Fig. S1B). Specifically, the first group of cells remained virtually unchanged. The second group of cells gradually overexpressed the *Red* gene, moved outwards in the horizontal direction, and continuously proliferated. The third group of cells gradually overexpressed *Green* gene and continued apoptosis. In addition, we observed that in the aligned space by stVCR, cells only moved horizontally and did not rotate, indicating that we found the optimal rigid body transformation to align the data at different time points while finding the optimal dynamics of cell evolution (Supplementary Video 1, Fig. 2D and Fig. S1B). In addition, stVCR interpolated the unobserved intermediate moments *t* = 0.25, 0.75, 1.25, 1.75 and 2.25 and predicted the future moments *t* = 2.75 based on the learned dynamics, and the results are close to ground truth (Fig. 2E and Fig. S1C,D).

To investigate the ability of stVCR to restore the effects of gene interactions, we compared the partial derivatives of *Green* gene velocity with respect to *Red* gene expression with ground truth, and visualized them in spatial coordinates (Fig. 2F). Qualitatively, they were consistent, and *Green* gene inhibited *Red* gene expression mainly in the second and third groups of cells. Next, to investigate the ability of stVCR to restore the cell migration effects on gene expression, we calculated the directional derivative of *Red* gene expression for the given direction ***n*** = (1, 0) (i.e., cells moving horizontally to the right) for both learned and true dynamics (Fig. S1E). Cells at the right end moving to the right will promote gene *Red* expression, and cells at the left end moving to the right will inhibit gene *Red* expression, which overall suggests that moving horizontally outward will promote *Red* expression. Lastly, to evaluate the spatial variability of cell proliferation and regulation of gene expression on proliferation, we compared the true and learned cell proliferation rates (Fig. 2G, Fig. S1F). The results show that the first group of cells has a growth rate close to 0, the second group has a large positive growth rate, and the third group has a large negative growth rate. Additionally, we calculated the derivative of the growth rate *g* with respect to the *Green* gene, which revealed that the *Green* gene promotes apoptosis (Fig. S1G). To further validate the applicability of stVCR to the model with more complex cell migration and spatial structure, we generate another simulated data with similar gene expression but arbitrarily variable shape (Fig. S2 and Supplementary Video 2). The result shows that stVCR works consistently well in the more complicated simulation data (Fig. S2)

To further illustrate the effectiveness of stVCR over existing approaches, we next compared it with other methods designed for time-series ST data. Our results show that stVCR enables a more comprehensive range of analyses (Supplementary Table S1). Next, we conducted systematic quantitative comparison experiments with other state-of-the-art methods and ablation experiments on various modules of stVCR. Specifically, we first performed leave-one-time-point-out interpolation experiments and compared them with state-of-the-art deep learning-based trajectory inference methods [36, 42] (although they are not designed for spatial transcriptomics) and various ablation versions of stVCR (Fig. 2H and Supplementary Table S4-S5, S8-S11). The evaluation measures included the 2-Wasserstein distance of gene expression of interpolation and 2-Wasserstein distance considering both gene expression and spatial location (Methods). In addition, for the simulated data with ground truth, we also compare the migration velocity and growth rate inference (Methods) with the advanced static OT-based method [48, 50, 51] and various ablation versions of stVCR (Supplementary Video 1-2). The results show that the performance of stVCR is superior in most cases, with few sub-optimal results, and each designed module of stVCR is indispensable (Supplementary Table S2-S3, S6-S7).

Finally, we checked the scalability and robustness of the stVCR with respect to important hyperparameters and input level. We first performed a scalability analysis, which shows that stVCR is scalable for dataset size, model size, and number of observation times when the proper sample batch size is chosen (Fig. S3). Moreover, we theoretically demonstrate that the computational complexity of stVCR is linear for dataset size, model size, number of observation times and time duration, and show that the memory complexity is only linear for model size and number of observation times while independent of dataset size and time duration (Supplementary Note 2). Next, we tested the robustness of stVCR with respect to the important hyperparameters *λ*_Mch_ (Fig. S4 and Supplementary Video 3), *κ*_Exp_ (Fig. S5 and Supplementary Video 4), and *α*_Gro_ (Fig. S6 and Supplementary Video 5), where *λ*_Mch_ measures the importance of the matching loss, *κ*_Exp_ weighs the importance of gene expression and spatial coordinates in the matching term and *α*_Gro_ measures the flexibility of cell proliferation. The results show that stVCR is robust over a wide range of these parameters. We further tested stVCR’s robustness to input noise by perturbing gene expression and spatial location, and the results show stable inference performance within wide noise ranges (Fig. S7).

In summary, our benchmark tests show that (1) it is necessary to align the spatial coordinates of different time snapshots to the same coordinate system; (2) stVCR simultaneously reconstructs cell transition, migration and growth are the keys to reconstructing correct spatiotemporal dynamics; and stVCR can accurately reconstruct key regulatory mechanisms. In addition, stVCR is a scalable and robust algorithm under key hyperparameter tuning.

### stVCR reconstructs cell transition and growth dynamics of axolotl brain regeneration

To validate stVCR’s capability to learn complex continuous dynamics from spatial snapshots, we next applied stVCR for the axolotl brain regeneration dataset using single-cell Stereo-seq technology [57]. The dataset includes brain samples at 2, 5, 10, 15, 20, 30, and 60 days post-injury (DPI) to dissect both immediate wound responses and later regeneration processes. According to the original study [57], the regeneration process is mainly concentrated in 2 DPI to 20 DPI, so we took the data of 5 temporal points at 2, 5, 10, 15 and 20 and reconstructed the dynamic regeneration process using stVCR (Fig. 3, Fig. S8 and Fig. S9).

**Fig. 3:**
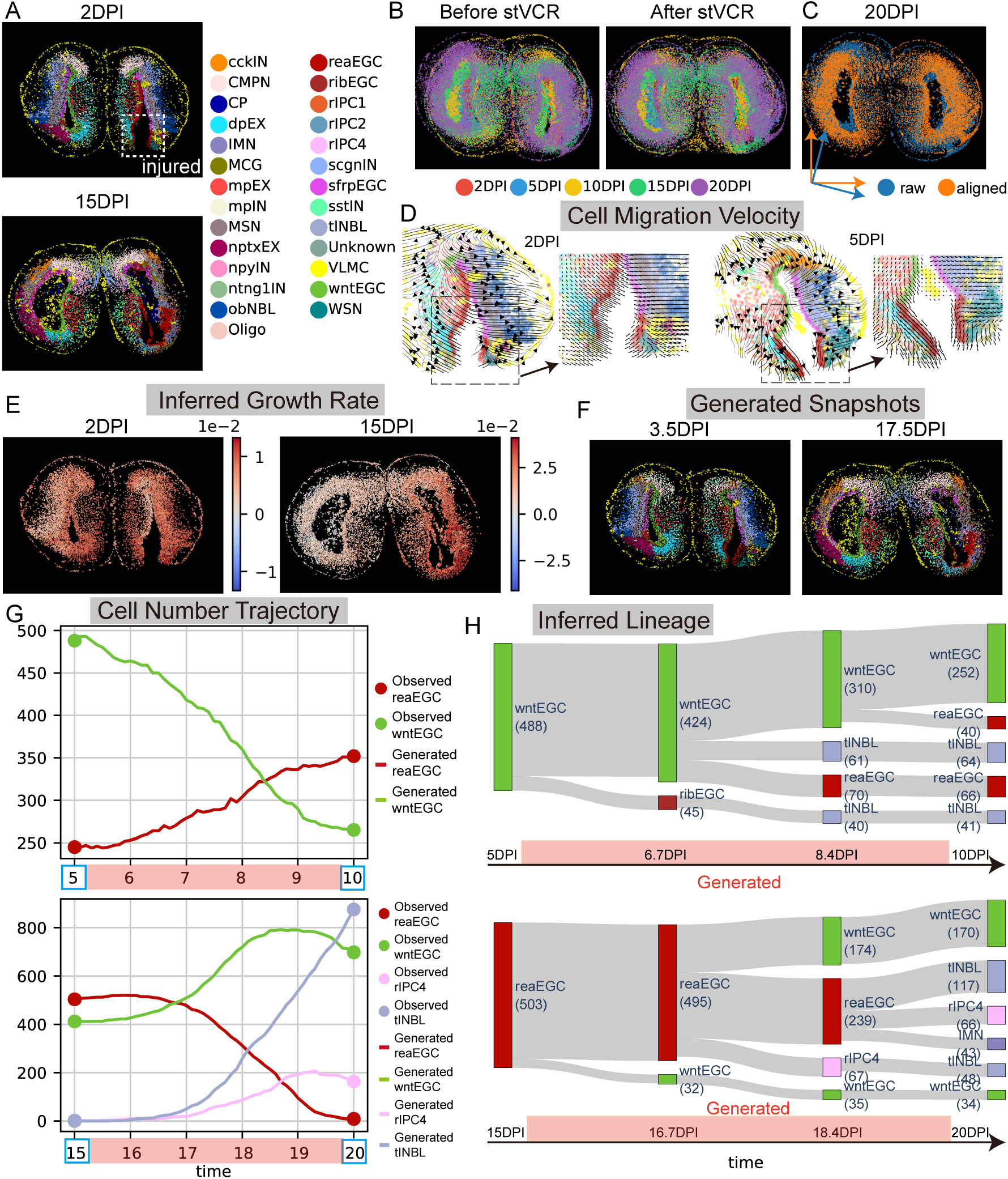
stVCR reconstructs spatiotemporal dynamics of axolotl brain regeneration. **A**. stVCR aligns the spatial coordinates of data at different time points to the same coordinate system. Top 2DPI; Bottom: 15DPI. Cell type annotations come from the original study. dpEX,dorsal palliumexcitatory neuron; IMN, immature neuron; MCG, microglial cell; MSN, medium spiny neuron; nptxEX, Nptx^+^ lateral pallium excitatory neuron; EGC, ependymoglial cell; reaEGC, reactive EGC; ribEGC, ribosomal EGC; rIPC, regeneration intermediate progenitor cell; sfrpEGC, Sfrp^+^ EGC; tlNBL, telencephalon neuroblast; VLMC, vascular leptomeningeal cell; wntEGC, Wnt^+^ EGC. **B**. Comparison of spatial coordinates of the data for all time points before and after stVCR alignment. Left: Before stVCR. Right: After stVCR. **C**. Comparison of spatial coordinates of 20 DPI data before and after alignment. **D**. stVCR inferred spatial cell migration velocity at 2DPI and 5DPI injured hemispheres. Left: Streamline plot; Right: Locally amplified grid velocity. **E**. stVCR inferred cell proliferation rate at 2DPI and 15DPI data. **F**. stVCR interpolated snapshots at 3.5DPI and 17.5DPI. Cell type annotations come from the stVCR’s time-dependent classifier based on the generated continuous gene expression values. **G**. Changes in cell number over time. Top: reaEGC and wntEGC from 5DPI to 10DPI. Bottom: reaEGC, wntEGC, rIPC4 and tlNBL from 15DPI to 20DPI. **H**. stVCR reconstructs the time-varying developmental lineages. The number in parentheses is the number of cells. Top: wntEGC from 5DPI to 10DPI; Bottom: reaEGC from 15DPI to 20DPI.

To inspect stVCR’s effect in aligning different samples, we demonstrated the aligned coordinate at different time points Fig. 3A and Fig. S8A. stVCR aligns the spatial coordinates of the data collected at different time points to the same coordinate system making them blend well (Fig. 3B). We further compared the spatial coordinates of each time point before and after the stVCR alignment (Fig. 3B and Fig. S8B), and observed that at each time point the data were adjusted to varying degrees based on the inferred rigid-body transformation, especially for the 20 DPI data (Fig. 3C). This suggests the necessity of sample alignment to infer dynamics correctly.

To further illustrate the continuous dynamics reconstructed, we trained a classifier based on existing cell annotations using a neural network, which allowed us to annotate cells at unobserved time points (Methods). stVCR recovered the gene expression, physical location, possible division and apoptosis, and possible transformation of the cell type of each cell at each moment (Supplementary Video S6). We visualized the calculated spatial velocity in coordinate space and observed cells in the vicinity of the wound migrating toward the wound when the wound was not yet fully healed, showing a response to injury, especially reactive ependymoglial cell (reaEGC) and microglial cell (MCG) (Fig. 3D).

Consistent with cell transition dynamics in response to the injury, we studied the spatial distribution of cell proliferation rates (Fig. 3E, Fig. S8D), which showed that cell proliferation rates in the injured hemispheres were significantly higher than those in the uninjured hemispheres especially near the wound site (Fig. 3E and Supplementary Table S12), implying that cell division was more active in the injured hemispheres. This phenomenon may be due to the need to compensate for cells lost due to injury. In addition, we show the interpolation results at 3.5, 7.5, 12.5 and 17.5 DPI (Fig. 3F and Fig. S8C).

To highlight stVCR’s function to generate unobserved lineage dynamics, we calculated the number of cells of each type over time based on reconstructed continuous trajectories (Fig. S9A). Interestingly, the inferred number of many types of cells does not simply vary monotonically and linearly outside the observed time point. Among the three ependymoglial cell (EGC) types, the number of reaEGC are increasing first and then decreasing, while the population of Wnt^+^ EGC (wntEGC) and Sfrp^+^ (sfrpEGC) are decreasing first and followed by increasing trend. Such a trend coincides with the original study [57] which revealed that sfrpEGC and wntEGC are transitioned into reaEGC in the earlier stage of immediate wound responses. In contrast, later reaEGC are transitioned into mature neurons (Fig. S9B). In particular, the number of wntEGC decreased while reaEGC population expanded synchronously from 5DPI to 10DPI (Fig. 3G Top).

To better visualize the lineage dynamics inferred by stVCR, we constructed the temporal developmental lineage of wntEGC from 5DPI to 10DPI, which allowed us to study the transformation of cell types at time periods other than the observed time points (Fig. 3H Top). The results showed that wntEGC were indeed partially transformed into reaEGC. Next we constructed the temporal developmental profile of reaEGC from 15DPI to 20DPI, and the results showed that it transformed into wntEGC and some neurons in intermediate and mature states (Fig. 3H Bottom), which also coincided with the trend of their cell number (Fig. 3G Bottom). In addition, we noticed a rapid increase in the number of immature neuron (IMN) and dorsal palliumexcitatory neuron (dpEX) when the number of regeneration intermediate progenitor cell (rIPC)1 and rIPC2 was sharply decreasing (Fig. S9C). Therefore, we constructed temporal developmental lineages of rIPC1 and rIPC2 from 15DPI to 20DPI, and the results showed that rIPC1 were mainly transitioned into IMN and rIPC2 were mainly transitioned into dpEX (Fig. S9D), which is also consistent with the experimental observations [57].

In summary, stVCR describes the complex dynamics of axolotl brain regeneration. In the early wound response phase of an injury, stVCR revealed that sfrpEGC and wntEGC transitioned into reaEGC, and the proliferation of cells in the injured hemisphere became active, especially EGC types in the vicinity of the wound. In addition, reaEGC moved toward the wound. As the wound gradually healed, reaEGC transformed back to wntEGC or differentiated into certain intermediate-state neurons. Eventually, neurons in the intermediate state are then transitioned into mature neurons to compensate for the loss of mature neurons due to injury.

### Gene-level mechanisms of axolotl brain regeneration revealed by stVCR

stVCR reconstructed the dynamics of axolotl brain regeneration process, in which cell migration and reaEGC proliferation play important roles. To further reveal the mechanism on the gene level, we performed stVCR analysis of these two biological processes based on the learned kinetics, and in addition, reconstructed the regulatory network between key genes (Fig. 4, Fig. S10 and Fig. S11).

**Fig. 4:**
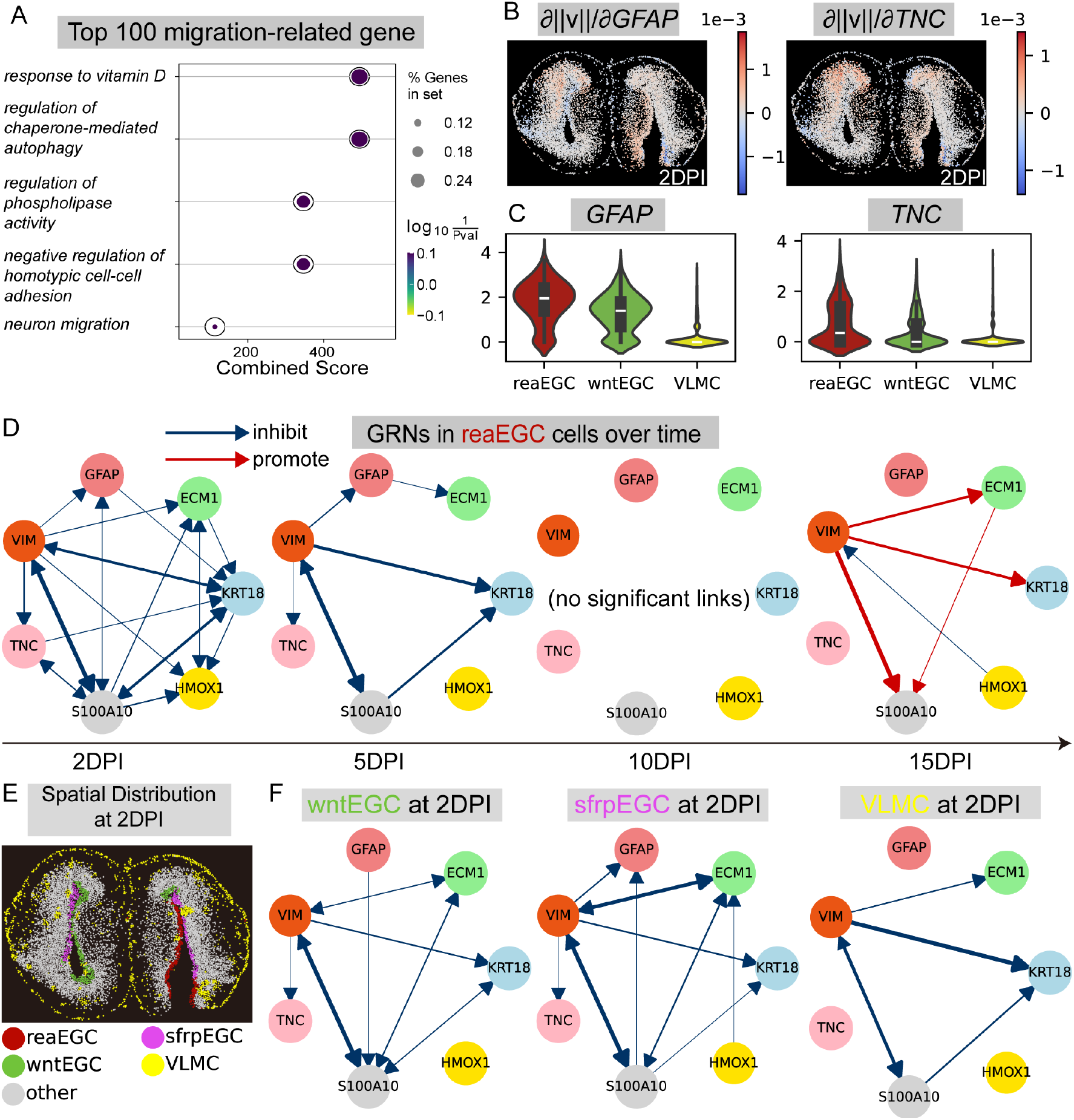
stVCR gene-level analysis of axolotl brain regeneration. **A**. GO biological process enrichment analysis of the top 100 migration-promoting genes in all cells. **B**. The partial derivative of the norm of spatial velocity ∥*v*_*z*_∥ with respect to gene expression. Two example genes in **A**, *GFAP* (Left) and *TNC* (Right). **C**. Violin plots of gene *GFAP* (Left) and *TNC* (Right) expression in reaEGC, wntEGC and VLMC cells. **D**. Gene regulatory networks in reaEGC cells for genes highly expressed in reaEGC. From left to right: 2DPI, 5DPI, 10DPI and 15DPI. **E**. Spatial distribution of reaEGC, wntEGC, sfrpEGC and VLMC cells at 2 DPI. **F**. Similar to **D**, but for sfrpEGC (Left), wntEGC (Middle) and VLMC (Right) cells at 2 DPI.

To infer gene-spatial interactions, we used stVCR to identify the top 100 migration driver genes (Methods). Through Gene Ontology (GO) biological process analysis, we identified several processes associated with cell migration (Fig. 4A), including neuron migration and negative regulation of homotypic cell-cell adhesion. Neuron migration is a crucial process for the proper positioning of neurons, while the negative regulation of homotypic cell-cell adhesion facilitates cell movement by reducing cell interactions. Interestingly, several marker genes of EGC-type cells identified in the original study, were included in stVCR’s migration driver genes, such as *GFAP, TNC, PTN, SLC1A3, GLUD1* and *ECM1*. We further visualized the migration driver gene score, i.e. *∂ ∥****v ∥****/∂q*^*j*^ of four example genes *GFAP, TNC, PTN* and *SLC1A3* (Fig. 4B and Fig. S10A), and showed that they have a promoting effect on cell migration in EGC-type cells. In addition, these genes are indeed highly expressed in EGC-type cells (Fig. 4C and Fig. S10B), especially the *GFAP, TNC* genes in reaEGC cells (Fig. 4C).

Next, we used stVCR to infer cell proliferation driver genes in reaEGC and ranked the results to obtain the top 100 growth-promoting genes (Methods). The GO analysis identified several processes closely related to cell proliferation (Fig. S10C) essential for ribosome biogenesis, protein synthesis, and the proper targeting of proteins. Similar to the migration driver gene results, the growth driver genes inferred by stVCR overlapped with several marker genes for EGCs, such as *FABP7* and *SFRP1* (Fig. S10E). We visualized the growth driver gene score *∂g/∂q*^*j*^ of these two example genes (Fig. S10D), and showed that they are significantly promoting cell proliferation and division at EGCs in the injured hemisphere.

Finally, to utilize stVCR’s function to infer dynamic gene interactions(Methods), we selected some genes (*KRT18, ECM1, GFAP, VIM, TNC, S100A10* and *HMOX1*) that were highly expressed in reaEGC (Fig. S11A), and investigated the regulatory relationship between these genes 1) at different time points and 2) in different cell types, and visualized the gene regulatory network (GRN) (Fig. 4D,E and Fig. S11B,C). We observed that in reaEGC these genes inhibit each other at an early stage (2DPI), followed by a gradual weakening of the inhibition (5DPI; 10DPI), and at a later stage (15DPI) they turn to promote each other (Fig. 4D and Fig. S11B). Thus, stVCR analysis suggests that gene regulatory relationships may be changing over time, even in the same cell type, which may be related to the discovery that reaEGC play different roles in early and late stages of injury. In addition, to investigate the gene regulatory relationships as affected by spatial distribution, we selected wntEGC and sfrpEGC, which are closer to reaEGC, and vascular leptomeningeal cell (VLMC), whereas more apart from reaEGC (Fig. 4E). We recovered the regulatory relationships between these cells at 2DPI at the previously mentioned genes (Fig. 4F and Fig. S11C). The results showed that the regulatory relationships of these genes were close in wntEGC and sfrpEGC, and closer to reaEGC, although there were some minor differences (Fig. 4F Left and Middle). In contrast, the regulatory relationship in VLMC was distinct from reaEGC (Fig. 4F Right). Thus, our results suggest that gene regulatory relationships might also be influenced by cell type and spatial location.

In summary, our gene-level mechanism of axolotl brain regeneration datasets demonstrates the ability of the stVCR to (1) find migration-driven and growth-driven genes; and (2) infer time-dependent and cell type-dependent GRN, suggesting its advantages over static OT-based methods.

### stVCR analysis of 3D Drosophila embryos and organs with optional prior

To illustrate the necessity of incorporating known biological priors for ST data with sparse temporal observations, we begin with a specially designed simulation dataset for benchmarking (Methods and Supplementary Note 4). The data consists of three types of cells named type 1, type 2, and type 3 (Fig. 5A), where type 3 cells express the *Red* and *Green* genes moderately and are at steady state. The type 1 cells will first highly express *Red* and *Green* genes, gradually decrease the expression of *green* genes, and migrate over time, transitioning to the type 2 cells that highly express only the *Red* gene. When there are sufficient observations and the time intervals are small enough, the correct result can be inferred by stVCR without any prior (Fig. S13 and Supplementary Video S9). Indeed, we can theoretically prove that the stVCR reconstructed dynamics will converge to the true dynamics when the sampling time intervals between consecutive observations converge to zero, which provides a rigorous guarantee for the algorithm (Supplementary Note 5). Meanwhile, due to the high cost of ST sequencing, the number of measurement time points is usually fewer and at longer intervals in real experiments. Thus we develop the strategy to allow users to assign known biology priors such as cell state-transition relations and structural continuity of tissues into stVCR (Methods).

**Fig. 5:**
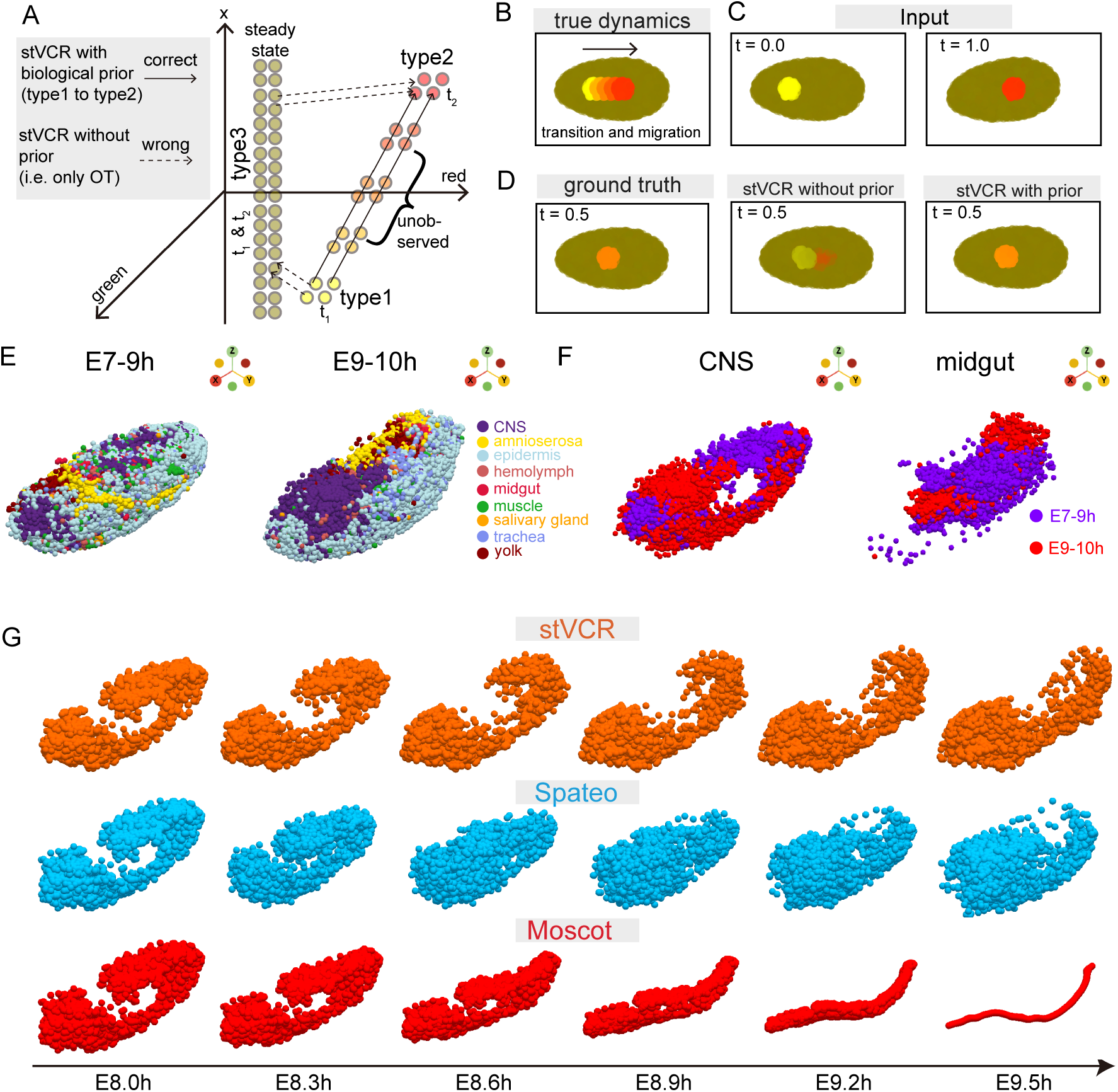
stVCR analysis of 3D Drosophila embryos and organs with biological state-transition prior and spatial structure preserving prior. **A**. Dynamic diagram of cell evolution over time in simulated data. Longer observation intervals and lack of biological knowledge guidance will result in incorrect type 3 to type 1 and type 2 transitions rather than type 1 to type 2 transitions. **B**. True dynamics of simulated data. Yellow type 1 cells transition and migrate to become red type 2 cells. Brown cells in the background are in steady state. The color was determined by the expression of three genes *Red, Green* and *Blue*. **C**. Input data to stVCR at only two time points. Left: *t* = 0.0. Right: *t* = 1.0. **D**. Interpolation results at *t* = 0.5 of stVCR with and without biological prior compared to the ground truth. Left: ground truth. Middle: stVCR without prior. Right: stVCR with prior. **E**. Spatial coordinates of 3D Drosophila embryos after stVCR alignment. Left: E7-9h. Right: E9-10h. CNS, central nervous system. **F**. Spatial coordinates of 3D Drosophila organs after stVCR alignment. Left: CNS. Right: Midgut. **G**. Comparison of spatial migration trajectories of CNS cells. From left to right: E8.0h, E8.3h, E8.6h, E8.9h, E9.2h and E9.5h. From top to bottom: stVCR, Spateo [50] and Moscot [48].

To show the effects of adding biological priors, we only input two observations at *t* = 0 and *t* = 1 (Fig. 5A,C) to stVCR. Without prior knowledge, stVCR would infer the wrong type 1 to type 3 and type 3 to type 2 transitions rather than the correct type 1 to type 2 transition (Fig. 5, Supplementary Video S7-S8). In comparison, the interpolation results show that stVCR with a type 1 to type 2 state-transition prior is closer to the ground truth at unobserved time points (Fig. 5D and Fig. S12). Overall, the above experimental results illustrate the benefit of adding the correct biological prior into datasets with fewer observations and longer intervals for more accurate results.

In order to validate the effectiveness of the strategy of combining biological prior and spatial structure preserving prior on real datasets, we next applied stVCR priors for the 3D Drosophila embryos and organs dataset using single-cell Stereo-seq technology [50]. This datasets include only two time points E7-9h and E9-10h. We set the former moment (E7-9h) in the 3D Drosophila embryo data to *t* = 8h and the latter moment (E9-10h) to *t* = 9.5h. The data contains 9 tissues (Fig 5E), and we added biological priors central nervous system (CNS) transition to CNS, midgut transition to midgut, and amnioserosa transition to amnioserosa. Additionally, we added the spatial structure preserving priors for CNS and midgut. We aligned the two observations of data and reconstructed the dynamics between the two moments using the stVCR with the above priors (Supplementary Video S10). Fig. 5E shows the aligned 3D Drosophila embryo. We focused on the CNS and midgut (Fig. 5F). We observed that the anterior of the CNS of Drosophila at the latter moment overlapped with the posterior of the CNS at the former moment (Fig. 5F Left), and the midgut consisted of two parts at the former moment and only one part at the latter moment (Fig. 5F Right).

To benchmark stVCR with other static OT-based methods and highlight its unique function to model continuous dynamics, we compared the spatial migration dynamic trajectories of CNS reconstructed by stVCR with Spateo [50] and Moscot [48] (Supplementary Video S11-S13 and Fig. 5G). Since both Spateo and Moscot are based on static OT and do not directly reconstruct the intermediate process, we obtained the intermediate process by linear interpolation based on the inferred static optimal map. In the spatial trajectory reconstructed by stVCR, the cells in the posterior of CNS gradually migrated to the anterior along the internal structure of the CNS (Supplementary Video S11 and Fig. 5G Top row). In Spateo, the cells at the posterior of the CNS were disconnected from the main body and then migrated to the anterior to merge into one part (Supplementary Video S12 and Fig. 5G Middle row). One possible explanation is that Spateo is based on static OT and does not constrain the consistency of the intermediate trajectory. In Moscot, cells migrate and aggregate to a few locations (Supplementary Video S13 and Fig. 5G Bottom row). We speculate the possible reason is that Moscot tackles the unbalanced OT problem by adding the KL divergence penalty, so that the cells at the first moment correspond to a few cells at the second moment. In addition, we reconstructed the spatial trajectory of midgut using stVCR and compared it with Spateo and Moscot (Supplementary Video S14-S16 and Fig. S14). The results showed that both stVCR and Spateo observed that two parts of the midgut at the first moment merged into one part at the later moment (Supplementary Video 14-15 and Fig. S13C First two row). In Moscot, the spatial trajectories of midgut migrated and aggregated to a small number of locations similarly to the CNS results.

In summary, we demonstrate the theoretical convergence of stVCR in large sample cases through mathematical derivations. We also highlight the benefit of adding known state-transition priors and spatial structure-preserving priors in case of limited observations through computations on simulated data. The application in 3D Drosophila datasets indicates the superiority of stVCR compared to existing methods based on static OT.

## Discussion

Time-series spatial transcriptomics data has made it possible to reconstruct the entire spatiotemporal dynamic process of cell fate determination. To dynamically connect unpaired snapshots and align temporal slices from various coordinate systems, we present stVCR to (1) simultaneously reconstruct and continuously generate cell differentiation, migration in physical space as well as division and apoptosis; (2) align spatial coordinates from data collected at different time points and (3) investigate the complex interactions between cell phenotype transitions, spatial migration, and proliferation.

The GW-OT framework provides an effective approach for modeling heterogeneous space data. In analyzing time-series spatial transcriptomics data, where spatial coordinates exhibit heterogeneity across observation time points [52], existing methods (e.g., Moscot [48], Spateo [50], and DeST-OT [51]) employ GW-OT for data modeling. Specifically, Spateo [50], and DeST-OT [51] further achieve spatial coordinate alignment between adjacent time points by solving the weighted Procrustes problem [58] based on the static OT mapping, thereby obtaining cell migration velocity. However, these methods remain within the static optimal transport framework, and could have limitations to nonlinearly generate the samples at unseen temporal points. Compared to existing methods that reconstruct trajectories from time-series spatial transcriptomics data [48, 50, 51], stVCR employs rigid body transformation invariant OT and particle evolution dynamics for unbalanced distribution matching, making it a unique method to use dynamic OT modeling for time-series spatial transcriptomics data. Additionally, stVCR can also model optional known biological priors and spatial structure-preserving priors in the dynamical framework. Once the entire dynamic process is reconstructed, stVCR can further perform a series of downstream analyses that are not feasible with static OT.

While this study provides valuable insights, there are certain aspects that could benefit from further exploration in future research. Firstly, stVCR model requires numerical integration of neural network-represented ODEs, which entails substantial computational complexity. Notably, recent studies have proposed simulation-free methods for reconstructing single-cell continuous dynamics [35, 36], which may demonstrate generalizability in time-series spatial transcriptomic data. Secondly, the accuracy of reconstruction based on 2D slices critically depends on slice localization precision, whereas 3D spatial transcriptomic techniques inherently circumvent this limitation. With remarkable advancements in 3D spatial transcriptomic experimental protocols [59] and reconstruction algorithms [50, 60], this technical challenge is anticipated to be resolved in the near future. Due to the limitations of 3D time-series spatial transcriptome data availability, further applications and performance evaluations on more challenging datasets are needed. Thirdly, the current stVCR model does not incorporate intercellular interaction, which could be improved by incorporting studies on cell-cell communication [61–65] and biomechanical regulation in developmental biology [66, 67]. Fourthly, learning both the low-dimensional representation of gene expression and the dynamics in the low-dimensions simultaneously instead of separate learning strategy could yield better results [18]. Finally, the integration of multimodal data (such as lineage tracing [68] and multi-omics measurements [69]) and the incorporation of intrinsic cellular dynamics and noise (encompassing transcriptional regulation, RNA splicing, and degradation processes) are crucial directions for enhancing model predictive capabilities.

Overall, stVCR provides a unified and robust method for dynamical generative modeling of time-series spatial transcriptomics data, which reconstructs the entire spatiotemporal processes of single cells from a few given snapshots and investigates the complex space-gene regulatory mechanisms. Further exploration along this line while overcoming its limitations will be a promising topic in the future.

## Methods

### Overview of optimal transport formulation

In stVCR, we utilize a newly modified dynamical optimal transport (OT) formulation to reconstruct the spatiotemporal dynamics of single cells for snapshot spatial transcriptomics data. For basic OT setup, let *α* and *β* be two probability distributions with probability mass ***a*** = (*a*_*i*_) and ***b*** = (*b*_*j*_) concentrated at *{****x***_*i*_*}*_*i*=1:*n*_ and *{****y***_*j*_*}*_*j*=1:*m*_, respectively (i.e., 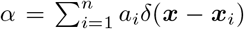 and 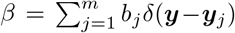, where *δ*(*·*) stands for the Dirac’s *δ*-function). Here the notation 1:*n* is a shorthand for the indices *{*1, …, *n}*, and the same rule applies to similar notations in other places. In the well-known Kantorovich’s OT formulation, one aims to find the optimal transportation plan *P* = (*p*_*ij*_) such that the transportation cost

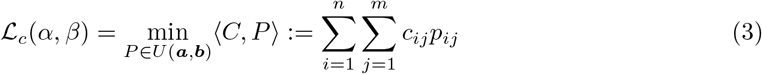

to be optimized, where 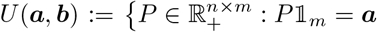 and *P* ^T^𝟙*n* = ***b***} and *C* = (*c*_*ij*_) is the cost matrix. When *c*_*ij*_ = *c*(***x***_*i*_, ***y***_*j*_) with 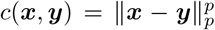, where ∥***x***∥_*p*_ is the vector *ℓ*^*p*^ norm, the *p*-Wasserstein distance is defined as *W*_*p*_(*α, β*) = (*ℒ*_*c*_(*α, β*))^1*/p*^. The optimal coupling matrix component *p*_*ij*_ characterizes the probability that ***x***_*i*_ will be transported to ***y***_*j*_ [70].

A special case is *p* = 2, i.e.,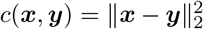. In this case, the above OT formulation has an equivalent dynamic form (Benamou-Brenier form [33]) by minimizing the transport kinetic energy

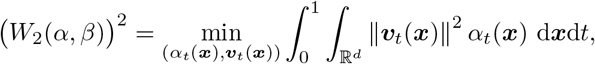

where *α*_*t*_, ***v***_*t*_ satisfies the continuity partial differential equation (PDE)

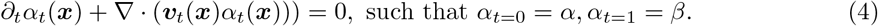

The vector field *{* ***v***_*t*_(***x***) *}* _*t∈*[0,1]_ is to be optimized such that the boundary conditions are satisfied and the minimal kinetic energy is achieved.

The dynamic formulation can be generalized to the case when the total mass of *α* and *β* are not equal (unbalanced setting). A common approach is to consider the so-called Wasserstein-Fisher-Rao (WFR) distance [43, 71, 72]

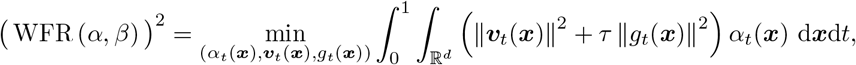

where *α*_*t*_, ***v***_*t*_, *g*_*t*_ satisfies the PDE

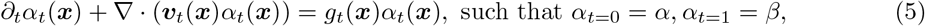

where *g* describes the rate of mass change, corresponding to the rate of cell proliferation. In the context of spatiotemporal transcriptomics, the state variable ***x*** will be extended to (***x, q***) which includes both the spatial and gene expression variables, and special treatments will be introduced for spatiotemporal matching as described below.

### Notation conventions in stVCR

To better describe the stVCR algorithm, we use the notation 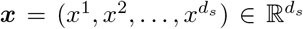, 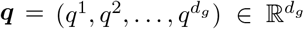 for spatial coordinates and gene expressions for each cell, respectively, where *d*_*s*_ is the dimension of the spatial coordinates (usually 2 or 3), and *d*_*g*_ is the dimension of the embedded gene expression space (*d*_*g*_ = 10 in our setup). For the considered spatiotemporal transcriptome data, we assume there are *n*_*k*_ cells at time *t* = *t*_*k*_ for *k* = 0, 1, …, *K*. We denote the available datasets by

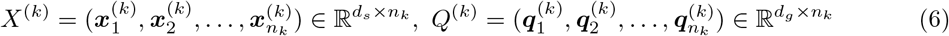

at *t* = *t*_*k*_, where the subscript *i* in 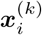and 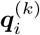 indicates the cell label. In the data analysis, we often need to consider the spatial coordinates after alignment with rigid body transformations at time points *t* = *t*_1_, …, *t*_*K*_ (we set *t* = *t*_0_ as the reference state), which we denote by

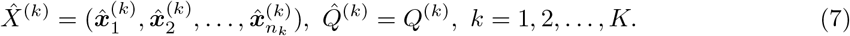

When only rotation *R* and translation ***r*** are considered, 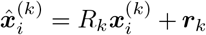. The data (empirical) distribution formed by the dataset (*X*^(*k*)^, *Q*^(*k*)^) is denoted by

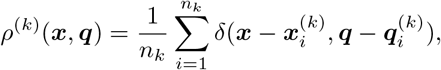

where *δ*(*·*) stands for the Dirac’s *δ*-function in (***x, q***)-space. Correspondingly, we can define 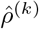 for the dataset 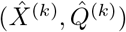 after alignment. Both *ρ*^(*k*)^ and 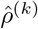 are probability distributions.

We use the notation *f*_*t*_(***x, q***) or the shorthand *f*_*t*_ to denote the time dependence of a function *f* (***x, q***, *t*), while for time dependent coordinates (***x***(*t*), ***q***(*t*)), we use the notation (***x***^(*t*)^, ***q***^(*t*)^) to be consistent with the notation (*X*^(*k*)^, *Q*^(*k*)^). All of the vectors are bold-faced, while the matrices are in normal fonts.

### Spatial alignment for temporal snapshots

Due to the unknown deformation of the tissue during spatial transcriptome (ST) sequencing, we need to align the spatial coordinates of the ST data at different time points before the subsequent analysis. We assume the large scale deformation of the observed coordinates contains only rigid body transformations (i.e., rotations and translations) in this work. We do not follow the Gromov-Wasserstein (GW) optimal transport framework for its over-generality and large computational cost. Yet, we adopt the optimal transport approach in [73] by explicitly modeling the set of rotation and translation manipulations *𝒢* = *{* (*R*, ***r***) *}*, and simultaneously finding the optimal matching of distributions and the optimal transformation through the optimization problem:

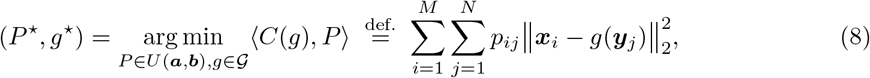

where *g*(***y***_*j*_) := *R****y***_*j*_ + ***r*** for *g* = (*R*, ***r***). It can be solved by an alternating iteration

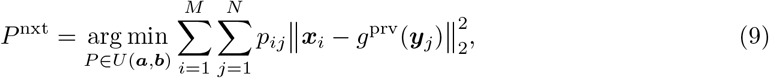

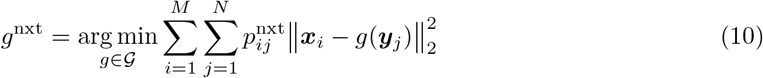

for *g* and *P* from the previous step to next step. The subproblem (9) is to solve a static OT, while the subproblem (10) is a weighted Procrustes problem [58] for solving the optimal rotation matrix *R* and translation vector ***r***.

We call this problem *rigid body transformation invariant optimal transport*, and it explicitly models unknown rigid body transformation, making it possible to compute ground cost functions between distributions at different time points. It can be easily modified to model more general cases, e.g., if affine transformations are allowed in the tissue measuring process. Note that the optimal rigid body transformation is obtained through the simulatneous optimization of the dynamical variables.

### Dynamics loss in stVCR

We are concerned with the entire dynamics of cell population evolution and therefore use dynamical optimal transport. Since the number of cell population is constantly changing due to cell division and apoptosis during evolution, unbalanced setting is necessary.

stVCR reconstructs the dynamical velocity (***v***_*t*_, ***p***_*t*_) (corresponding to the coordinate (***x***_*t*_, ***q***_*t*_)) and the distributions *ρ*_*t*_(***x, q***) by interpolating the input population densities *ρ*^(*k*)^ up to a normalization at *t* = *t*_*k*_ using a transport-with-growth PDE

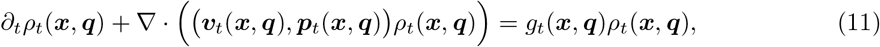

where ***v***_*t*_(***x, q***) refers to the spatial velocity, ***p***_*t*_(***x, q***) refers to the RNA velocity of cells, and *g*_*t*_(***x, q***) refers to the proliferation rate of cells. Eq. (11) is a natural extension of (5) in the spatiotemporal transcriptomics setup.

We take the coordinate system at *t*_0_ as the reference, and assume that the coordinate system at *t*_*k*_ (*k* = 1, 2, …, *K*) differs from the coordinate system at *t*_0_ by a rotation *R*_*k*_ and translation ***r***_*k*_. Borrowing the idea of Wasserstein-Fisher-Rao (WFR) distance for unbalanced optimal transport [42], we obtain the optimal dynamics (*ρ*_*t*_, ***v***_*t*_, ***p***_*t*_, *g*_*t*_) and the optimal rigid body transformation (*R*_*k*_, ***r***_*k*_) for *k* = 1, 2, …, *K* by minimizing the kinetic energy of spatial migration and gene expression change and growth energy

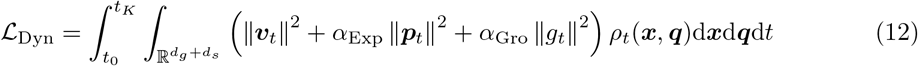

for (*ρ*_*t*_, ***v***_*t*_, ***p***_*t*_, *g*_*t*_; *R*_1:*K*_, ***r***_1:*K*_) such that the obtained solution *ρ*_*t*_ optimally matches 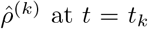, where 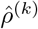 refers to the distribution formed by 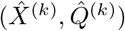 after alignment through rigid body transformation (*R*_*k*_, ***r***_*k*_). See details of the distribution matching in next subsection.

Direct derivation of the solution of the Feynman-Kac type PDE (11) by characteristics [42, 74] shows the above loss function can be rewritten as the following dynamics loss

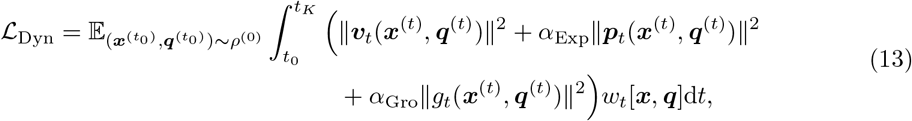

where ***x***^(*t*)^, ***q***^(*t*)^ satisfies the characteristic ordinary differential equations (ODEs)

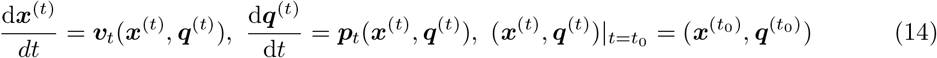

and the weights 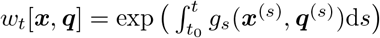 satisfies the ODE

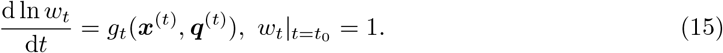

Under this setup, the density *ρ*_*t*_ has the representation 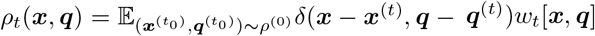 with the total probability mass 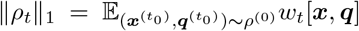, where ∥*ρ*_*t*_∥_1_ := ∫*ρ*_*t*_(***x, q***)d***x***d***q***. The three parts in *ℒ*_Dyn_ correspond to the kinetic energy of spatial migration, *ℒ*_Spa_, kinetic energy of gene expression change, *ℒ*_Exp_, and growth energy, *ℒ*_Gro_, respectively, considered in Eq. (2) or (12).

The formulation (13) is suitable for the numerical evaluation of the loss function through Monte Carlo particle simulations instead of density estimation in high dimensional space. We also remark that the evolution of (11) can be implemented in the forward (from *t*_0_ to *t*_*K*_) or backward (from *t*_*K*_ to *t*_0_) way, and similar formulations as above can be obtained correspondingly. Both directions are taken in our computations for a more robust performance.

### Matching loss in stVCR

There is a challenge in the distribution matching between *ρ*_*t*_ and 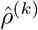 since *ρ*_*t*_ is not a normalized probability distribution due to the proliferation. To deal with this issue, we impose the matching condition ∥*ρ*_*t*_ ∥_1_ = *n*_*k*_*/n*_0_ and 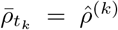 where 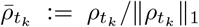 is the normalized version of. These conditions are indeed realized as soft penalties to perform distribution matching. This matching between the total mass, distributions, and the determination of *{* (*R*_1:*K*_, ***r***_1:*K*_) *}* are obtained simultaneously in terms of the rigid body transformation invariant OT.

Define the weights *w*_*t,j*_ = *w*_*t*_[***x***_*j*_, ***q***_*j*_] for the cell *j* with initial state 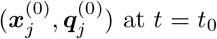. With this notation, we have the evolved distribution

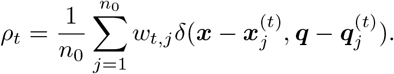

The total mass of *ρ*_*t*_ is 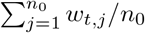, which no longer corresponds to a probability distribution. This non-normalization is due to the cell proliferation.

The matchings between the total mass, and the normalized distributions 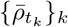 and 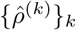 is performed through the loss function

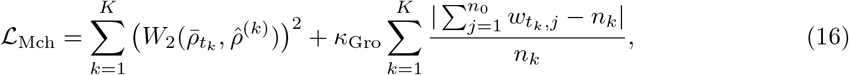

where the first term penalizes the normalized distribution mismatch through the square of the 2-Wasserstein distance between 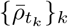 and 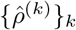, which promotes the gene expression and spatial coordinates of learned dynamics close to the observed data. The second term penalizes the total mass mismatch through their relative error, which promotes the matching of cell numbers with observed data. Naturally, we take the cost matrix *C*^(*k*)^ by integrating gene expression and spatial coordinates with components

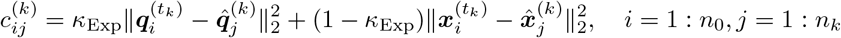

where 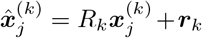 is the coordinate after the alignment manipulation, *κ*_Exp_ is an adjustable hyperparameter that measures the importance of gene expression or spatial coordinates.

### Modeling known cell type transition prior

Modeling known biological prior can help to infer the spatiotemporal dynamics of single cells more accurately, especially for situations where there are few observed time points or long time intervals between adjacent time points. For known permissible cell type transitions, we grouped cells with permissible transitions at different time points into the same type. We use 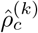 to refer to the empirical distribution of type *c* cells in the observed data (after rotation and translation) at the time point *t*_*k*_, and 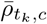 to refer to the normalized distribution of type *c* cells evolving from *t*_0_ to *t*_*k*_. Assuming a total of *C* types, we realize the distribution matching for each known permissible cell type transitions, i.e.,

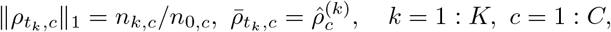

where *n*_0,*c*_ and *n*_*k,c*_ are the number of type *c* cells at time *t*_0_ and *t*_*k*_, respectively. In this circumstance, the matching loss (16) has to be revised as

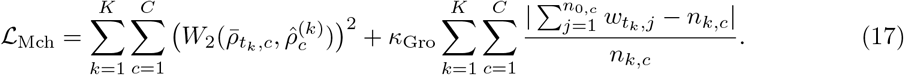

### Modelling spatial structure preserving prior

Organ development obeys physical rules, and its spatial structure cannot change at will. For example, some organs remain connected as they develop without breaking into multiple parts. However, when the time interval between observations is long, usual OT-methods often produce results that violate the physical rules in order to minimize the energy, so we need to explicitly model this prior to maintain the spatial structure of the specified organ. For the organs that need to be spatially structure-preserved, we first construct a graph according to spatial coordinates and gene expression. More specifically, at *t* = *t*_0_, we first construct a *k*_spa_ nearest-neighbor graph using the spatial coordinates, and then find the *k*_nbr_ cells with the closest gene expression as the final neighbors from the *k*_spa_ spatial neighbors of each cell in the specified structure to complete the graph construction. We denote the index set of these *k*_nbr_ neighbor cells, which is a subset of {1 : *n*_0_}, by 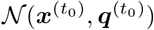 for each cell 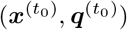 in the specified structure at *t* = *t*_0_. In order to keep the specified organ development obeying the physical rules, we add the optional spatial structure preserving loss function

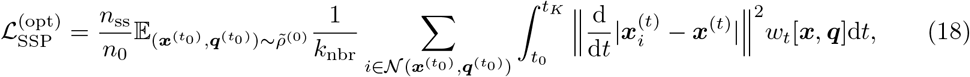

where 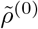 is the probability distribution for the cells in the specified structure and *n*_ss_ is the is the number of cells in the specified structure. 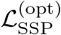 preservers spatial structure by promoting as little change as possible in the distance between spatial trajectories of neighboring cells. It can be understood as a continuous time limit of the Gromov-Wasserstein OT distance.

### Deep learning-based solver in stVCR

Optimizing the total loss *ℒ* in (1) is generally difficult, we use deep learning to find an approximate solution. For the arguments (*ρ*_*t*_, ***v***_*t*_, ***p***_*t*_, *g*_*t*_; *R*_1:*K*_, ***r***_1:*K*_) in the optimization, *ρ*_*t*_ is indeed determined by (***v***_*t*_, ***p***_*t*_, *g*_*t*_). We suppose that changes in gene expression, cell migration, and cell proliferation depend on current gene expression and spatial location, and are parameterized by neural networks, i.e., ***v***_*t*_(***x, q***) = ***v***(***x, q***, *t*; *θ*_***v***_), ***p***_*t*_(***x, q***) = ***p***(***x, q***, *t*; *θ*_***p***_) and *g*_*t*_(***x, q***) = *g*(***x, q***, *t*; *θ*_*g*_), where *θ* := (*θ*_***v***_, *θ*_***p***_, *θ*_*g*_) are the parameters of the neural networks. For the rigid body transformations, the rotation matrix *R* can be explicitly parameterized. In 2D case, we take

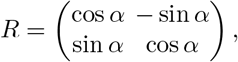

where *α* is the rotation angle. In 3D case, we parameterize the rotation matrix *R* by the Euler angles *α, β*, and *γ* with *R* = *R*_*x*_(*α*)*R*_*y*_(*β*)*R*_*z*_(*γ*), where

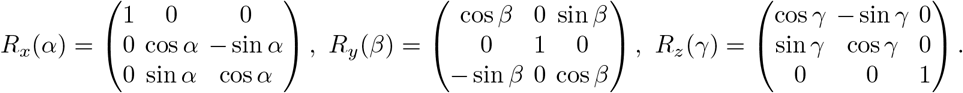

Therefore, the overall parameters we need to optimize are the neural network parameters *θ*, as well as the rotation angles *α*_*k*_ (or Euler angles *α*_*k*_, *β*_*k*_ and *γ*_*k*_ in 3D case) and translation vectors ***r***_*k*_ for *k* = 1 : *K*.

With the above parameterization, the constraint PDE *∂*_*t*_*ρ*_*t*_ + *∇ ·* ((***v***_*t*_, ***p***_*t*_)*ρ*_*t*_) = *g*_*t*_*ρ*_*t*_ with initial value 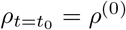 can be solved by the particle approximations through the ODEs (14) and (15) by replacing the functions (***v***_*t*_, ***p***_*t*_, *g*_*t*_) with (***v***(***x, q***, *t*; *θ*_***v***_), ***p***(***x, q***, *t*; *θ*_***p***_), *g*(***x, q***, *t*; *θ*_*g*_)). The evaluation of the integral in (13) can be also performed by numerical quadrature in time with the parameterized (***v, p***, *g*). Finally, the overall loss

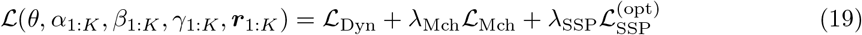

can be evaluated through the deep learning approximations. The optimization of the parameters are achieved by the Adam optimizer [75]. In (19), *ℒ*_Dyn_ and *ℒ*_Mch_ are required and 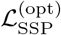 is optional. Whether or not to include cell type transition prior in *ℒ*_Mch_ is also optional.

### Parameter initialization and training details

The structure of our neural networks ***v***(***x, q***, *t*; *θ*_***v***_), ***p***(***x, q***, *t*; *θ*_***p***_) and *g*(***x, q***, *t*; *θ*_*g*_) use multilayer perceptron (MLP) with 128 neurons per layer for a total of 6 layers. To obtain an initial rotation and translation, We downsample 5000 cells in the data, and then use static rigid-body transformation invariant OT on the downsampled data. The training process involves solving the ODEs represented by the neural network, that is neural ODE, which we implement using the torchdiffeq package [56]. It also involves computing the static OT distance, which we implement using the POT package [55]. Finally, in order to enhance the matching with the observed data at each time point and improve the robustness of the algorithm (the new organs or cell types may appear in later time points), we not only compute the loss function defined by (19) by sampling the data from *t*_0_ to later time points, but also compute a similar loss function by sampling the data from other time points *t*_*k*_, evolving forward and backward. In actual computations, we choose to sample from the first time point *t*_0_ and the last time point *t*_*K*_ to balance the accuracy and computational overhead. We provide a detailed pseudocode description of the stVCR training process in the Supplementary Note 6.

### Data preprocessing

For gene expression count matrix, we first normalized the raw counts data using size factor. Then we selected the top 2000 highly variable genes. Finally, we utilize an Autocoder to project highly variable genes to low dimensions. Specifically, we represent an encoder ***q***_emb_ = *f*_enc_(***q***_ori_, *θ*_enc_) and a decoder 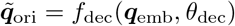 using neural networks, where the input of the encoder is the original gene expression ***q***_ori_ and the output is the low-dimensional embedding ***q***_emb_, and the decoder is the opposite. The loss function is taken as

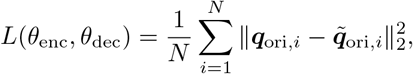

where *N* refers to the total number of cells. In actual computations, we take the dimension of the low-dimensional embedding ***q***_emb_ to be 10. Also for simplicity of notation, we still use ***q*** to refer to ***q***_emb_ to denote the low-dimensional embedding of gene expression unless otherwise stated.

### Quantitative metric used in benchmark experiments

For simulation data with ground truth, we use the mean absolute error (MAE) of the inferred growth rate and ground truth

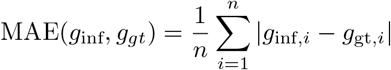

and the mean root mean square error (MRMSE) of the inferred spatial velocity and ground truth

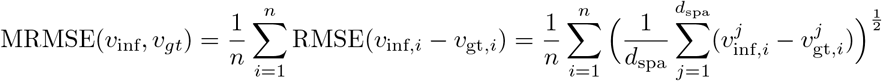

as quantitative metrics, which correspond to Supplementary Tables S2-S3 and S6-S7.

Additionally, we conducted leave-one-time-out experiments. For all methods, we used the 2-Wasserstein distance between the interpolated gene expression and the ground truth as an evaluation metric, which correspond to Supplementary Tables S4, S8 and S10. For methods capable of interpolating both spatial locations and gene expression (only stVCR and some of its ablated versions), we employed the 2-Wasserstein distance that jointly considers both spatial locations and gene expression, which correspond to Supplementary Tables S5, S9 and S11. Specifically, for simulated data with ground truth spatial locations, we computed the cost matrix using the actual spatial locations

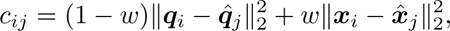

where *w* is a weight coefficient and is taken as 0.9 in the actual calculation. For biological data, we computed the cost matrix using the spatial locations aligned via rigid transformation (Supplementary Note 1).

### Time-dependent cell type classifier

After we recovered the entire cell dynamics, we could obtain the gene expression and spatial location of cells at unobserved moments. In order to obtain type annotations for these cells at unobserved moments, we train a time-dependent cell type classifier using a neural network in cells that already have cell type annotations, so that we can use it for cell type annotation at unobserved moments. Specifically, we represent a classifier by a neural network ***f*** = ***f***_type_(***x, q***, *t*; *θ*_type_). The inputs are gene expression ***q***, spatial coordinates ***x***, and time *t*, and the outputs are probability distributions indicating the probability with which the cell (***x, q***) at time *t* belongs to each cell type. The loss function is taken as

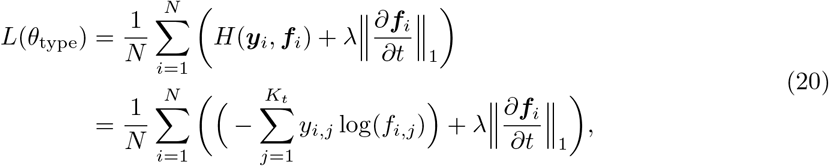

where *H*(*·, ·*) is the cross entropy, ***f***_*i*_ := ***f*** (***x***_*i*_, ***q***_*i*_, *t*), ***y***_*i*_ is the annotated cell type for cell *i, K*_*t*_ is the total number of cell types, and *y*_*i,j*_, *f*_*i,j*_ is the annotated and trained probabilities that the cell *i* is of type *j*. The first term in (20) is to force the trained classifier to be consistent with the known cell type annotations, while the second term is a regularization to promote the smoothness of the neural network classifier in time through an ^1^ norm of the time derivative on observation points.

### Downstream analysis

For notational simplicity, we omit the notation for the parameters of the neural networks in the following description. When we use the stVCR to obtain the spatial velocity ***v***(***x, q***_emb_, *t*), the gene expression velocity ***p***_emb_(***x, q***_emb_, *t*) in the embedded space, the cell proliferation rate *g*(***x, q***_emb_, *t*), and the time-dependent classifier *f*_type_(***x, q***_emb_, *t*), we can perform a series of downstream analyses, including interpolation, prediction, and study of cell-specific gene-gene, gene-space and space-space interaction and the effects of gene and spatial migration on cell proliferation, etc. Below we give a detailed description of some downstream analysis tasks.

#### Recovery of cell evolution rates in original space

As stated in the data preprocessing step, the original gene expression and its dimension-reduced expression value in embedded space, ***q***_ori_ and ***q***_emb_, are related by the trained encoder-decoder neural network *f*_enc_, *f*_dec_ as ***q***_emb_ = *f*_enc_(***q***_ori_) and ***q***_ori_ = *f*_dec_(***q***_emb_). With this representation, we can easily recover the spatial velocity, gene expression velocity and proliferation rate in original gene expression space:

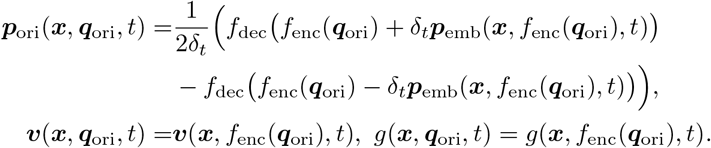

#### Interpolation and prediction

For some interpolation time *t*_int_ which the user is interested in, we choose the observation time *t*_obs_ that is closest to *t*_int_. Without losing generality, we assume that *t*_obs_ *< t*_int_. We take the observed data ***q***_*i*_ and ***x***_*i*_ at *t*_obs_ as the initial values, and then evolve according to the dynamics learned by stVCR to obtain the interpolation result. More specifically, for the *i*_th_ cell to evolve from time *t* to *t* + *δt*, we first compute

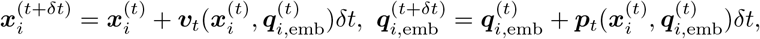

then generate a random number *U ∼* Uniform[0, 1]: if 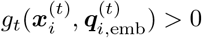 and 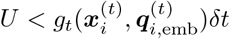, perform cell division; if 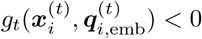 and 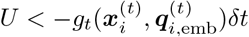, then perform cell apoptosis. The prediction task is completely similar to the interpolation task, which only needs to take the data of the last time point as the initial value.

#### Cell specific gene-gene, gene-space and space-space interaction

We can study the interaction between genes and space from the learned quantities. For a cell *i*, as well as the target gene *j* and the source gene *k* of interest, we can calculate 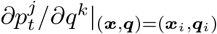, which represents how increased expression of gene *k* changes the velocity of gene *j* in cell *i*. If it is positive, it means that gene *k* promotes gene *j* in cell *i*, otherwise gene *k* inhibits gene *j*.

Similarly, for cell *i*, and axes *j* and *k* of interest, we can calculate 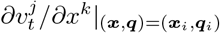 to study space-space interaction. This is used in Spateo [50], although their spatial vector fields ***v***(***x***) are independent of gene expression ***q*** and time *t*.

In stVCR, gene expression ***q*** and spatial coordinates ***x*** interact, which means we can study how cell migration affects gene expression. For cell *i*, a target gene *j*, and a given unit direction ***n*** = (*n*^1^, *n*^2^, *n*^3^) (or ***n*** = (*n*^1^, *n*^2^) for 2D case), we can calculate the directional derivative

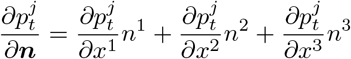

at (***x, q***) = (***x***_*i*_, ***q***_*i*_), which describes how the migration of cell *i* to the given direction ***n*** affects the expression of gene *j*, with positive values representing promotion and negative values the opposite. In addition, we can study how gene expression affects cell migration. For cell *i* and gene *j* of interest, we can define

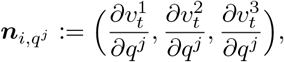

which describes that increased expression of gene *j* will promote the cell *i* migration in the direction 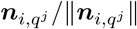 and 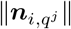 indicates the promotion intensity.

Finally, we can define partial derivative of the norm of cell migration velocity with respect to gene *∂∥****v***_*i*_*1∥∂q*^*j*^ of cell *i* and gene *j* as migration driver gene score based on the above calculations

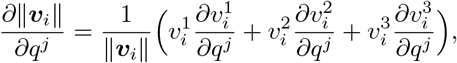

where ***v***_*i*_ := ***v***_*t*_(***x***_***i***_, ***q***_***i***_).

#### Cell specific effects of gene and spatial migration on growth

For cell *i* and gene *j* of interest, we can calculate 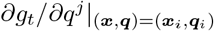, which describes the effect of gene *j* on cell proliferation, where a positive value means promoting, and a negative value the opposite. This concept is used in TIGON [42]. However, their growth function *g*_*t*_(***q***) does not depend on spatial coordinates ***x***, so the effect of cell spatial migration on growth cannot be studied.

In stVCR, cell proliferation depends on both its gene expression and its spatial location, so we can also study how cell migration affects its growth. For cell *i* and a given unit direction ***n*** = (*n*^1^, *n*^2^, *n*^3^), we can calculate the directional derivative

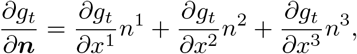

which describes how cell migration in the direction ***n*** affects its proliferation.

Finally, we can define partial derivatives of the cell proliferation rate with respect to gene *j* for cell *i*, 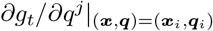, as growth driver gene score based on the above calculations.

#### Temporal Developmental lineage construction

Since we can interpolate for any time points of interest and can annotate cells at these unobserved time points with the time-dependent classifier ***f***_type_(***x, q***_emb_, *t*), we can construct temporal developmental lineages of cells of interest.

### Simulated data setup

In this paper, three simulation data are included. The first corresponding to Fig. 2, the second corresponding to Fig. S2, and the third corresponding to Fig. 5 and Fig. S13. Below, we will introduce how these three simulation data are generated.

For the first simulation data corresponding to Fig. 2, the dynamics consists of three genes *Red, Green* and *Blue* and two spatial coordinates *x* and *y*, whose regulation is shown in Fig. 2A. Such a regulatory relationship can be described by a system of stochastic differential equations for gene expression and spatial coordinates

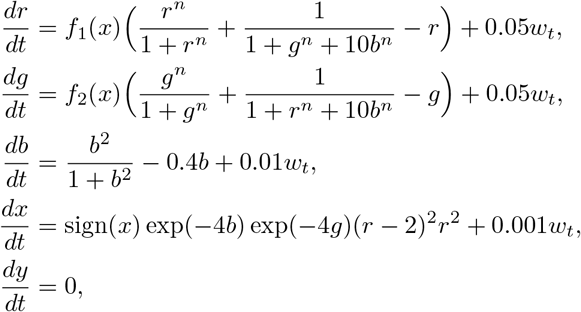

where *r, g* and *b* refer to gene *Red, Green* and *Blue, f*_1_(*x*) and *f*_2_(*x*) refer to the factors that depend on the coordinates *x*, and *w*_*t*_ is a standard Brownian motion. In the computation, we take *n* = 4 to simulate the nonlinear regulation between genes. If we ignore these two *x*-related factors *f*_1_(*x*) and *f*_2_(*x*), *r* and *g* are a toggle switch of equal status. We hope |*x*| *>* 1, *f*_1_(*x*) *>* 1 and *f*_2_(*x*) = 1, this will promote *Red* expression. Conversely, when |*x*| *<* 1, *f*_2_(*x*) *>* 1 and *f*_1_(*x*) = 1, this will promote *Green* expression. The specific forms of *f*_1_(*x*) and *f*_2_(*x*) are detailed in the Supplementary Note 3. cell proliferation modeled as division and apoptosis:

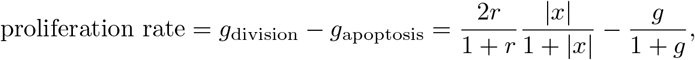

which means that the gene *Red* and migrating outward in the horizontal direction will promote cell proliferation while the gene *Green* will inhibit cell proliferation. We sampled three groups of cells at the initial moment and discretized time in order to obtain data through numerical simulations. We simulate gene expression and spatial coordinates according to the forward Euler scheme and simulate cell division and apoptosis by numerically simulating a special Markov process, the birth and death process. Specific details of the sampling of initial values and numerical simulation can be found in the Supplementary Note 3. We evolved the cells at the initial time point from *t* = 0 to *t* = 3.0 according to the given dynamics and took a total of six time points at *t* = 0, 0.5, 1.0, 1.5, 2.0 and 2.5 as observations. Considering that the spatial coordinates obtained at different time points using spatial transcriptome sequencing are not in the same coordinate system, we rotated the spatial coordinates of the second to sixth time points counterclockwise by 8, 16, 24, 32 and 40 degrees, respectively.

For the second simulation data corresponding to Fig. S2, it shares the similar gene expression dynamics as the first dataset and is divided into three groups. In terms of spatial migration, it simulates designated transformations with two arbitrary shapes (Fig. S2A). Additionally, we assume group-specific proliferation rates, which is calculated based on the initial and final cell numbers for each group. The data were sampled at five time points *t* = 0, 0.25, 0.5, 0.75, and 1, with the spatial coordinates at the second to fourth time points being rotated counterclockwise by -8, 16, -16, and 8 degrees, respectively. The detailed calculations can be found in Supplementary Note 3.

For the third simulation data corresponding to Fig. 5 and Fig. S13, similar to the first, the dynamics consists of three genes *Red, Green* and *Blue* and two spatial coordinates *x* and *y*. There are two types of cells in this simulation data, background cells and migratory cells. The background cells are in steady state, and their spatial coordinates and gene expression do not change with time. The migratory cells transited from high expression of *Red* and *Green* gene to high expression of only *Red* gene while moving to the right. Initial values for background cells and migratory cells can be found in the Supplementary Note 3. Gene expression and spatial coordinates of migrating cells evolve over time and obey stochastic dynamical systems

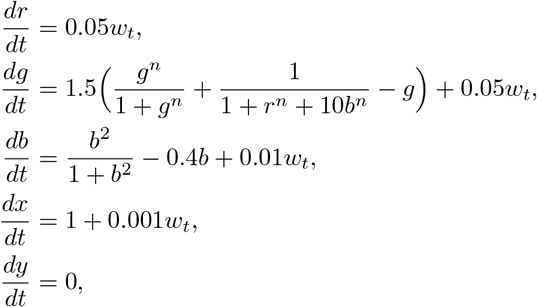

where we take *n* = 4 to model non-linear regulatory relationships. Unlike the first simulation data, we do not consider growth in the second simulation data. We evolved the cell at the initial time point from *t* = 0 to *t* = 1.0 according to the given dynamics. In Fig. 5, we took only two time points at *t* = 0 and 1.0 as observations. Additionally we rotated the spatial coordinates of the second time point by 8 degrees counterclockwise. In Fig. S13, we took a total of five time points at *t* = 0, 0.25, 0.5, 0.75 and 1 as observations and rotated the spatial coordinates of the second to fifth time points counterclockwise by 8, 16, 24 and 32 degrees, respectively.

### Details of GO enrichment analysis

We used the python package GSEApy=1.0.3 [76] to perform GO enrichment analyses on migration genes and growth genes. In addition, the gene sets used were GO Biological Process 2018 (https://maayanlab.cloud/Enrichr/#libraries).

## Data Availability

All the datasets used in this paper are publicly available. The simulation datasets of synthetic circuits are available at https://github.com/QiangweiPeng/stVCR/tree/main/tutorial. The axolotl brain regeneration datasets are freely accessible in CNGB Nucleotide Sequence Archive under accession code CNP0002068. Processed data can be downloaded from https://db.cngb.org/stomics/artista/ [57]. The processed 3D Drosophila embryo datasets can be downloaded from the Spateo package [50] (https://www.dropbox.com/s/bvstb3en5kc6wui/E7-9h_cellbin_tdr_v2.h5ad?dl=1 and https://www.dropbox.com/s/q02sx6acvcqaf35/E9-10h_cellbin_tdr_v2.h5ad?dl=1).

## Code Availability

stVCR is implemented in Python and is available at https://github.com/QiangweiPeng/stVCR. The notebooks to reproduce all the results in the manuscript are available at https://github.com/QiangweiPeng/stVCR/tree/main/tutorial.

## Acknowledgments

The authors thank the anonymous referees for careful reading and constructive suggestions. TL and QP acknowledge the support from National Key R&D Program of China under grant 2021YFA1003301, and National Science Foundation of China under grant 12288101. PZ acknowledge the support from National Science Foundation of China under grants 12288101, 8206100646, and the Fundamental Research Funds for the Central Universities. We also thank the High-performance Computing Platform of Peking University.

## Contribution

All authors conceived the project. QP designed and implemented the algorithm, and performed data analysis. All authors interpreted the results and wrote the manuscript. QP wrote the supplementary materials. PZ and TL supervised the research.

