## Supplementary Information (Figures, Tables, Notes, and Videos) for "stVCR: Spatiotemporal dynamics of single cells": Supplementary_Materials.pdf

### stVCR: Spatio-temporal dynamics of single cells

**References** **42**

### Supplementary Notes

#### Supplementary Note 1: Optimal transport and its variants related to stVCR

##### Wasserstein optimal transport

The Monge problem aims to find a mapping that assigns each point $\boldsymbol{x_i}$ to exactly one point $\boldsymbol{y_j}$, directing the mass of measure $\alpha$ towards that of measure $\beta$, where $\alpha = \sum_{i=1}^{n} a_i \delta(\boldsymbol{x} - \boldsymbol{x}_i)$ and $\beta = \sum_{j=1}^{m} b_j \delta(\boldsymbol{y} - \boldsymbol{y}_j)$. The assignment problem is combinatorial, and the feasible set for the Monge problem is nonconvex. Kantorovich suggests an alternative approach where the mass located at any point $x_i$ could potentially be dispatched across multiple locations. Kantorovich's optimal transport problem reads

$$\mathcal{L}_c(\alpha, \beta) = \min_{P \in U(\mathbf{a},\mathbf{b})} \langle C, P \rangle \stackrel{\text{def.}}{=} \sum_{i=1}^{n} \sum_{j=1}^{m} c_{ij} p_{ij}, \tag{1}$$

where $U(\mathbf{a}, \mathbf{b}) \stackrel{\text{def.}}{=} \left\{ P \in \mathbb{R}_+^{n \times m} : P \mathbb{1}_m = \mathbf{a} \quad \text{and} \quad P^{\mathrm{T}} \mathbb{1}_n = \mathbf{b} \right\}$.

##### Dynamical optimal transport

When $c(\boldsymbol{x}, \boldsymbol{y}) = \|\boldsymbol{x} - \boldsymbol{y}\|_2^2$, the optimal transport distance $\mathcal{L}_c(\alpha, \beta)$ as defined in Eq. (1) can be computed by finding a shortest path $(\alpha_t(\boldsymbol{x}))_{t=0}^{1}$ between these two measures. This path is described by advecting the measure using a vector field $\boldsymbol{v}_t(\boldsymbol{x})$ defined at each time point. The path $\alpha_t(\boldsymbol{x})$ and vector field $\boldsymbol{v}_t(\boldsymbol{x})$ must satisfy the mass conservation formula, yielding

$$\partial_t \alpha_t(x) + \nabla \cdot (\alpha_t(x) \boldsymbol{v}_t(x)) = 0 \quad \text{and} \quad \alpha_{t=0} = \alpha_0, \alpha_{t=1} = \alpha_1. \tag{2}$$

Benamou and Brenier [1] proved that

$$\mathcal{L}_c(\alpha, \beta) = \min_{(\alpha_t(\boldsymbol{x}), \boldsymbol{v_t}(\boldsymbol{x}))_t \text{ sat. } (2)} \int_0^1 \int_{\mathbb{R}^d} \|\boldsymbol{v}_t(\boldsymbol{x})\|^2 \alpha_t(\boldsymbol{x}) \, \mathrm{d}\boldsymbol{x} \mathrm{d}t. \tag{3}$$

##### Dynamical unbalanced optimal transport

To accommodate input measures with varying masses and address local mass discrepancies (unbalanced setting), many normalizations or relaxations have been proposed. A common approach involves incorporating a source term $s_t(\boldsymbol{x})$ into the continuity Eq. (2). Consequently, we consider

$$\overline{\mathcal{C}}\left(\alpha,\beta\right) \stackrel{\text{def.}}{=} \{(\alpha_t(\boldsymbol{x}),\boldsymbol{v}_t(\boldsymbol{x}),s_t(\boldsymbol{x})) : \partial_t\alpha_t(\boldsymbol{x}) + \nabla\cdot(\alpha_t(\boldsymbol{x})\boldsymbol{v}_t(\boldsymbol{x})) = s_t(\boldsymbol{x}), \\ \alpha_{t=0} = \alpha, \alpha_{t=1} = \beta\}\,. \tag{4}$$

The crucial question is how to measure the cost associated to this source term and introduce it in the original dynamic formulation, Wasserstein-Fisher-Rao (WFR) distance

$$(\text{WFR}\left(\alpha,\beta\right))^2 = \min_{(\alpha_t(\boldsymbol{x}),\boldsymbol{v}_t(\boldsymbol{x}),s_t(\boldsymbol{x}))_t\in\overline{\mathcal{C}}(\alpha,\beta)} \int_0^1\int_{\mathbb{R}^d}\left(\|\boldsymbol{v}_t(\boldsymbol{x})\|^2 \right. \\ \left. +\tau\frac{(s_t(\boldsymbol{x}))^2}{(\alpha_t(\boldsymbol{x}))^2}\right)\alpha_t(\boldsymbol{x})\;\mathrm{d}\boldsymbol{x}\mathrm{d}t. \tag{5}$$

was proposed to avoid having mass which travels at infinite speed and suddenly grows in a region where there was no mass before [2–4]. When describing biological processes, $s_t(\boldsymbol{x}) = g_t(\boldsymbol{x})\alpha_t(\boldsymbol{x})$ is often used, where $g_t(\boldsymbol{x})$ measures the rate of cell proliferation.

#### Gromov-Wasserstein optimal transport

Optimal transport relies on the ground cost $C$ to compare two distributions $\mathbf{a}$ and $\mathbf{b}$, but it does not apply when the two distributions are not defined on identical underlying spaces. To overcome this constraint, a weaker assumption is made, that two matrices $D \in \mathbb{R}^{n\times n}$ and $D' \in \mathbb{R}^{m\times m}$ measure the similarity among the points defining the own space. The GW problem [5] reads

$$\text{GW}\left((\mathbf{a},D),(\mathbf{b},D')\right)^2 \stackrel{\text{def.}}{=} \min_{P\in U(\mathbf{a},\mathbf{b})}\sum_{i,j,i',j'}\left|D_{i,i'} - D'_{j,j'}\right|^2 P_{i,j}P_{i',j'}. \tag{6}$$

When modeling spatial coordinates using GW-OT, the matrices $D$ and $D'$ typically use the Euclidean distance between points. This is a reasonable choice in order to obtain a optimal mapping $P$ because the GW-OT is rigid body transformation invariant in this setting. However, dynamical OT requires that distributions at different time points are defined on identical underlying spaces. The GW-OT does not align the spatial coordinates of data at different time points to the same coordinate system, and it is still not possible to compute the ground cost between the spatial coordinates of the data at different time points, and therefore cannot be generalized to dynamical OT.

#### Optimal transport under an invariant set

Cohen and Guibas [6] explicitly model optimal transport under an invariant set $\mathcal{G}$, defining the problem as simultaneously finding the optimal matching and the optimal transformation

$$P^{\star}, g^{\star} = \arg \min_{P \in U(\mathbf{a},\mathbf{b}), g \in \mathcal{G}} \langle C(g), P \rangle \overset{\text{def.}}{=} \sum_{i=1}^{m} \sum_{j=1}^{n} P_{ij} d\left(x_i, g\left(y_j\right)\right), \tag{7}$$

and design a monotonically iterative algorithm

$$P^{(k)} = \arg \min_{P \in U(\mathbf{a},\mathbf{b})} \sum_{i=1}^{m} \sum_{j=1}^{n} P_{ij} d\left(x_i, g^{(k)}\left(y_j\right)\right), \tag{8}$$

$$g^{(k+1)} = \arg \min_{g \in \mathcal{G}} \sum_{i=1}^{m} \sum_{j=1}^{n} P_{ij}^{(k)} d\left(x_i, g\left(y_j\right)\right). \tag{9}$$

Subproblem (8) is to solve a static OT. When the invariant set $\mathcal{G}$ is taken as the set of rigid body transformations, we call the problem rigid body transformation invariant OT (RIT-OT).

#### Fused rigid body transformation invariant optimal transport

For spatial transcriptome data, there are gene expressions in addition to spatial coordinates. To measure the distance between two spatial transcriptome data, we consider the fused rigid body transformation invariant OT (FRIT-OT), as in fused GW-OT [7]. Assuming that $\mathcal{G}$ is a set of rigid body transformations and $C^{exp}$ refers to the distance matrix between gene expressions and $w$ is a weight coefficient, the FRIT-OT problem reads

$$\begin{aligned} P^{\star}, g^{\star} &= \arg \min_{P \in U(\mathbf{a},\mathbf{b}), g \in \mathcal{G}} \langle wC(g) + (1-w)C\text{exp}, P \rangle \\ &\overset{\text{def.}}{=} \sum_{i=1}^{m} \sum_{j=1}^{n} P_{ij} \left(wd\left(x_i, g\left(y_j\right)\right) + (1-w)C^{\text{exp}}(i,j)\right), \end{aligned} \tag{10}$$

can be solved iteratively similar to Eq 7. In the following quantitative comparison, we choose this index to measure the distance between the interpolated ST data and the observed data.

### Supplementary Note 2 Computational complexity analysis of stVCR

We rigorously analyze the computational demands of the stVCR framework through time and space complexity formalisms. Let $N$ denote the average cell number per observation time point, $K+1$ the number of time points, $T$ the total length of the integration interval, $\delta_t$ the step size for ODE solvers, $W$ the parameter dimension of the neural networks, and $B$ the mini-batch size.

#### Time Complexity

The epoch-level time complexity originates from two computational primitives:

- **Bidirectional Neural ODE Integration**: Each training iteration requires forward and backward integration of the neural ODE system. The forward pass from $t_0$ to $t_{K+1}$ for a mini-batch of $B$ cells incurs:

$$\mathcal{O}\left(B\frac{T}{\delta_t}W\right) \tag{11}$$

  where $\frac{T}{\delta_t}$ represents the number of integration steps. The backward propagation via adjoint sensitivity method [8] maintains the same asymptotic complexity $\mathcal{O}\left(B\frac{T}{\delta_t}W\right)$. The reverse integration from $t_{K-1}$ to $t_0$ is similar to from $t_0$ to $t_{K-1}$. So the time complexity of this part is

$$\mathcal{T}_{\text{ode}} = \mathcal{O}\left(2(B\frac{T}{\delta_t}W + B\frac{T}{\delta_t}W)\right) = \mathcal{O}\left(4B\frac{T}{\delta_t}W\right) \tag{12}$$

- **Optimal Transport Computation**: The time complexity of an exact static OT of size B is $\mathcal{O}\left(B^3 \log B\right)$ [9]. A total of $2K$ exact OT problem need to be calculated for each iteration, so the time complexity of this part is

$$\mathcal{T}_{\text{ot}} = \mathcal{O}\left(2KB^3 \log B\right) \tag{13}$$

Aggregating these components over $\frac{N}{B}$ iterations per epoch yields the total time complexity:

$$\begin{aligned} \mathcal{T}_{\text{epoch}} &= \mathcal{O}\left(\frac{N}{B}\left(\mathcal{T}_{\text{ode}} + \mathcal{T}_{\text{ot}}\right)\right) \\ &= \mathcal{O}\left(4N\frac{T}{\delta_t}W + 2NKB^2 \log B\right) \\ &= \mathcal{O}\left(N(\frac{T}{\delta_t}W + KB^2 \log B)\right) \end{aligned} \tag{14}$$

#### Memory Complexity

The space complexity per epoch is dominated by:

- **Bidirectional Neural ODE Integration**: The adjoint sensitivity method [8] eliminates the need to store intermediate states during ODE integration, reducing memory consumption to:

$$\mathcal{M}_{\text{ode}} = \mathcal{O}(2BW) \tag{15}$$

- **Transport Cost Matrices**: Each static optimal transport computation requires storage of a pairwise cost matrix. For $2K$ concurrent computations:

$$\mathcal{M}_{\text{ot}} = \mathcal{O}\left(2KB^2\right) \tag{16}$$

The total space complexity therefore combines Equations 15 and 16:

$$\mathcal{M}_{\text{total}} = \mathcal{O}\left(2BW + 2KB^2\right) = \mathcal{O}\left(BW + KB^2\right) \tag{17}$$

In practice, batch size $B$ and the integral step size $\delta_t$ are usually fixed constants, so the time complexity of each epoch

$$\mathcal{T}_{\text{epoch}} = \mathcal{O}\left(NTW + K\right) \tag{18}$$

and the total space complexity is .

$$\mathcal{M}_{\text{total}} = \mathcal{O}\left(W + K\right). \tag{19}$$

In summary, the above analysis shows that the time complexity of stVCR is linear with respect to data size $N$, integral interval $T$, model size $W$ and the number of observation time points $K+1$, while the space complexity is independent of data size $N$ and integral interval $T$, and is linear only with respect to model size $W$ and the number of observation points $K+1$.

#### Supplementary Note 3: Details of simulated data generation

In this paper, three simulation datas are included, the first corresponding to Fig. 2, the second corresponding to Fig. S2, and the third corresponding to Fig. 5 and Fig. S13. Below, we will introduce how these three simulation data are generated.

##### Simulated data 1

For the first simulation data corresponding to Fig. 2, the dynamics of the simulated data consists of three genes *Red*, *Green* and *Blue* and two spatial coordinates $x$ and $y$, whose regulation is shown in Fig. 2A. Such a regulatory relationship can be described by a system of stochastic differential equations for gene expression and spatial coordinates

$$\begin{aligned}
\frac{dr}{dt} &= f_1(x)(\frac{r^n}{1+r^n} + \frac{1}{1+g^n+10b^n} - r) + 0.05w_t \\
\frac{dg}{dt} &= f_2(x)(\frac{g^n}{1+g^n} + \frac{1}{1+r^n+10b^n} - g) + 0.05w_t \\
\frac{db}{dt} &= \frac{b^2}{1+b^2} - 0.4b + 0.01w_t \\
\frac{dx}{dt} &= \text{sign}(x)\exp(-4b)\exp(-4g)(r-2)^2r^2 + 0.001w_t \\
\frac{dy}{dt} &= 0
\end{aligned} \tag{20}$$

where $r$, $g$ and $b$ refer to gene *Red*, *Green* and *Blue*, $f_1(x)$ and $f_2(x)$ refer to the factors that depend on the coordinates $x$, and $w_t$ is standard Brownian motion. In the calculation, we take $n = 4$ to simulate the nonlinear regulation between genes. If we ignore these two x-related factors $f_1(x)$ and $f_2(x)$, $r$ and $g$ are a toggle switch of equal status. The specific definitions of $f_1(x)$ and $f_2(x)$ are

$$f_1(x) = \begin{cases} \max(\exp(0.75(x-1)), 1), & x > 0 \\ -f_1(-x), & x < 0 \end{cases} \tag{21}$$

and

$$f_2(x) = \begin{cases} \max(\exp(0.75(1-x)), 1), & x > 0 \\ -f_2(-x), & x < 0. \end{cases} \tag{22}$$

When $|x| > 1$, $f_1(x) > 1$ and $f_2(x) = 1$, this will promote *Red* expression. Conversely, when $|x| < 1$, $f_2(x) > 1$ and $f_1(x) = 1$, this will promote *Green* expression. Cell growth modeled as division and apoptosis

$$\text{growth} = g_{\text{division}} - g_{\text{apoptosis}} = \frac{2r}{1+r}\frac{|x|}{1+|x|} - \frac{g}{1+g}, \tag{23}$$

which means that the gene *Red* and migrating outward in the horizontal direction will promote cell growth while the gene *Green* will inhibit cell growth. We discretized time in order to obtain data through numerical simulations. Within the time step $\delta_t$, we simulate gene expression and spatial coordinates after $\delta_t$ according to the forward Euler scheme and simulate cell division and apoptosis by numerically simulating a special Markov process, the birth and death process. More specifically, cell division or apoptosis is determined by the growth function, specifically within the $\delta_t$, the probability of cell division is $p_{\text{division}} = g_{\text{division}}\delta_t$, the probability of apoptosis is $p_{\text{apoptosis}} = g_{\text{apoptosis}}\delta_t$, and the probability of remaining unchanged is $1 - p_{\text{division}} - p_{\text{apoptosis}}$. When a cell divides into two, we add white noise with a standard deviation of 0.01 to the gene expression and spatial coordinates of the original cell and become the data of the new cell. In addition, at each step of the simulation when there is a negative value of gene expression, we force it to be corrected to 0.

At the initial time point we randomly sampled three groups of cells. The gene expression $(r, g, b)$ of the first group of cells was sampled from the normal distribution with mean $(0, 0, 2)$ and covariance matrix of $0.1I$, and the spatial coordinates $(x, y)$ were sampled from the uniform distribution $[-2, 2] \times [-3, 3]$. Gene expression in both the second and third groups of cells was sampled from the normal distribution with mean $(0, 0, 0)$ and covariance matrix of $0.1I$, in addition the spatial coordinates of the second group of cells were sampled from a uniform distribution $[-2, -1.5] \cap [1.5, 2] \times [-3, 3]$, and the spatial coordinates of the third group of cells were sampled from a uniform distribution $[-0.5, 0.5] \times [-3, 3]$. We evolved the cell at the initial time point from t=0 to t=3.0 according to the given dynamics and took a total of six time points at t=0, 0.5, 1.0, 1.5, 2.0 and 2.5 as observations. Considering that the spatial coordinates obtained at different time points using spatial transcriptome sequencing are not in the same coordinate system, we rotated the spatial coordinates of the second to sixth time points counterclockwise by 8, 16, 24, 32 and 40 degrees, respectively.

#### Simulated data 2

For the second simulation data corresponding to Fig. S2, it shares the similar gene expression dynamics as the first dataset and is divided into three groups

$$\begin{aligned}
\frac{dr}{dt} &= f_1(\text{group id})\left(\frac{r^n}{1+r^n} + \frac{1}{1+g^n+10b^n} - r\right) + 0.05w_t \\
\frac{dg}{dt} &= f_2(\text{group id})\left(\frac{g^n}{1+g^n} + \frac{1}{1+r^n+10b^n} - g\right) + 0.05w_t \\
\frac{db}{dt} &= \frac{b^2}{1+b^2} - 0.4b + 0.01w_t,
\end{aligned} \tag{24}$$

where

$$f_1(\text{group id}) = \begin{cases} 1.5, & \text{group id} = 2 \\ 1, & \text{other} \end{cases} \tag{25}$$

and

$$f_2(\text{group id}) = \begin{cases} 1.5, & \text{group id} = 1 \\ 1, & \text{other} \end{cases} \tag{26}$$

The initial gene expression values for each group are same as those in the first simulated dataset. In terms of spatial migration, it simulates designated transformations with two arbitrary shapes (Fig. S2A). The initial $(t = 0)$ cell positions $(x_i^{\text{id}}(0), y_i^{\text{id}}(0))$ and the number of cells $n^{\text{id}}(0)$ in each group as well as the final cell positions $(x_j^{\text{id}}(1), y_j^{\text{id}}(1))$ and the number of cells $n^{\text{id}}(1)$, are obtained through uniform sampling within the respective regions of the initial and final shapes. The spatial migration velocity for each group is determined by simple static OT interpolation, that is $v_i^{\text{id}} = n^{\text{id}}(0) \sum_{j=1}^{n^{\text{id}}(1)} P_{ij}^{\text{id}}(x_j^{\text{id}}(1), y_j^{\text{id}}(1)) - (x_i^{\text{id}}(0), y_i^{\text{id}}(0))$. Assuming that the growth rate is group-specific, it can be computed by $g^{\text{id}} = \ln \frac{n^{\text{id}}(1)}{n^{\text{id}}(0)}$. We evolved the cell at the initial time point from t=0 to t=1.0 according to the dynamics and took a total of five time points at t=0, 0.25, 0.5, 0.75 and 1.0 as observations. Finally, the spatial coordinates at the second to fourth time points being rotated counterclockwise by -8, 16, -16, and 8 degrees, respectively.

#### Simulated data 3

For the second simulation data corresponding to Fig. 5 and Fig. S13, similar to the first, the dynamics of the simulated data consists of three genes *Red*, *Green* and *Blue* and two spatial coordinates $x$ and $y$. There are two types of cells in this simulation data, background cells and migratory cells. The background cells are in steady state, and their spatial coordinates and gene expression do not change with time. The spatial coordinates of the background cells follow a uniform distribution of region $D$, where the left half of $D$ consists of the left half of an ellipse with major axis 2.25 and minor axis 1, and the right half of $D$ similarly consists of the right half of an ellipse with major axis 1.5 and minor axis 1. The gene expression of background cells obeys a Gaussian distribution with mean $(1.1, 1, 0)$ and covariance matrix $0.001I$. The spatial coordinates of migrating cells at the initial moment obey a uniform distribution on a circle with circle centre $(-1, 0)$ and radius 0.3. Their gene expression obeys a Gaussian distribution with mean $(2, 2, 0)$ and covariance matrix $0.001I$. Gene expression and spatial coordinates of migrating cells evolve over time and obey stochastic dynamical systems

$$\begin{aligned}
\frac{dr}{dt} &= 0.05w_t \\
\frac{dg}{dt} &= 1.5\left(\frac{g^n}{1+g^n} + \frac{1}{1+r^n+10b^n} - g\right) + 0.05w_t \\
\frac{db}{dt} &= \frac{b^2}{1+b^2} - 0.4b + 0.01w_t \\
\frac{dx}{dt} &= 1 + 0.001w_t \\
\frac{dy}{dt} &= 0,
\end{aligned} \tag{27}$$

where we take $n = 4$ to model non-linear regulatory relationships. Unlike the first simulation data, we do not consider growth in the second simulation data. We evolved the cell at the initial time point from t=0 to t=1.0 according to the given dynamics. In Fig. 5, we took only two time points at t=0 and 1.0 as observations. Additionally we rotated the spatial coordinates of the second time point by 8 degrees counterclockwise. In Fig. S13, we took a total of five time points at t=0, 0.25, 0.5, 0.75 and 1 as observations and rotated the spatial coordinates of the second to fifth time points counterclockwise by 8, 16, 24 and 32 degrees, respectively.

#### Supplementary Note 4: Details of simulated data analysis

##### Simulated data 1

We tested the performance of stVCR on the simulated dataset 1 (Fig. 2, Fig. S1). There are regulatory relationships between different genes and different spatial coordinates, in addition, gene expression and cell migration also affect cell growth (Fig. 2A). The three genes are named *Red*, *Green* and *Blue* genes. *Blue* is the upstream gene, and high expression of *Blue* will inhibit *Red* and *Green*. *Red* and *Green* are a toggle-switch, which self-activate and inhibit each other. In addition, *Red* promotes cell growth and *Green* inhibits cell growth. The horizontal direction in spatial coordinates (i.e., the x-axis) is the key to the simulated data. Cells moving outward horizontally (i.e., $|x|$ is larger) will promote *Red* expression and cell growth, while the opposite will promote *Green* expression. In summary, the difference in spatial location makes the two genes in the toggle-switch unequal in status. The cells at initial time point included three groups of cells (Fig. 2 B). The gene expression of the second and third groups at initial time point is similar, all genes are underexpressed, but the spatial location of the second group is on the outside and the third group is on the inside (Fig. 2B Left). While the second group of cells migrated outwards in the horizontal direction, they continuously divided and the *Red* gene was gradually highly expressed. The third group of cells showed gradually high expression of *Green* gene and continued apoptosis. Without spatial information, it is not possible to distinguish between the second and third group of cells at the initial moment, which would lead to erroneous trajectories (Fig. 2B Left and Fig. S5 last row). The first group of cells had high *Blue* expression and were in steady state, which remained virtually unchanged over time. Notice that the second group of cells is constantly dividing and increasing in number, while the third group of cells is constantly apoptotic and decreasing in number. If balanced OT is used for modeling and cell growth is ignored it will lead to wrong cell trajectories of the small number group to the large number group (Fig. 2B Right and Fig. S6 last row). We took as input data at t=0.0, 0.5, 1.0, 1.5, 2.0 and 2.5 for a total of 6 time points. In addition, we rotated the spatial coordinates of the second to sixth time points by 8, 16, 24, 32, and 40 degrees counterclockwise, respectively, to simulate the possible rotation of tissues by spatial transcriptome sequencing (Fig. 2C and Fig. S1A).

First, we take the data from the first time point and evolve them according to the learned dynamics. The results show that the learned dynamics are close to the real dynamics (Supplementary Video 1; Fig. 2D and Fig. S1B). The first group of cells remained virtually unchanged. The second group of cells gradually overexpressed the *Red* gene, moved outwards in the horizontal direction, and continuously proliferated. The third group of cells gradually overexpressed *Green* gene and continued apoptosis. In addition, we observed that the cells only moved horizontally and did not rotate, indicating that we found the optimal rigid body transformation to align the data at different time points while finding the optimal dynamics of cell evolution (Supplementary Video 1, Fig. 2D and Fig. S1B). And then we interpolate the unobserved moments t=0.25, 0.75, 1.25, 1.75 and 2.25 and the results are close to ground truth (Fig. 2E and Fig. S1C). We take the data at the last time point and make the prediction for t=2.75 based on the learned dynamics, which is also close to the ground truth (Fig. 2F). To investigate the ability of stVCR to restore the effects of gene interactions, we calculate the partial derivatives of *Green* with respect to *Red* in the learned dynamics and true dynamics and visualize them in coordinate space (Fig. 2G Left). Qualitatively, they were close, and *Green* inhibited *Red* expression mainly in the second and third group of cells. To investigate the ability of stVCR to restore the effects cell migration on gene expression, we calculate the directional derivative of *Red* gene expression for the given direction $n = (1, 0)$ (i.e., cells moving horizontally to the right) for both learned and true dynamics (Fig. 2G Right). Cells at the right end of the second group moving to the right will promote gene *Red* expression, and cells at the left end moving to the right will inhibit gene *Red* expression, which overall suggests that moving horizontally outward in the second group of cells will promote *Red* expression. Finally, we investigated the spatial variability of cell growth and the effect of cell migration on growth. We compared true and learned cell growth rates and visualized them in coordinate space (Fig. 2H Left). The results show that the first group of cells has a growth rate close to 0, the second group has a large positive growth rate, and the third group has a large negative growth rate. Additionally we calculated the directional derivative of the growth rate $g$ with respect to a given direction $n = (1, 0)$, similarly this shows that cells moving outward in the horizontal direction will promote cell growth (Fig. 2H Right).

Finally, we checked the scalability of the stVCR and its robustness with respect to important hyperparameters on this simulated dataset. First, we tested the time-consumption of the stVCR for different sizes of datasets (measured by the number of cells at the initial moment $t_0$), the number of observation time points, the batch size of the sampling, and the model size (measured by the number of hidden layers) (Fig. S3). The results show that the time-consumption of stVCR increases linearly with the size of the dataset, the number of observation time points, and the model size, while it decreases and then increases rapidly with the batch size of sampling. We speculate that the possible reason is that when the batch size is small, the parallel computing power of GPU is not fully utilized; when the batch size is large, the time consumed increases rapidly because the computational complexity of static OT is $O(n^3)$ and each iteration needs to compute the static OT of the batch size scale. We then tested the robustness of stVCR with respect to the important hyperparameters $\lambda_{\text{Mch}}$ (Fig. S4 and Supplementary Video 2), $\kappa_{\text{Exp}}$ (Fig. S5 and Supplementary Video 3), and $\alpha_{\text{Gro}}$ (Fig. S6 and Supplementary Video 4). Parameter $\lambda_{\text{Mch}}$ measures the importance of the loss term $\mathcal{L}_{\text{Mch}}$ (Methods). When $\lambda_{\text{Mch}}$ is small, gene expression and spatial location of all cells are unchanged (Fig. S4 second row), whereas when $\lambda_{\text{Mch}}$ is large enough, stVCR obtains correct results and is robust with respect to $\lambda_{\text{Mch}}$ (Fig. S4). Parameter $\kappa_{\text{Exp}}$ weighs the importance of gene expression and spatial coordinates in the matching term, when $\kappa_{\text{Exp}} = 0$ means that only spatial coordinates is considered, and conversely when $\kappa_{\text{Exp}} = 1$ means that only gene expression are considered (Methods). The results show that incorrect dynamics are produced when either only spatial coordinates (Fig. S5 second row) or only gene expression (Fig. S5 last row) are considered, while the results are correct and robust when both are considered (Fig. S5). The parameter $\alpha_{\text{Gro}}$ measures the cost of cell growth, with lower values promoting cells that can divide and apoptose more flexibly, and higher values promoting all cells to have the same growth rate (equivalently, balanced OT) (Methods). The results show that the results are correct and robust when $\alpha_{\text{Gro}}$ is low, and wrong transitions from green to blue cells and blue to red cells are observed when $\alpha_{\text{Gro}}$ is large (Fig. S6).

#### Simulated data 2

We tested the performance of stVCR on the simulated dataset 2 to demonstrate its ability to learn arbitrary shape transformations (Fig. S2). This simulation describes the transition between two cartoon images, from a heart shape to a duck (Fig. S2A, B). stVCR accurately aligned the spatial coordinates of observations at different time points (Fig. S2B, C) and successfully learned the overall dynamics (Supplementary Video 2). Moreover, stVCR effectively interpolated unobserved time points and inferred the proliferation rate (Fig. S2D, E). Finally, ablation experiments demonstrated that all components of stVCR, including spatial information, gene expression, proliferation, and alignment, are essential for learning the complete dynamics (Supplementary Video 2).

#### Simulated data 3

We tested the performance of stVCR on the simulated dataset 3 to illustrate the necessity to add known biological priors for data with long observation intervals and fewer observation time points to obtain more accurate results (Fig. 5 and Fig. S13). The simulated data consists of three types of cells type1, type2, and type3 (Fig. 5A). The type3 expresses the *Red* and *Green* genes but neither to a high degree, type1 and type2 cells highly express the *Red* gene, and in addition to this type1 cells also highly express the *Green* gene (Fig. 5A). The type3 cells are already at steady state and no longer changing, while the *Green* gene expression of type1 cells gradually decreases and migrates positively along the x-axis transitioning to type2 cells over time (Fig. 5A,B).We assumed that the observed data only contained data at t=0 and t=1 (Fig. 5C), and the information at the intermediate moments was not observed (Fig. 5A,C). At this time, both the spatial location and gene expression of type3 cells are more close to type1 and type2 cells, while type1 and type2 cells are more different. Therefore, we speculate that the stVCR will incorrectly infer type3 to type1 and type3 to type2 transitions rather than the correct type1 to type2 transition if no additional information is provided about the type1 to type2 cell transition (Fig. 5A). We use stVCR with and without the biological knowledge prior of type1 to type2 transition to reconstruct the whole dynamic process, and the results show that the stVCR with priors correctly reconstructs the whole dynamic process as expected (Supplementary Video S7), while the stVCR without priors fails (Supplementary Video S8). We visualize the interpolation results and the results show that the stVCR with prior is consistent with ground truth, while the stVCR without prior is much different (Fig. 5D and Fig. S12). The above experimental results illustrate the benefit of adding the correct biological prior for data with fewer observations and longer intervals to obtain more accurate results. It is worth noting that when there are enough observation time points and the time intervals are small enough, the correct result can be inferred by stVCR even without a prior. We theoretically prove that the optimal transport dynamics will converge to the true dynamics when the time intervals between neighboring observation time points converges to 0, which provides a theoretical guarantee for our method (Supplementary Note 4). We still utilize the simulated dataset of Fig. 5A, but increase the observation time points to a total of five time points at t = 0, 0.25, 0.5, 0.75, 1.0 (Fig. S13A). The results show that the entire dynamics are accurately recovered by stVCR without prior (Supplementary Video S9). We visualize the interpolated results at t=0.125, 0.375, 0.625 and 0.875 and the stVCR results is close to the ground truth (Fig. S13B). However, due to the high cost of ST data, each additional moment of data will increase the total cost, so it is still advantageous to model known cell type transitions as an option.

#### Supplementary Note 5: Proof of convergence of optimal transport dynamics

We show that under certain conditions trajectory data can be recovered from snapshot data when the time interval between observations is sufficiently small, which allows the OT-based dynamics to converge to the true dynamics. This provides a theoretical guarantee for the use of OT to model time-series sequencing data.

Consider the initial distribution $\rho(0) = \frac{1}{N}\Sigma_{i=1}^{N}\delta(\boldsymbol{x}_i(0))$, which means that we consider a finite number of data points. The distribution $\rho(t) = \frac{1}{N}\Sigma_{i=0}^{N}\delta(\boldsymbol{x}_i(t))$ for subsequent times is obtained by evolving the initial distribution $\rho(0)$ according to the true dynamics $\boldsymbol{v}_{\text{true}}(\boldsymbol{x}, t)$, i.e. $\boldsymbol{x}_i(t)$ is obtained by solving the initial value problem

$$\begin{cases} \dfrac{d\boldsymbol{x}}{dt} = \boldsymbol{v}_{\text{true}}(\boldsymbol{x}, t) \\ \boldsymbol{x}(0) = \boldsymbol{x}_i(0) \end{cases} \tag{28}$$

The observation time interval is $\delta_t$, so the data collected is $\rho(k\delta_t) = \frac{1}{N}\Sigma_{i=0}^{N}\delta(\boldsymbol{x}_i(k\delta_t)), k = 0, 1, 2 \ldots K$. Assume that condition

$$\min\{\|\boldsymbol{x}_i(0) - \boldsymbol{x}_j(0)\| | \forall i \neq j, i, j = 1, 2 \ldots N\} := a(0) > 0, \tag{29}$$

condition

$$\exists M > 0, \|\boldsymbol{v}_{\text{true}}(\boldsymbol{x}, t)\| < M, \forall \boldsymbol{x} \in \mathbb{R}^n, t \in \mathbb{R}^+, \tag{30}$$

and condition

$$\exists L > 0, \|\boldsymbol{v}_{\text{true}}(\boldsymbol{x}, t) - \boldsymbol{v}_{\text{true}}(\boldsymbol{y}, t)\| < L\|\boldsymbol{x} - \boldsymbol{y}\|, \forall \boldsymbol{x}, \boldsymbol{y} \in \mathbb{R}^n, t \in \mathbb{R}^+, \tag{31}$$

are satisfied, where the Eq. (29) implies that each data point at the initial moment is different, and the Eq. (30) implies that the true dynamics $\boldsymbol{v}_{\text{true}}(\boldsymbol{x}, t)$ are bounded and the Eq. (31) implies that true dynamics $\boldsymbol{v}_{\text{true}}(\boldsymbol{x}, t)$ satisfies the Lipschitz condition for the variable $\boldsymbol{x}$. According to Eq. (31), the initial value problem in Eq. (28) satisfies the existence and uniqueness of the solution. Also according to Eq. (29), the initial values of the orbits $\boldsymbol{x}_i(t)$ are different, so they will not intersect, i.e. $\boldsymbol{x}_i(t) \neq \boldsymbol{x}_j(t), \forall i \neq j$, which will result in

$$\min\{\|\boldsymbol{x}_i(t) - \boldsymbol{x}_j(t)\| | \forall i \neq j, i, j = 1, 2 \dots N\} := a(t) > 0. \tag{32}$$

Below we show that the OT-based mapping of two neighbouring time points is always correct when $\delta_t$ is small enough. We consider the distributions $\rho(k\delta_t) = \frac{1}{N}\Sigma_{i=0}^{N}\delta(\boldsymbol{x}_i(k\delta_t))$ and $\rho((k+1)\delta_t) = \frac{1}{N}\Sigma_{i=0}^{N}\delta(\boldsymbol{x}_i((k+1)\delta_t))$ for the $k$ and $k+1$ time points. According to the Taylor expansion we have

$$\boldsymbol{x}_i((k+1)\delta_t) - \boldsymbol{x}_i(k\delta_t) = \boldsymbol{v}_{\text{true}}(\boldsymbol{x}_i(k\delta_t), k\delta_t)\delta_t + o(\delta_t^2),$$

and then according to Eq. (30)

$$\|\boldsymbol{x}_i((k+1)\delta_t) - \boldsymbol{x}_i(k\delta_t)\| = \|\boldsymbol{v}_{\text{true}}(\boldsymbol{x}_i(k\delta_t), k\delta_t)\delta_t + o(\delta_t^2)\| \leq M\delta_t + o(\delta_t^2).$$

But for any $j \neq i$, when $\delta_t \to 0$ we have

$$\begin{aligned}
\|\boldsymbol{x}_j((k+1)\delta_t) - \boldsymbol{x}_i(k\delta_t)\| &= \|\boldsymbol{x}_j((k+1)\delta_t) - \boldsymbol{x}_j(k\delta_t) + \boldsymbol{x}_j(k\delta_t) - \boldsymbol{x}_i(k\delta_t)\| \\
&\geq \|\boldsymbol{x}_j(k\delta_t) - \boldsymbol{x}_i(k\delta_t)\| - \|\boldsymbol{x}_j((k+1)\delta_t) - \boldsymbol{x}_j(k\delta_t)\| \\
&\geq a(k\delta_t) - M\delta_t + o(\delta_t^2) \\
&> \|\boldsymbol{x}_i((k+1)\delta_t) - \boldsymbol{x}_i(k\delta_t)\|.
\end{aligned} \tag{33}$$

Therefore, when $\delta_t \to 0$, the OT-based mapping is correct. We can recover the trajectory $\hat{\boldsymbol{x}}_i(t)$ and dynamics $\hat{\boldsymbol{v}}(\boldsymbol{x}, t)$ from the observed snapshot data, and more specifically when $t \in [k\delta_t, (k+1)\delta_t), k = 0, 1 \dots K$, we have

$$\begin{aligned}
\hat{\boldsymbol{x}}_i(t) &= \boldsymbol{x}_i(k\delta_t) + \frac{t - k\delta_t}{\delta_t}(\boldsymbol{x}_i((k+1)\delta_t) - \boldsymbol{x}_i(k\delta_t)), \\
\hat{\boldsymbol{v}}(\hat{\boldsymbol{x}}_i(t), t) &= (\boldsymbol{x}_i((k+1)\delta_t) - \boldsymbol{x}_i(k\delta_t))/\delta_t.
\end{aligned} \tag{34}$$

which is equivalent to piecewise linear interpolation for the correct orbital data $\boldsymbol{x}_i(t)$, and the convergence of $\hat{\boldsymbol{x}}_i(t) \to \boldsymbol{x}_i(t)$ and $\hat{\boldsymbol{v}}(\boldsymbol{x}, t) \to \boldsymbol{v}_{\text{true}}(\boldsymbol{x}, t)$ is guaranteed by the convergence of the piecewise linear interpolation.

#### Supplementary Note 6 Pseudocode for stVCR training

**Algorithm 1:** stVCR Training

**Input:** Neural ODEs $v_\theta$, $p_\theta$, $g_\theta$;

$K$ rotation angles $\alpha_{1:K}$ (or $\alpha_{1:K}, \beta_{1:K}, \gamma_{1:K}$ for 3D case); $K$ translation vectors $r_{1:K}$;

$B$ batch size; Other hyperparameters

**while** *training* **do**

  **for** $k = 1$ **to** $K$ **do**

    Apply rigid transformations $\alpha_k$ and $r_k$ to spatial coordinates at $k$th observation time points ;

  **end**

  Sample data at initial time $t_0$: $(x^{(t_0)}_{1:B}, q^{(t_0)}_{1:B}) \sim \rho^{(t_0)}$, $w^{(t_0)}_{1:B} = \mathbf{1}_{1:B}$;

  **for** $k = 1$ **to** $K$ **do**

    Compute via NeuralODE solver:;

      $(x(t_k)_{1:B}, q(t_k)_{1:B}, w(t_k)_{1:B}), L^0_{\text{Dyn}}(t_k), L^0_{\text{SSP}}(t_k)^{\text{opt}} =$

      NeuralODESolver$(x^{(t_0)}_{1:B}, q^{(t_0)}_{1:B}, w^{(t_0)}_{1:B})$ (see equations (18)–(20) and (24));

    Sample data at $t_k$: $(x^{(t_k)}_{1:B}, q^{(t_k)}_{1:B}) \sim \rho^{(t_k)}$, $w^{(t_k)}_{1:B} = \mathbf{1}_{1:B}$;

    Calculate matching error $L^0_{\text{Mch}}(t_k)$ (see equation (21));

  **end**

  Sample data at final time $t_K$: $(x^{(t_K)}_{1:B}, q^{(t_K)}_{1:B}) \sim \rho^{(t_K)}$, $w^{(t_K)}_{1:B} = \mathbf{1}_{1:B}$;

  **for** $k = K - 1$ **to** $0$ **do**

    Compute via NeuralODE solver:;

      $(x(t_k)_{1:B}, q(t_k)_{1:B}, w(t_k)_{1:B}), L^{-1}_{\text{Dyn}}(t_k), L^{-1}_{\text{SSP}}(t_k)^{\text{opt}} =$

      NeuralODESolver$(x^{(t_0)}_{1:B}, q^{(t_0)}_{1:B}, w^{(t_0)}_{1:B})$ (see equations (18)–(20) and (24));

    Sample data at $t_k$: $(x^{(t_k)}_{1:B}, q^{(t_k)}_{1:B}) \sim \rho^{(t_k)}$, $w^{(t_k)}_{1:B} = \mathbf{1}_{1:B}$;

    Calculate matching error $L^{-1}_{\text{Mch}}(t_k)$ (see equation (21));

  **end**

  Compute total loss $L = L^0_{\text{Dyn}}(t_K) + L^{-1}_{\text{Dyn}}(t_0) + \lambda_{\text{Mch}}(\sum L^{-1}_{\text{Mch}}(t_k) + \sum L^0_{\text{Mch}}(t_k)) + \lambda_{\text{SSP}}(L^0_{\text{SSP}}(t_K)^{\text{opt}} + L^{-1}_{\text{SSP}}(t_0))^{\text{opt}}$ (see equation (25));

  Update dynamics parameters $\theta$ and rigid transformation parameters $\alpha_{1:K}$, $r_{1:K}$ via backpropagation;

**end**

### Supplementary Figures

#### Supplementary Figure S1

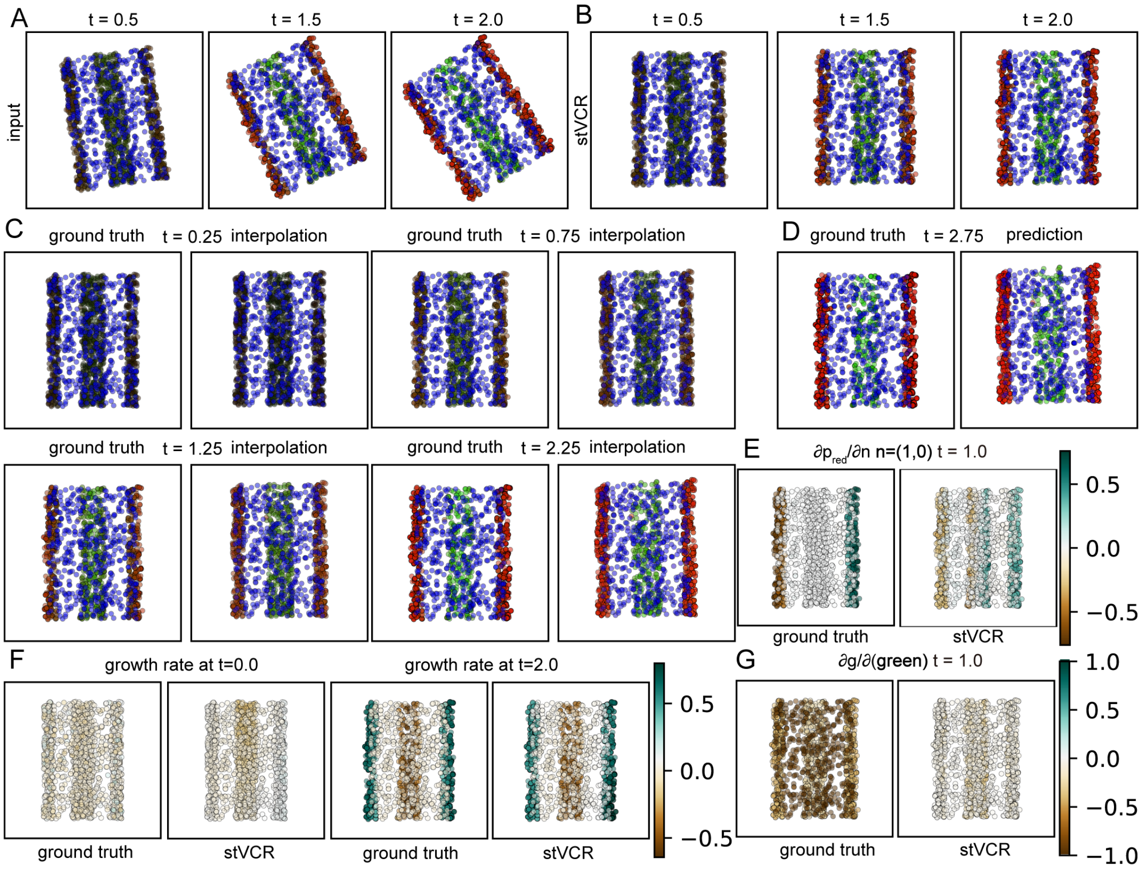

**Fig. S1. Benchmark on the first simulated time-series ST data, related to Fig. 2. A.** Input data at t=0.5, 1.5 and 2.0. The color was determined by the expression of three genes *Red*, *Green* and *Blue*. **B.** The aligned results at $t = 0.5$, 1.5 and 2.0 of cells at $t = 0.0$ using stVCR. **C.** Results of stVCR interpolation at $t = 0.25$, 0.75, t=1.25 and t=2.25 and comparison with ground truth. Left: ground truth; Right: stVCR. **D.** Same as **D**, but for results of stVCR prediction at $t = 2.75$. **E.** Derivative of *Red* gene velocity with respect to direction $n = (1, 0)$ at $t = 1.0$ of true dynamics and learned dynamics. **F.** Growth rates of cells at $t = 0.0$ and 2.0 of true dynamics and learned dynamics. **G.** Same as **E**, but for derivative of growth rate with respect to *Green* gene on cells.

#### Supplementary Figure S2

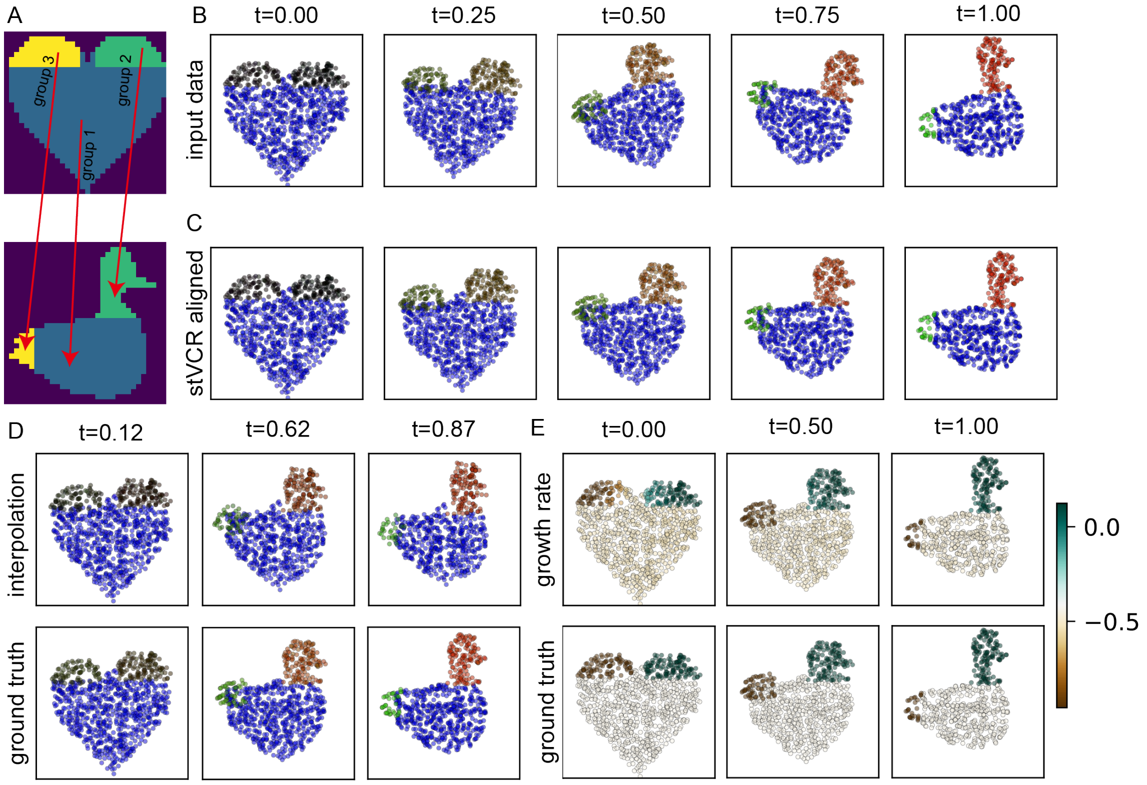

**Fig. S2. Benchmark on the second simulated time-series ST data, related to Fig. 2.** **A.** Dynamic diagram of cell evolution over time. The evolution of genes is similar to Fig. 2. The migration of spatial position is shown by the arrow. **B.** Input data. The color was determined by the expression of three genes *Red*, *Green* and *Blue*. **C.** The aligned results using stVCR. **D.** Results of stVCR interpolation at $t = =0.12$, $t = =1.62$ and $t = =0.87$ and comparison with ground truth. Left: ground truth; Right: stVCR. **E.** Growth rates of cells at $t = 0.00$, $t = 0.50$ and $t = 1.00$ of true dynamics and learned dynamics.

#### Supplementary Figure S3

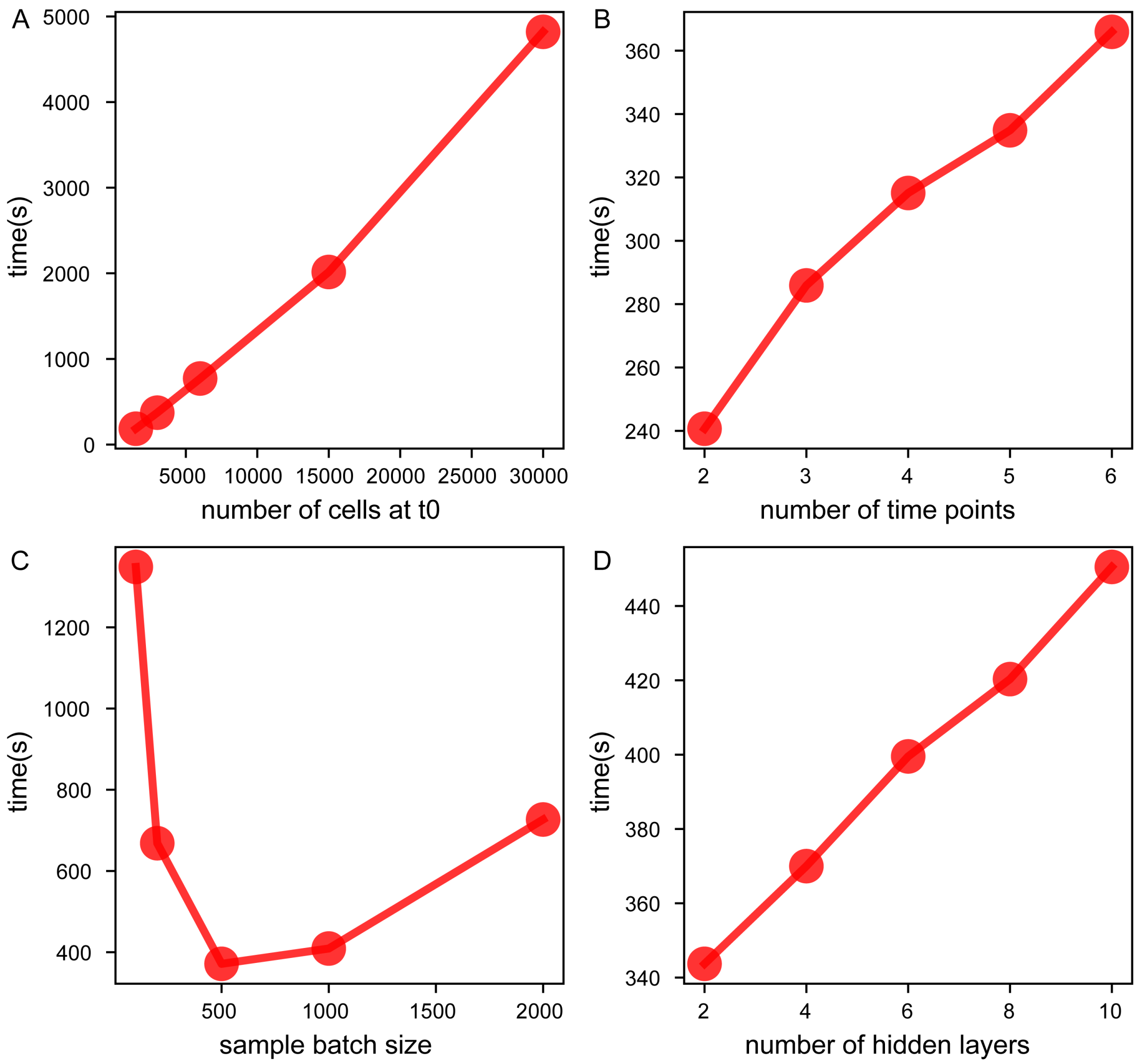

**Fig. S3. Time-consuming comparison of the stVCR on the simulated dataset, related to Fig. 2.** The standard settings are an initial number of cells of 3000, a total of 6 time points, a model size of 4 hidden layers of 128 neurons per layer and a batch size of 500. **A.** Datasets of different scales (characterized by the initial number of cells). **B.** Different number of time points. **C.** Different sample batch sizes. **D.** Different model sizes (characterized by the number of hidden layers).

#### Supplementary Figure S4

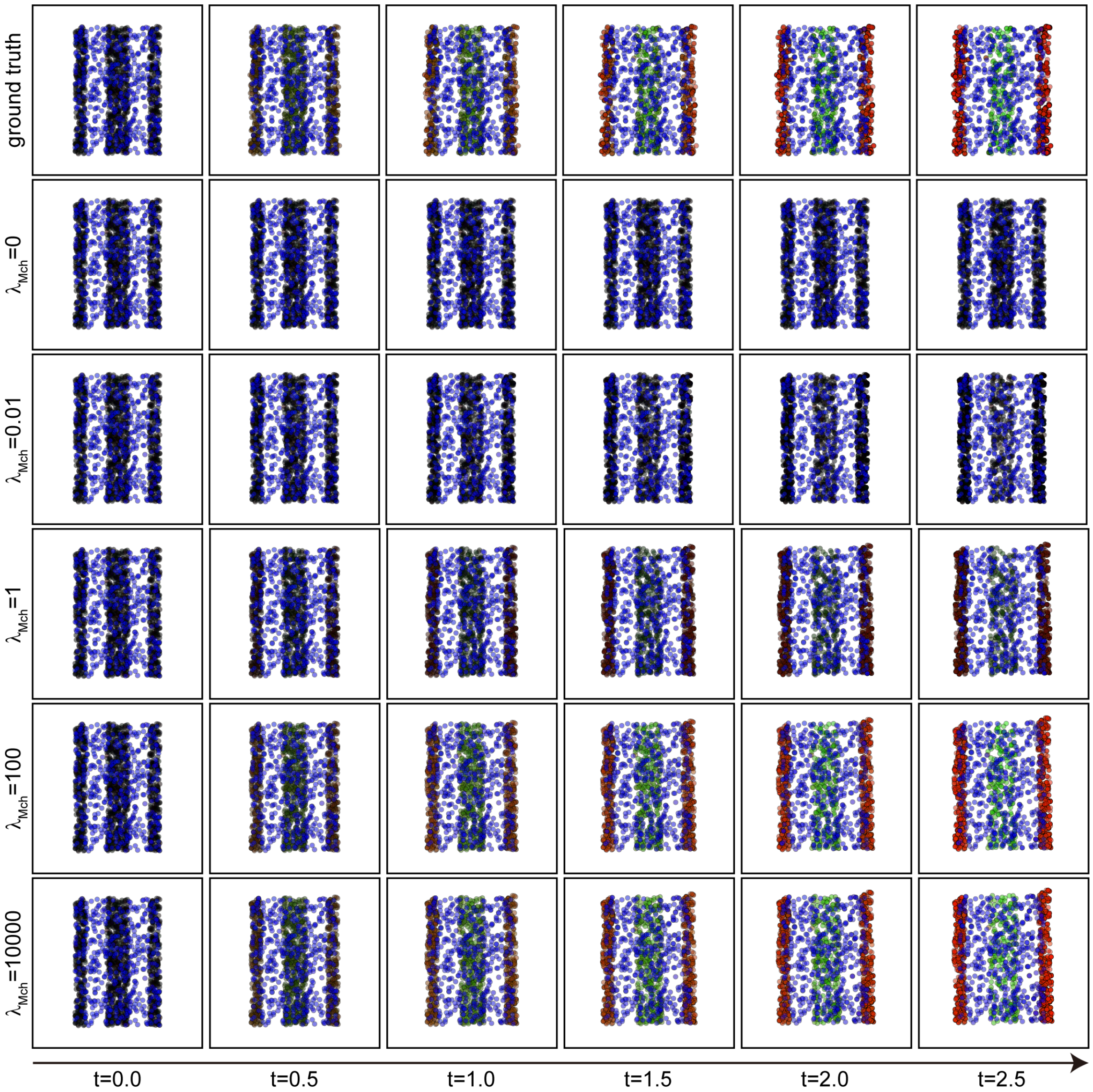

**Fig. S4. Robustness analysis of parameter $\lambda_{\text{Mch}}$ of the stVCR on the simulated dataset, related to Fig. 2.** Parameter $\lambda_{\text{Mch}}$ measures the importance of the loss term $\mathcal{L}_{\text{Mch}}$. Larger values of parameter $\lambda_{\text{Mch}}$ imply that $\mathcal{L}_{\text{Mch}}$ is more important.

363 ## Supplementary Figure S5

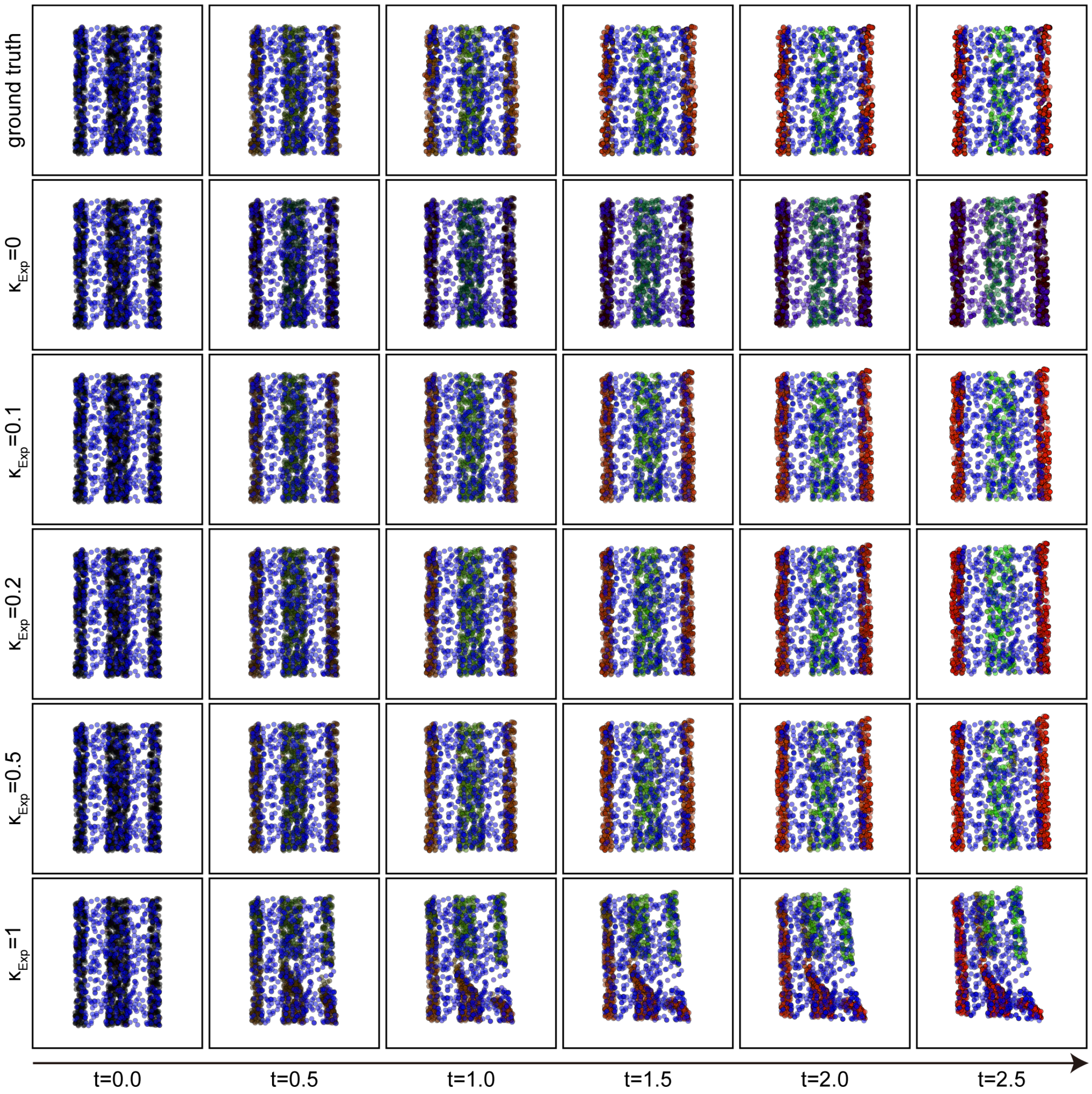

**Fig. S5. Robustness analysis of parameter $\kappa_{\mathbf{Exp}}$ of the stVCR on the simulated dataset, related to Fig. 2.** Parameter $\kappa_{\text{Exp}}$ weighs the importance of gene expression and spatial coordinates in the matching term, when $\kappa_{\text{Exp}} = 0$ means that only spatial coordinates is considered, and conversely when $\kappa_{\text{Exp}} = 1$ means that only gene expression are considered.

#### Supplementary Figure S6

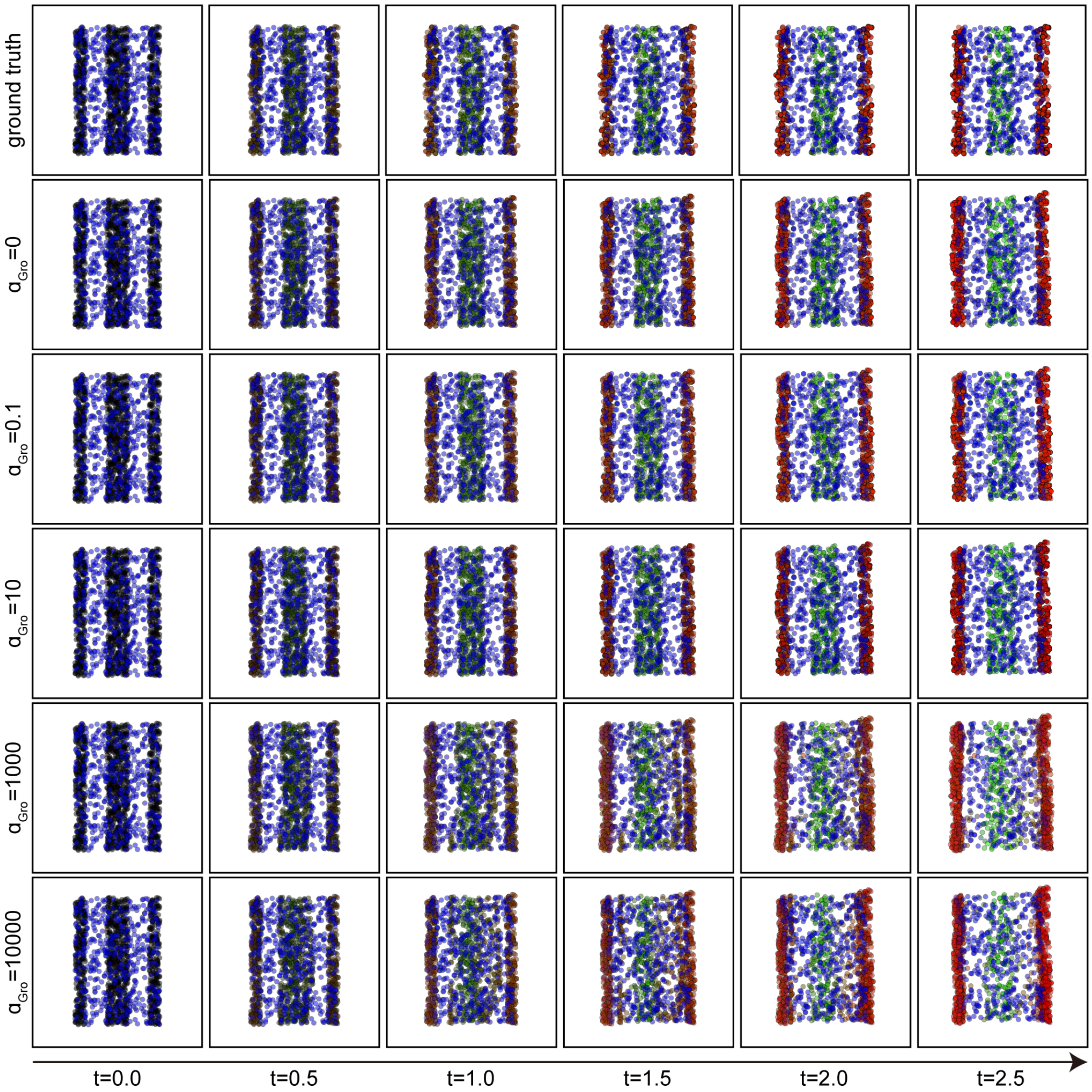

**Fig. S6. Robustness analysis of parameter $\alpha_{\mathbf{Gro}}$ of the stVCR on the simulated dataset, related to Fig. 2.** Parameter $\alpha_{\mathrm{Gro}}$ measures the cost of cell growth, with lower values promoting cells that can divide and apoptose more flexibly, and higher values promoting all cells to have the same growth rate (equivalently, balanced OT).

#### Supplementary Figure S7

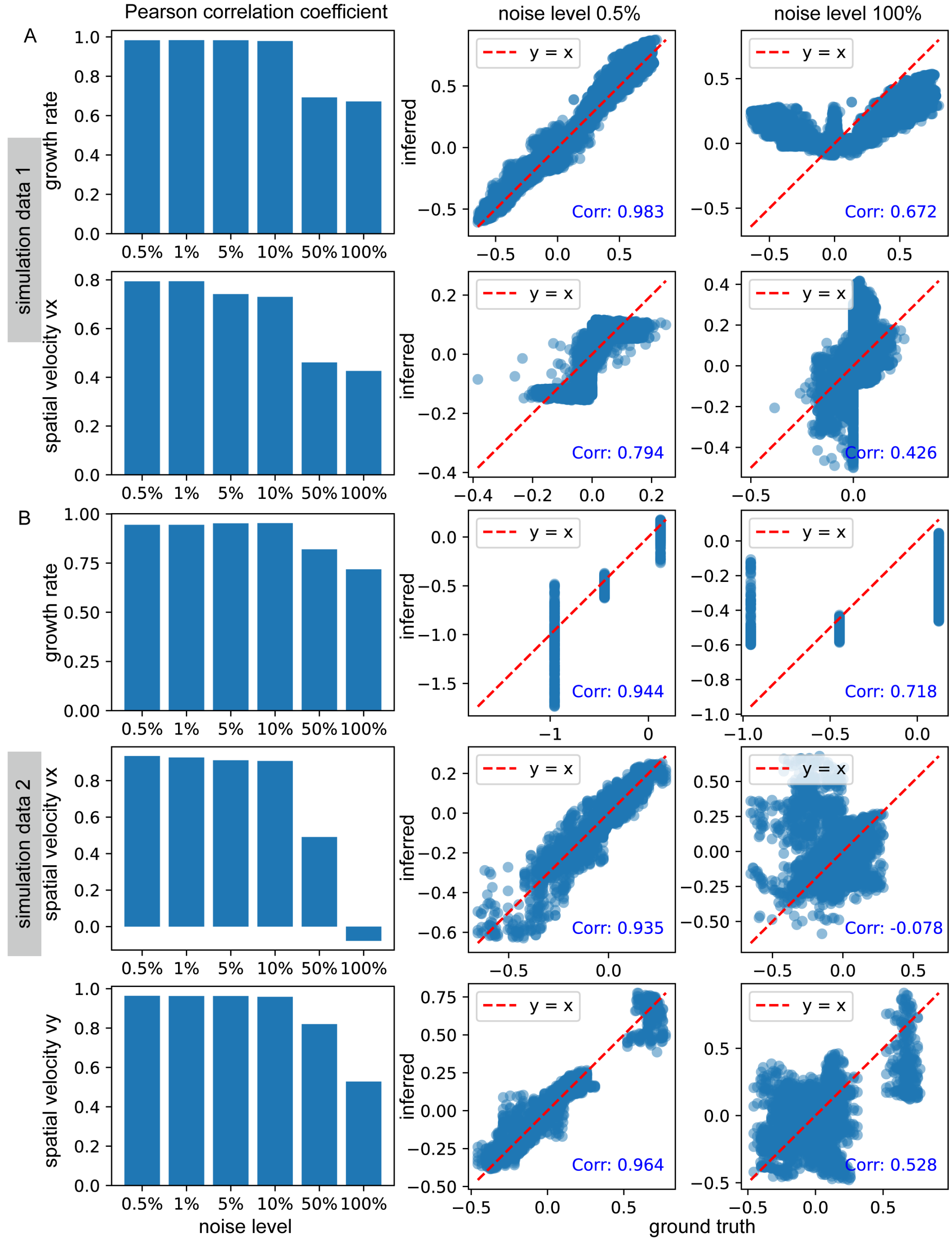

**Fig. S7. Gene and spatial coordinate perturbation benchmark experiment, related to Fig. 2.** From left to right are Pearson correlation between the inferred results and the ground truth under different noise levels and scatter plots at noise levels of 0.5% and 100%. The noise level here refers to Gaussian noise with different percentages of mean gene expression and mean spatial coordinate norm. **A.** Results of simulation data 1. **B.** Similar to **A**, but simulated data 2.

### Supplementary Figure S8

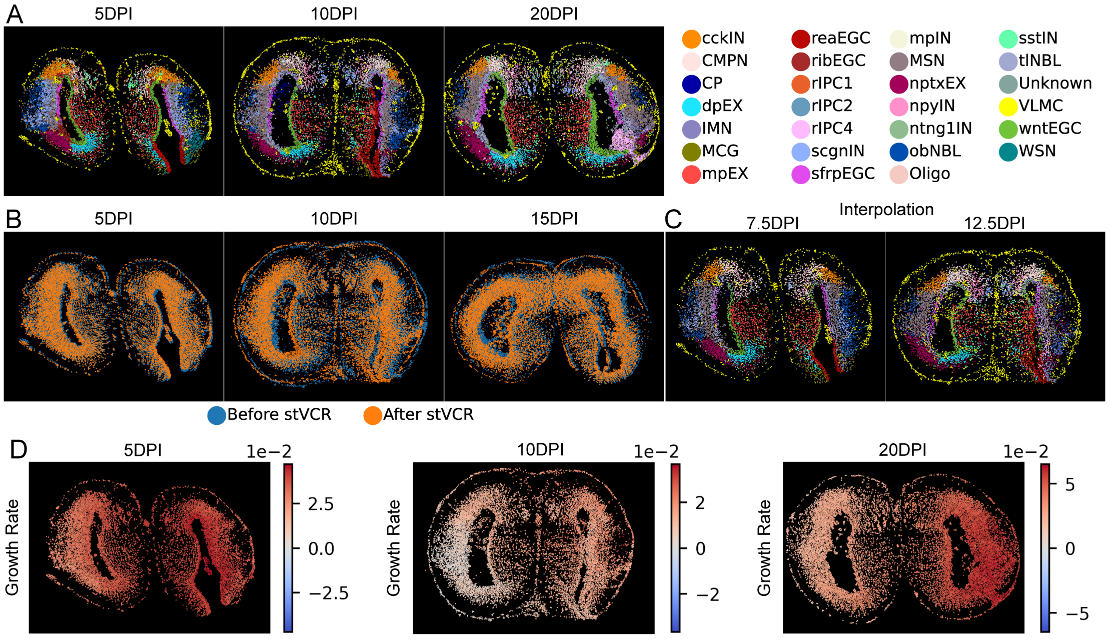

**Fig. S8. stVCR reconstructs axolotl brain regeneration, related to Fig. 3. A.** stVCR aligns the spatial coordinates of data at different time points to the same coordinate system. Left: 5DPI; Middle: 10DPI; Right: 20DPI. Cell type annotations come from the original study. **B.** Comparison of spatial coordinates before and after stVCR alignment. Left: 5DPI; Middle: 10DPI; Right: 15DPI. **C.** stVCR interpolation at 7.5DPI and 12.5DPI. Cell type annotations come from the stVCR's time-dependent classifier. **D.** stVCR cell growth rate at 5DPI, 10DPI and 20DPI data.

#### Supplementary Figure S9

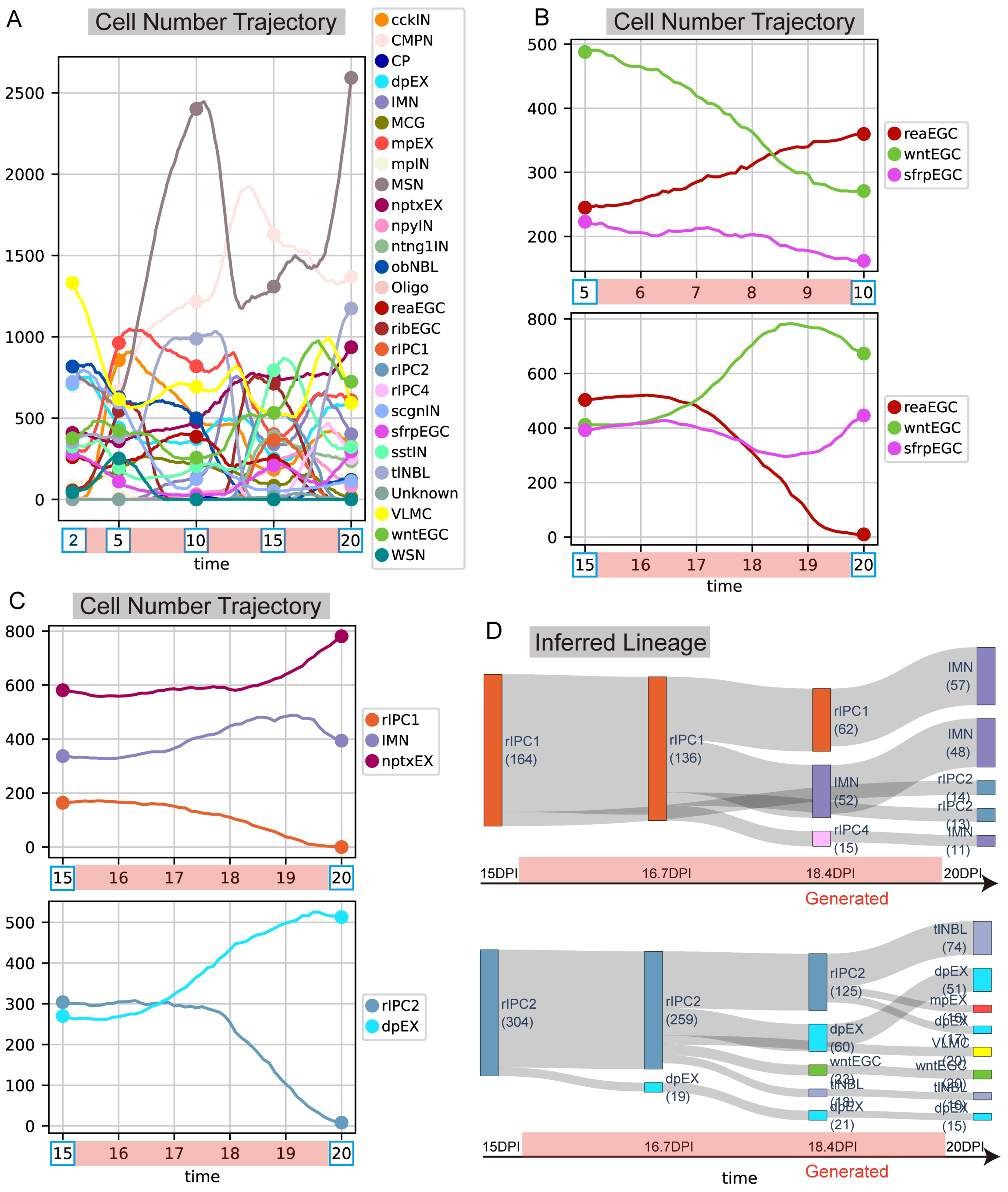

**Fig. S9. Cell number and developmental lineage inference of axolotl brain regeneration, related to Fig. 3. A.** Cell number of different cell types over time 2∼20 DPI. **B.** Similar to **A**, but for part EGC-type cells and part time. Top: 5∼10 DPI; Bottom: 15∼20 DPI. **C.** Similar to **A**, but for part cell types and time 15∼20 DPI. Top: rIPC1, IMN, nptxEX; Bottom: rIPC2 and dpEX. **D.** Time-varying cell types developmental lineages of rIPC1 (Top) and rIPC2 (Bottom) 15∼20 DPI.

#### Supplementary Figure S10

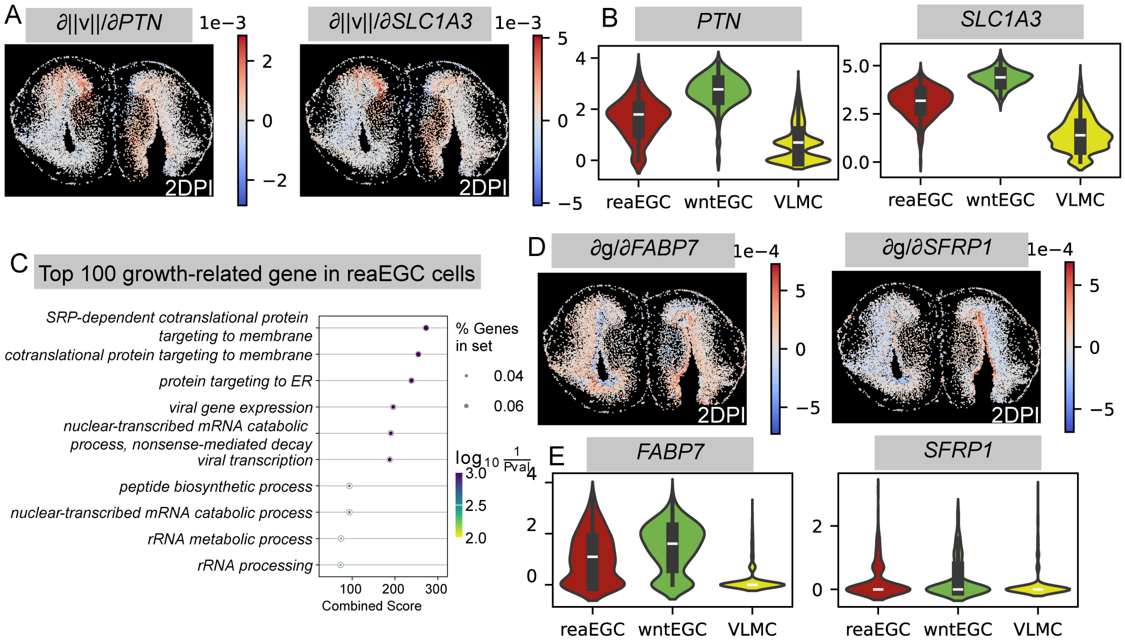

**Fig. S10. stVCR gene-level analysis of axolotl brain regeneration, related to Fig. 4. A.** The partial derivative of the norm of spatial velocity $\|v_z\|$ with respect to gene expression. Two example genes *PTN* (Left) and *SLC1A3* (Right). **B.** Violin plots of gene *PTN* (Left) and *SLC1A3* (Right) expression in reaEGC, wntEGC and VLMC cells. **C.** GO biological process enrichment analysis of the top 100 growth-promoting genes in reaEGC cells. **D.** The partial derivative of growth rate $g$ with respect to gene expression. Two example genes in **C**, *FABP7* (Left) and *SFRP1* (Right). **E.** Similar to **B**, but for gene *FABP7* (Left) and *SFRP1* (Right).

369 ## Supplementary Figure S11

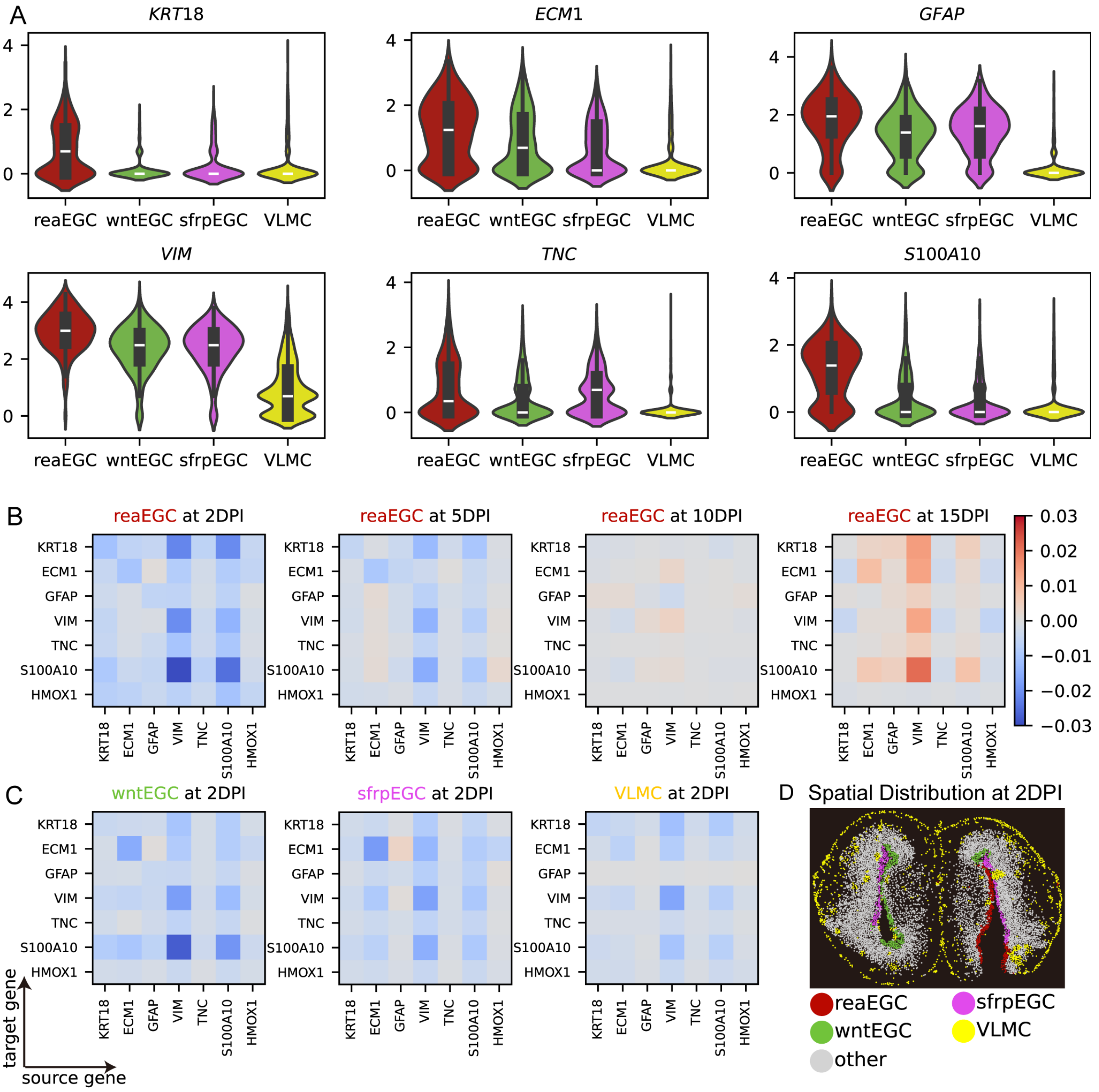

**Fig. S11. stVCR gene regulation analysis of axolotl brain regeneration, related to Fig. 4. A.** Violin plots of genes highly expressed in reaEGC. From left to right and from top to bottom are *KRT18*, *ECM1*, *GFAP*, *VIM*, *TNC* and *S100A10*. **B.** Heat map of gene regulatory relationships in reaEGC cells. From left to right are 2DPI, 5DPI, 10DPI and 10DPI. **C.** Heat map of gene regulatory relationships in wntEGC (Left), sfrpEGC (Middle) and VLMC (Right) cells at 2DPI. **D.** Spatial distribution of reaEGC, wntEGC, sfrpEGC and VLMC cells at 2 DPI.

#### Supplementary Figure S12

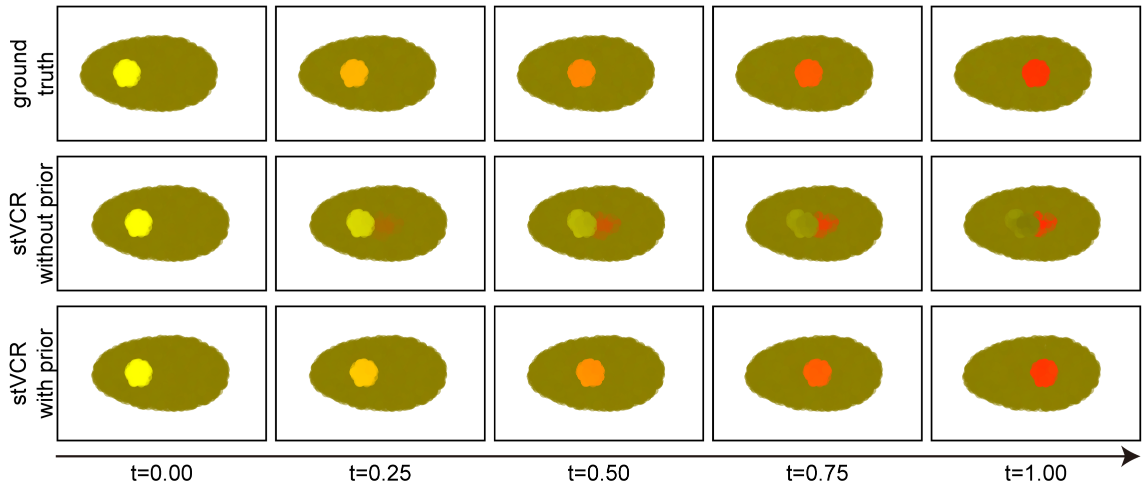

**Fig. S12. Comparison of the interpolation results of stVCR witout biological prior, stVCR with biological prior and ground truth on the simulated data with only two time points, related to Fig. 5.**

371 ## Supplementary Figure S13

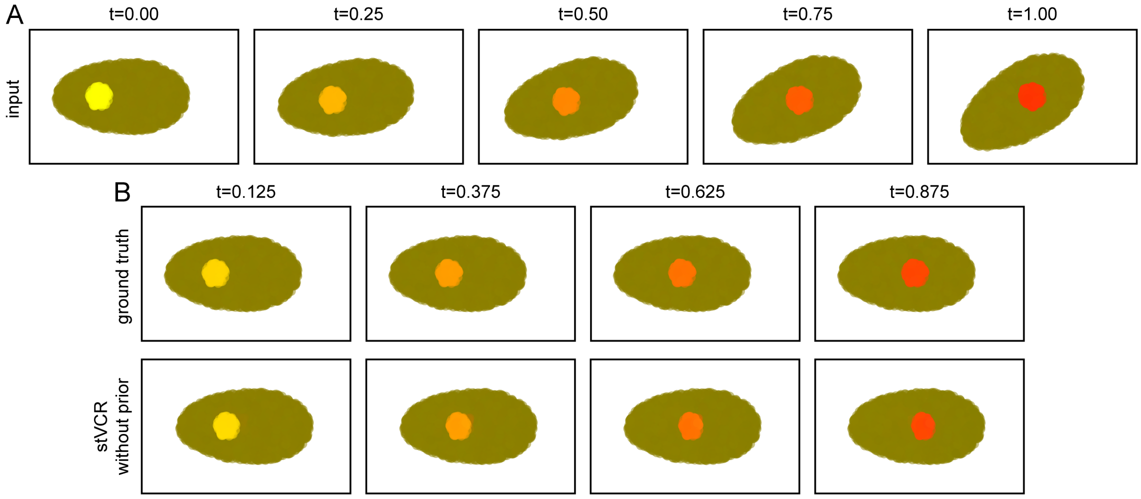

**Fig. S13. Results of stVCR witout biological prior on the simulated data with multiple time points, related to Fig. 5. A.** Input data for 5 time points of simulated data in **Fig. 5A** and **B**. From left to right are t=0.00, t=0.25, t=0.50, t=0.75 and t=1.00. **B.** Interpolation results at t=0.125, t=0.375, t=0.625 and t=0.875 of stVCR without biological prior compared to the ground truth. Top: ground truth. Bottom: stVCR without prior.

#### Supplementary Figure S14

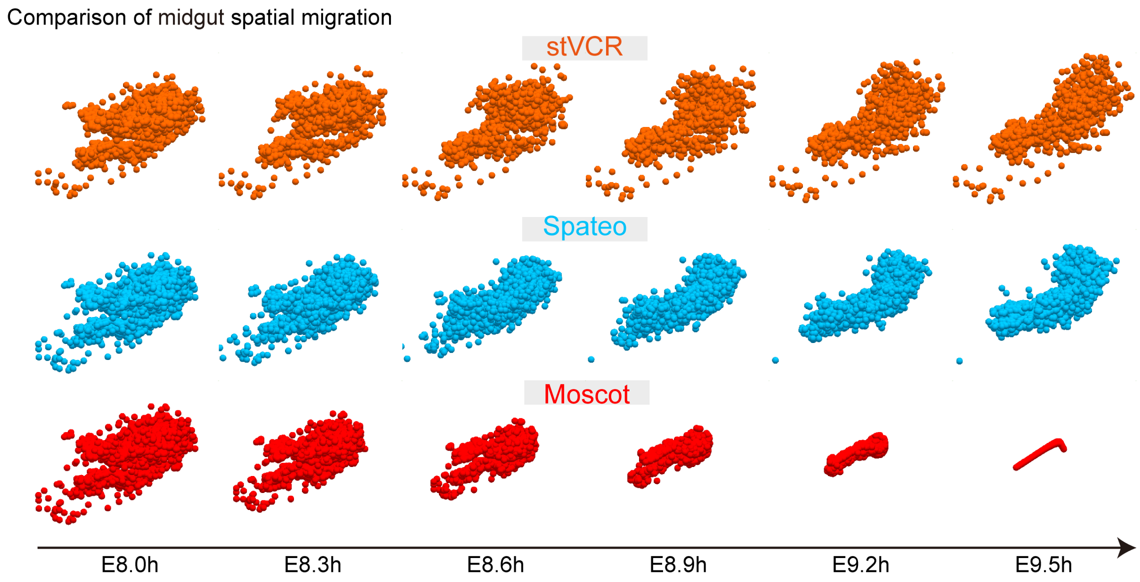

**Fig. S14. Comparison of spatial migration trajectories of midgut cells, related to Fig. 5.** From left to right: E8.0h, E8.3h, E8.6h, E8.9h, E9.2h and E9.5h. From top to bottom: stVCR, Spateo [10] and Moscot [11].

### Supplementary Videos

#### Supplementary Video S1

Comparison of stVCR reconstruction results with ground truth in the simulated data in Figure 2.

Play Video S1

#### Supplementary Video S2

Comparison of stVCR reconstruction results with ground truth in the simulated data in Figure S2.

Play Video S2

#### Supplementary Video S3-S5

Robustness analysis of hyper-parameter $\lambda_{\mathrm{Mch}}$ (S3), $\kappa_{\mathrm{Exp}}$ (S4) and $\alpha_{\mathrm{Gro}}$ (S5) in the simulated data in Figure 2.

Play Video S3 Play Video S4 Play Video S5

#### Supplementary Video S6

stVCR reconstructs axolotl brain regeneration.

Play Video S6

#### Supplementary Video S7-S8

Comparison of stVCR with (S7) and without (S8) biological prior reconstruction results with ground truth in the simulated data in Figure 5.

Play Video S7 Play Video S8

#### Supplementary Video S9

Comparison of stVCR without biological prior reconstruction results with ground truth in the simulated data in Figure S13.

Play Video S9

#### Supplementary Video S10

stVCR reconstructs development of 3D Drosophila embryo with biological prior and spatial structure preservation prior

Play Video S10

#### Supplementary Video S11-S13

Comparison of spatial migration trajectories of CNS cells inferred from stVCR (S11), Spateo [10] (S12) and Moscot [11] (S13).

Play Video S11 Play Video S12 Play Video S13

#### Supplementary Video S14-S16

Comparison of spatial migration trajectories of midgut cells inferred from stVCR (S14), Spateo [10] (S15) and Moscot [11] (S16).

Play Video S14 Play Video S15 Play Video S16

### Supplementary Tables

#### Supplementary Table S1 Capabilities of methods designed for time-series ST data

| **Methods** | **stVCR** | **Moscot** [11] | **Spateo** [10] | **DeST-OT** [12] |
|---|---|---|---|---|
| Cell transition | Yes | Static | Static | Static |
| Cell migration | Yes | Static | Static | Static |
| Cell growth | Yes | Prior | No | Static |
| Prediction | Yes | No | No | No |
| GRN reconstruction | Yes | No | No | No |
| Gene-space interaction | Yes | No | No | No |
| Space-space interaction | Yes | No | Yes | No |
| Growth promotion/ inhibition gene discovery | Yes | No | No | No |

**Table S1. Summary of reconstruction methods for time series spatial transcriptome data. The "Static" here refers to the result obtained by linear interpolation.** The "Prior" here refers to derived from prior knowledge rather than inferred from data.

#### Supplementary Table S2-S5 Quantitative comparison results of the first simulated dataset

| algorithm ↓ time points → | t=0.0 | t=0.5 | t=1.0 | t=1.5 | t=2.0 | t=2.5 |
|---|---|---|---|---|---|---|
| **DeST-OT** [12] | *0.0535* | 0.1652 | 0.1811 | 0.1728 | 0.1824 | N/A |
| **Spateo** [10] | N/A | N/A | N/A | N/A | N/A | N/A |
| **Moscot** [11] | N/A | N/A | N/A | N/A | N/A | N/A |
| **SF2M** [13] | N/A | N/A | N/A | N/A | N/A | N/A |
| **stVCR w/o spatial (that is TIGON** [14]**)** | **0.0520** | 0.1642 | 0.1894 | 0.1504 | 0.1112 | 0.1046 |
| **stVCR w/o gene** | 0.0733 | 0.1062 | 0.1287 | 0.1573 | 0.1758 | 0.1794 |
| **stVCR w/o growth** | N/A | N/A | N/A | N/A | N/A | N/A |
| **stVCR w/o alignment** | 0.0849 | *0.0593* | *0.0561* | **0.0498** | **0.0482** | **0.0520** |
| **stVCR** | 0.0845 | **0.0497** | **0.0491** | *0.0548* | *0.0616* | *0.0694* |

**Table S2.** The mean absolute error of **growth rate** inference at different observed time points in the first simulated dataset.

| algorithm ↓ time points → | t=0.0 | t=0.5 | t=1.0 | t=1.5 | t=2.0 | t=2.5 |
|---|---|---|---|---|---|---|
| **DeST-OT** [12] | **0.0087** | **0.0412** | **0.0328** | **0.0307** | **0.0247** | N/A |
| **Spateo** [10] | 0.1117 | 0.1801 | 0.2461 | 0.2504 | 0.2245 | N/A |
| **Moscot** [11] | 0.2505 | 0.2779 | 0.3066 | 0.3147 | 0.2929 | N/A |
| **SF2M** [13] | N/A | N/A | N/A | N/A | N/A | N/A |
| **stVCR w/o spatial (that is TIGON** [14]**)** | N/A | N/A | N/A | N/A | N/A | N/A |
| **stVCR w/o gene** | 0.0631 | 0.0576 | 0.0573 | 0.0589 | 0.0648 | *0.0728* |
| **stVCR w/o growth** | 0.0973 | 0.0867 | 0.1016 | 0.1151 | 0.1410 | 0.2002 |
| **stVCR w/o alignment** | 0.3982 | 0.3924 | 0.4018 | 0.4178 | 0.4439 | 0.4687 |
| **stVCR** | *0.0560* | *0.0495* | *0.0478* | *0.0486* | *0.0548* | **0.0633** |

**Table S3.** The mean RMSE of **spatial velocity** inference at different observed time points in the first simulated dataset.

| **algorithm ↓ leave time point →** | **t=0.5** | **t=1.0** | **t=1.5** | **t=2.0** |
|---|---|---|---|---|
| **SF2M** [13] | 0.0620 | 0.1018 | 0.3889 | 1.1397 |
| **stVCR w/o spatial (that is TIGON** [14]) | **0.0215** | **0.0462** | *0.0551* | *0.1023* |
| **stVCR w/o gene** | N/A | N/A | N/A | N/A |
| **stVCR w/o growth** | 0.0304 | 0.1260 | 0.2710 | 0.2176 |
| **stVCR w/o alignment** | 0.0401 | 0.0547 | 0.0482 | 0.1220 |
| **stVCR** | *0.0262* | *0.0476* | **0.0432** | **0.0603** |

**Table S4.** Leave-one-time point-out testing of dynamics interpolation methods measuring the error between the predicted **gene expression** and ground truth left out time point using the 2-Wasserstein in the first simulated dataset. Each column represents the time point that was not used during training.

| **algorithm ↓ leave time point →** | **t=0.5** | **t=1.0** | **t=1.5** | **t=2.0** |
|---|---|---|---|---|
| **SF2M** [13] | N/A | N/A | N/A | N/A |
| **stVCR w/o spatial (that is TIGON** [14]) | N/A | N/A | N/A | N/A |
| **stVCR w/o gene** | N/A | N/A | N/A | N/A |
| **stVCR w/o growth** | *0.0235* | *0.0597* | *0.0888* | *0.0649* |
| **stVCR w/o alignment** | 0.0622 | 0.1762 | 0.3186 | 0.5804 |
| **stVCR** | **0.0129** | **0.0222** | **0.0273** | **0.0619** |

**Table S5.** Leave-one-time point-out testing of dynamics interpolation methods measuring the error between the predicted **gene and spatial coordinates interpolation** and ground truth left out time point using the 2-Wasserstein in the first simulated dataset. Each column represents the time point that was not used during training.

#### Supplementary Table S6-S9 Quantitative comparison results of the second simulated dataset

| algorithm ↓ time points → | t=0 | t=0.25 | t=0.50 | t=0.75 | t=1.0 |
|---|---|---|---|---|---|
| **DeST-OT** [12] | 0.1643 | 0.1573 | 0.1962 | *0.0748* | N/A |
| **Spateo** [10] | N/A | N/A | N/A | N/A | N/A |
| **Moscot** [11] | N/A | N/A | N/A | N/A | N/A |
| **SF2M** [13] | N/A | N/A | N/A | N/A | N/A |
| **stVCR w/o spatial (that is TIGON** [14]) | 0.1844 | 0.1715 | 0.1546 | 0.1366 | 0.1272 |
| **stVCR w/o gene** | 0.1359 | 0.1368 | 0.1350 | 0.1230 | 0.1200 |
| **stVCR w/o growth** | N/A | N/A | N/A | N/A | N/A |
| **stVCR w/o alignment** | *0.0987* | *0.0875* | *0.0908* | 0.1239 | *0.0920* |
| **stVCR** | **0.0854** | **0.0701** | **0.0718** | **0.0738** | **0.0655** |

**Table S6.** The mean absolute error of **growth rate** inference at different observed time points in the second simulated dataset.

| algorithm ↓ time points → | t=0 | t=0.25 | t=0.50 | t=0.75 | t=1.0 |
|---|---|---|---|---|---|
| **DeST-OT** [12] | 0.1631 | 0.1584 | 0.1557 | 0.1511 | N/A |
| **Spateo** [10] | 0.1241 | 0.1284 | 0.1572 | 0.1631 | N/A |
| **Moscot** [11] | 0.1526 | 0.1474 | 0.1737 | 0.1972 | N/A |
| **SF2M** [13] | N/A | N/A | N/A | N/A | N/A |
| **stVCR w/o spatial (that is TIGON** [14]) | N/A | N/A | N/A | N/A | N/A |
| **stVCR w/o gene** | 0.0884 | 0.0838 | 0.0850 | 0.0822 | *0.0841* |
| **stVCR w/o growth** | *0.0738* | *0.0679* | *0.0625* | *0.0679* | 0.0986 |
| **stVCR w/o alignment** | 0.1240 | 0.1256 | 0.1615 | 0.1359 | 0.5246 |
| **stVCR** | **0.0486** | **0.0465** | **0.0459** | **0.0469** | **0.0504** |

**Table S7.** The mean RMSE of **spatial velocity** inference at different observed time points in the second simulated dataset.

| **algorithm ↓ leave time point →** | **t=0.25** | **t=0.5** | **t=0.75** |
|---|---|---|---|
| **SF2M** [13] | *0.0498* | 0.1380 | 0.3211 |
| **stVCR w/o spatial (that is TIGON** [14]**)** | **0.0323** | **0.0134** | **0.0331** |
| **stVCR w/o gene** | N/A | N/A | N/A |
| **stVCR w/o growth** | 0.0578 | 0.1001 | 0.1501 |
| **stVCR w/o alignment** | 0.0558 | 0.0720 | *0.0673* |
| **stVCR** | 0.0527 | *0.0574* | 0.0823 |

**Table S8.** Leave-one-time point-out testing of dynamics interpolation methods measuring the error between the predicted **gene expression** and ground truth left out time point using the 2-Wasserstein in the second simulated dataset. Each column represents the time point that was not used during training.

| **algorithm ↓ leave time point →** | **t=0.25** | **t=0.5** | **t=0.75** |
|---|---|---|---|
| **SF2M** [13] | N/A | N/A | N/A |
| **stVCR w/o spatial (that is TIGON** [14]**)** | N/A | N/A | N/A |
| **stVCR w/o gene** | N/A | N/A | N/A |
| **stVCR w/o growth** | *0.0098* | *0.0177* | 0.0224 |
| **stVCR w/o alignment** | 0.0115 | 0.0228 | *0.0171* |
| **stVCR** | **0.0071** | **0.0086** | **0.0127** |

**Table S9.** Leave-one-time point-out testing of dynamics interpolation methods measuring the error between the predicted **gene and spatial coordinates interpolation** and ground truth left out time point using the 2-Wasserstein in the second simulated dataset. Each column represents the time point that was not used during training.

#### Supplementary Table S10-S12 Quantitative comparison results of the axolotl regenerative datasets [15]

| algorithm ↓ leave time point → | t=5DPI | t=15DPI |
|---|---|---|
| **SF2M** [13] | 0.2003 | 0.9302 |
| **stVCR w/o spatial (that is TIGON** [14]) | 0.1899 | 0.6930 |
| **stVCR w/o gene** | N/A | N/A |
| **stVCR w/o growth** | 0.1837 | *0.6674* |
| **stVCR w/o alignment** | *0.1799* | 0.7371 |
| **stVCR** | **0.1778** | **0.6614** |

**Table S10.** Leave-one-time point-out testing of dynamics interpolation methods measuring the error between the predicted **gene expression** and ground truth left out time point using the 2-Wasserstein in the axolotl regenerative datasets [15]. Each column represents the time point that was not used during training.

| algorithm ↓ leave time point → | t=5DPI | t=15DPI |
|---|---|---|
| **SF2M** [13] | N/A | N/A |
| **stVCR w/o spatial (that is TIGON** [14]) | N/A | N/A |
| **stVCR w/o gene** | N/A | N/A |
| **stVCR w/o growth** | 0.1096 | *0.3585* |
| **stVCR w/o alignment** | **0.1066** | 0.3945 |
| **stVCR** | *0.1069* | **0.3542** |

**Table S11.** Leave-one-time point-out testing of dynamics interpolation methods measuring the error between the predicted **gene and spatial coordinates interpolation** and ground truth left out time point using the fused rigid body transformation invariant OT (see Eq. (10)) in the axolotl regenerative datasets [15]. Each column represents the time point that was not used during training.

| algorithm → time points ↓ injured state → | stVCR | | DeST-OT [12] | |
|---|---|---|---|---|
| | **inj** | **uninj** | **inj** | **uninj** |
| **t=2DPI** | 0.0078 | 0.0074 | 0.0181 | 0.0189 |
| **t=5DPI** | 0.0386 | 0.0261 | 0.0305 | 0.0303 |
| **t=10DPI** | 0.0134 | 0.0076 | 0.0046 | 0.0049 |
| **t=15DPI** | 0.0197 | 0.0098 | 0.0312 | 0.0319 |
| **t=20DPI** | 0.0484 | 0.0360 | N/A | N/A |

**Table S12.** Comparison of the mean growth rate of cells in injured and uninjured hemispheres in the axolotl regenerative datasets [15].
