## Supplementary figures and images for "stVCR: Spatiotemporal dynamics of single cells"

### video_S1_ablation_rectangle.gif

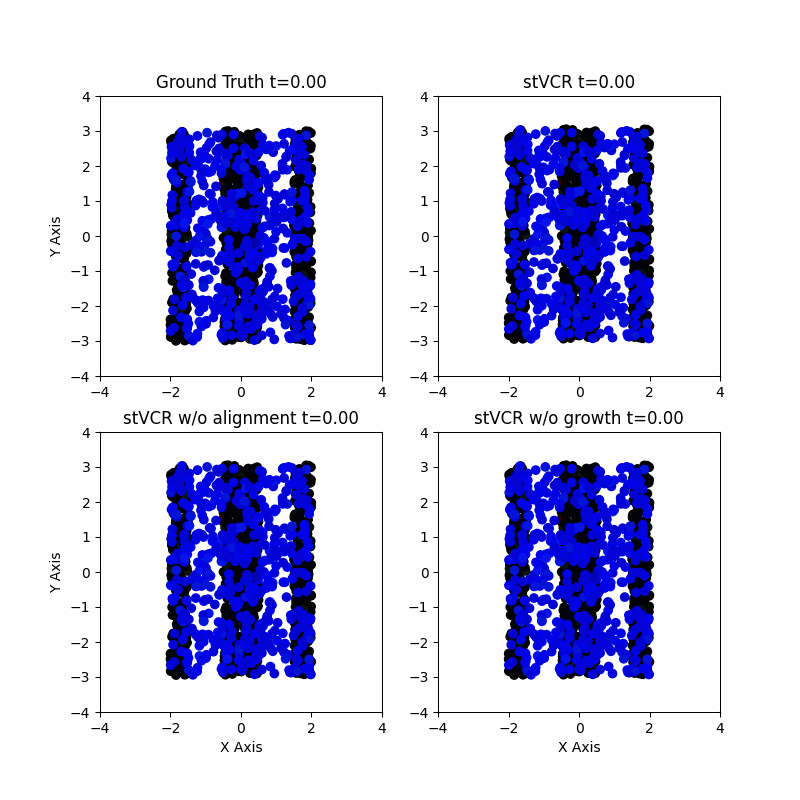

### video_S2_ablation_heart2duck.gif

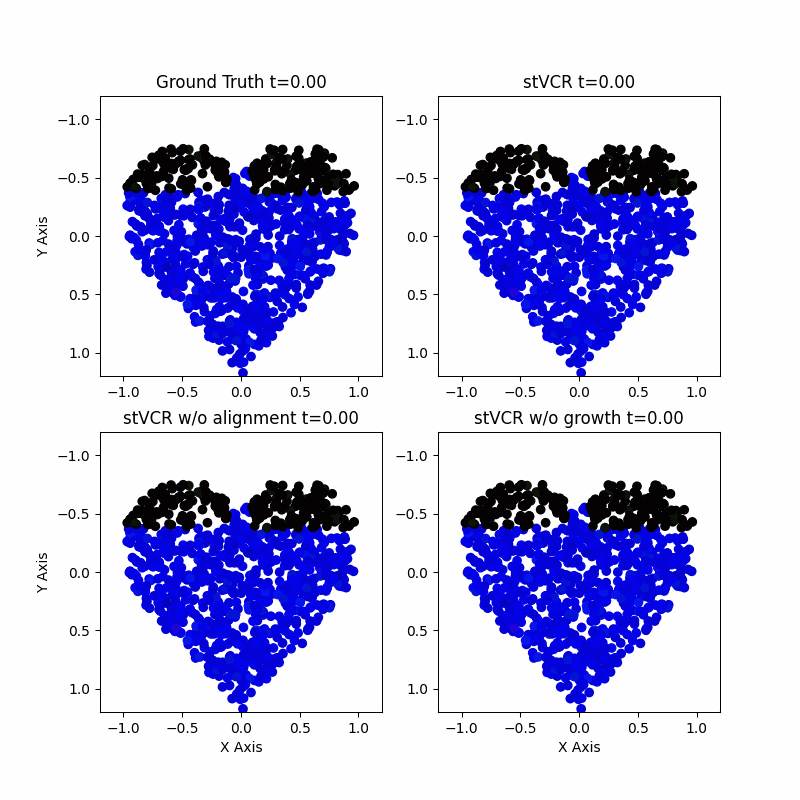

### video_S3_robust_lambda_match.gif

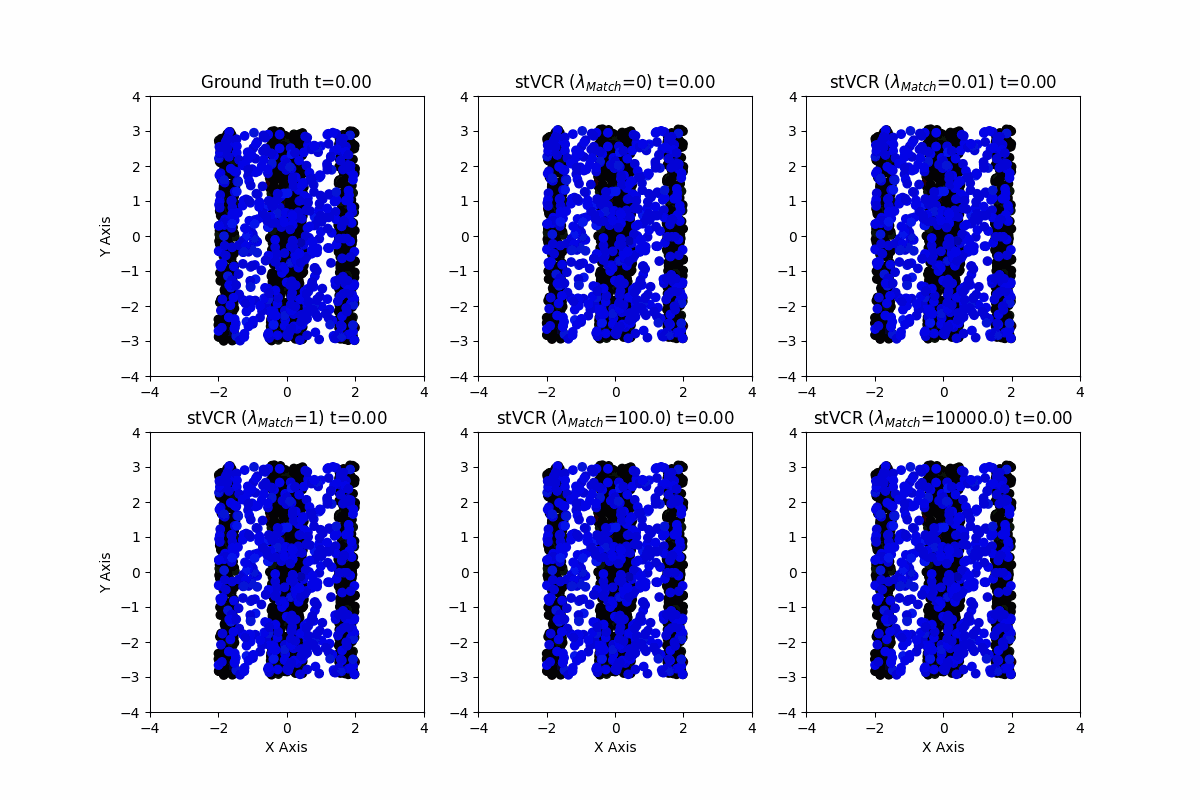

### video_S4_robust_kappa_exp.gif

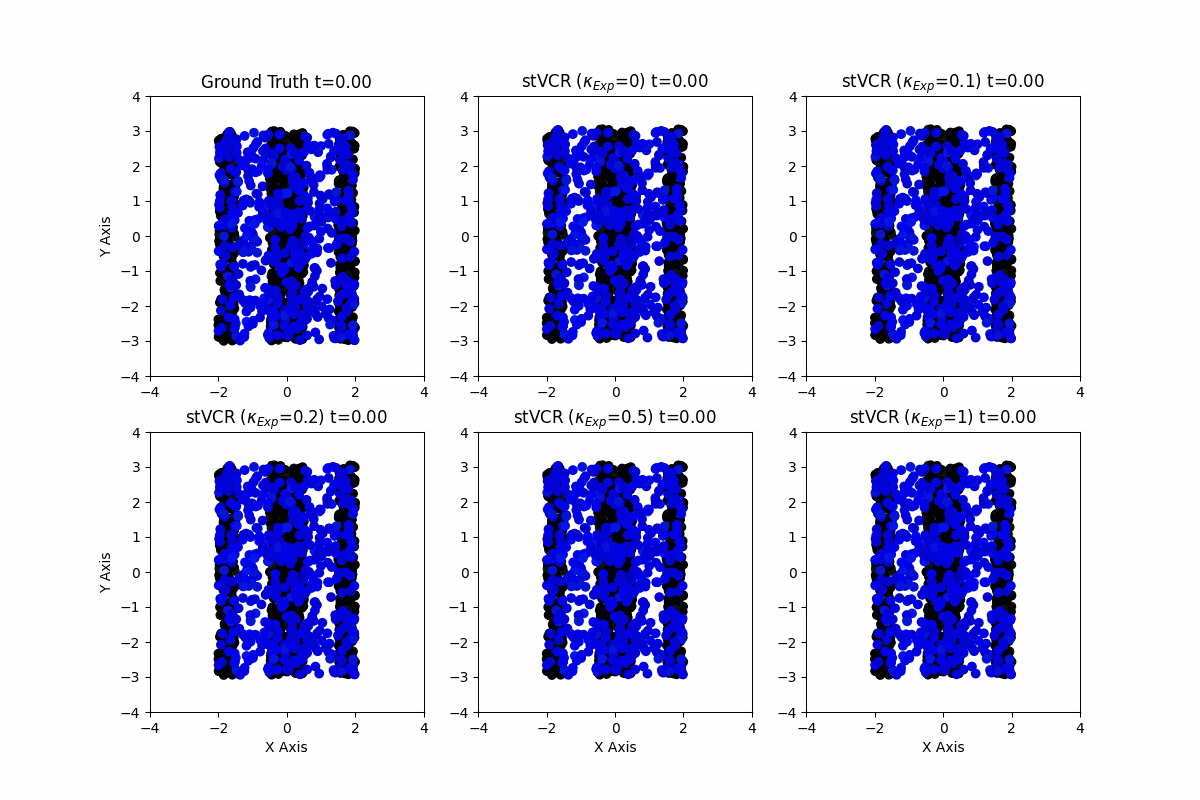

### video_S5_robust_alpha_growth.gif

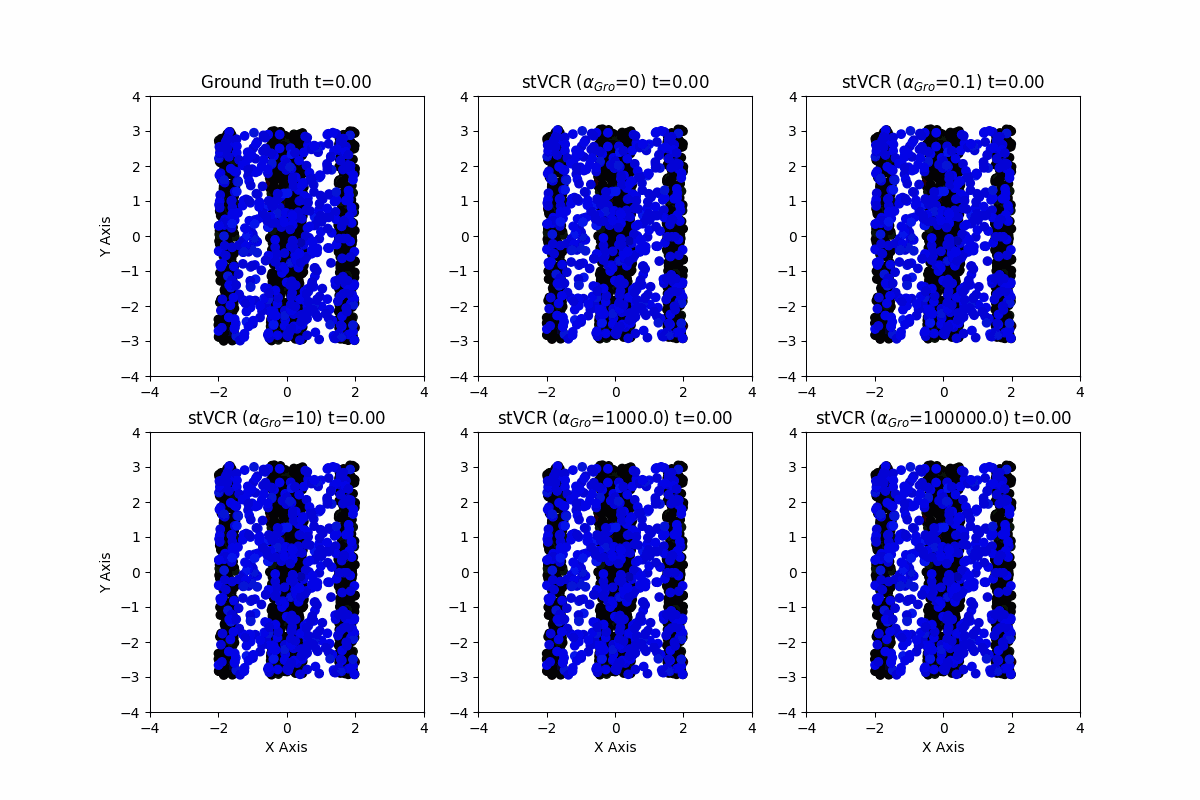

### video_S6_ARTISTA.gif

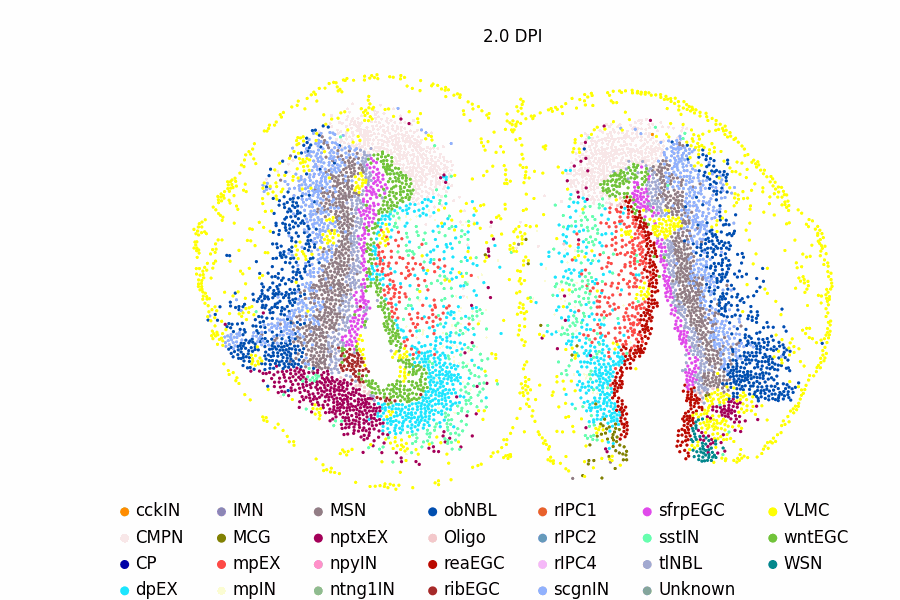

### video_S7_sim_migration_with_bio.gif

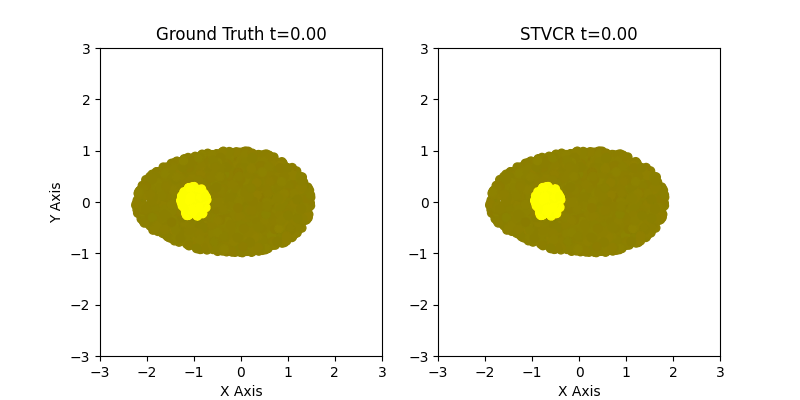

### video_S8_sim_migration_without_bio.gif

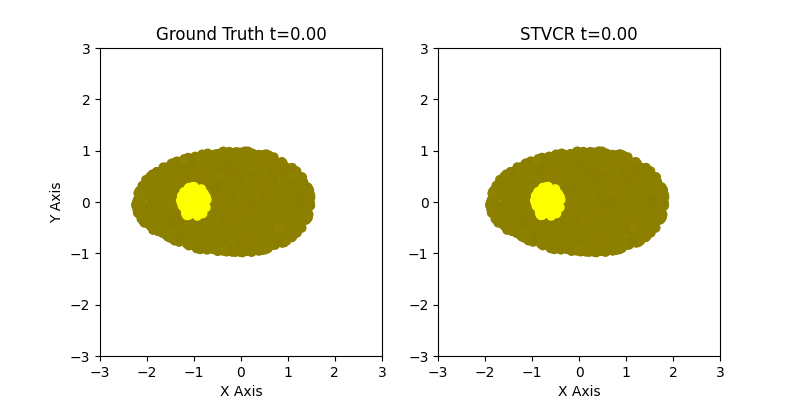

### video_S9_sim_migration_5pts_without_bio.gif

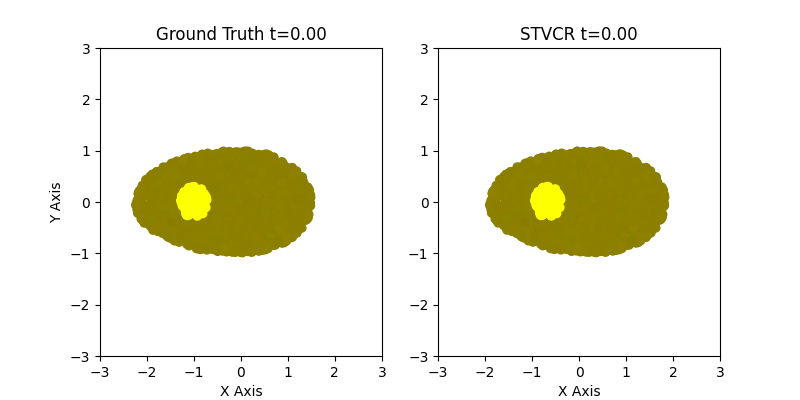

### video_S10_Drosophila3D.gif

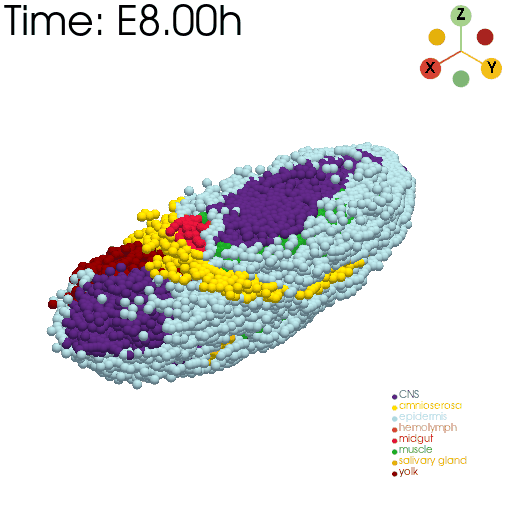

### video_S11_CNS_stVCR.gif

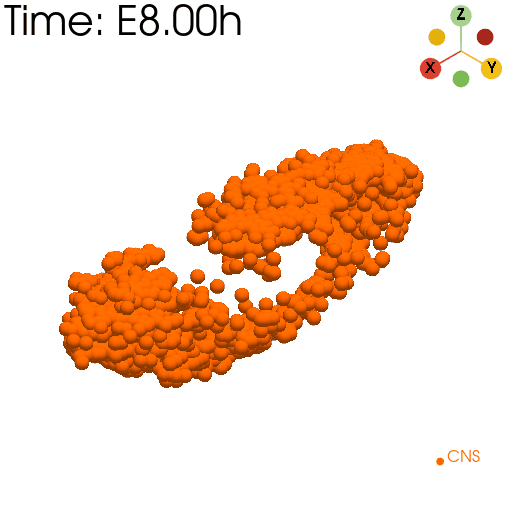

### video_S12_CNS_Spateo.gif

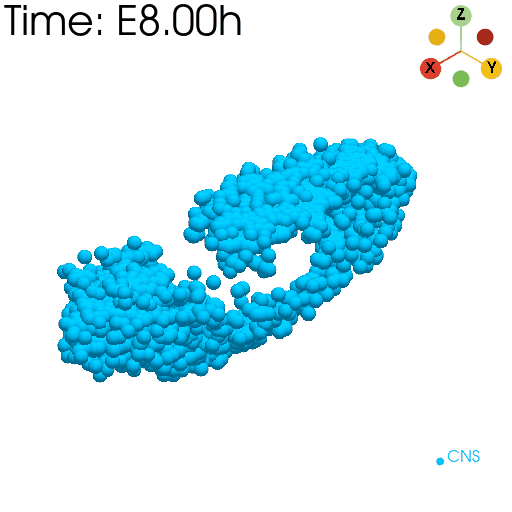

### video_S13_CNS_Moscot.gif

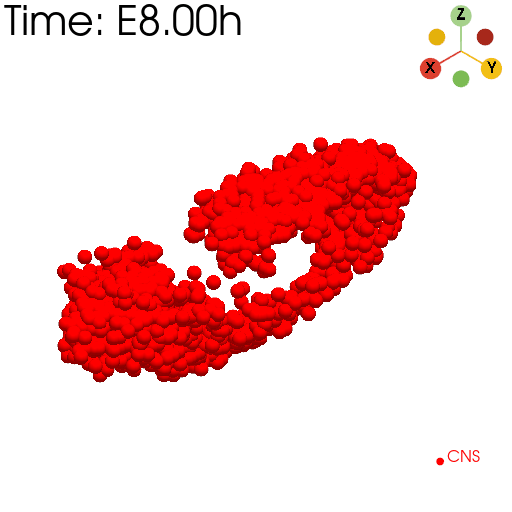

### video_S14_midgut_stVCR.gif

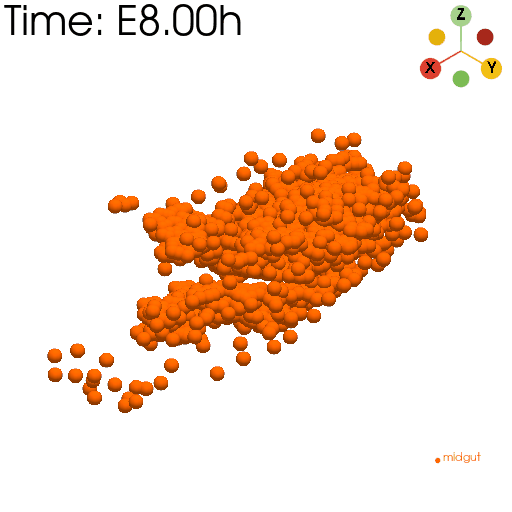

### video_S15_midgut_Spateo.gif

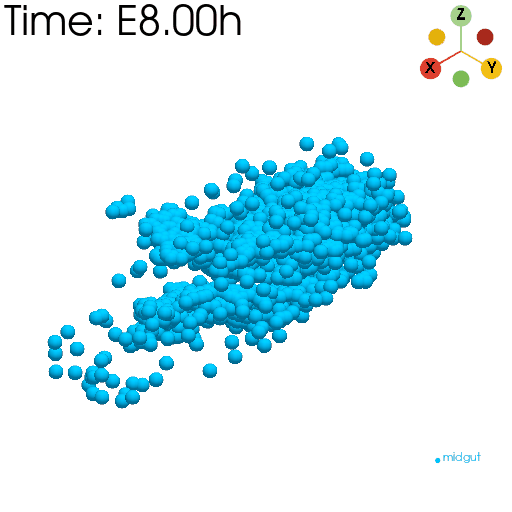

### video_S16_midgut_Moscot.gif

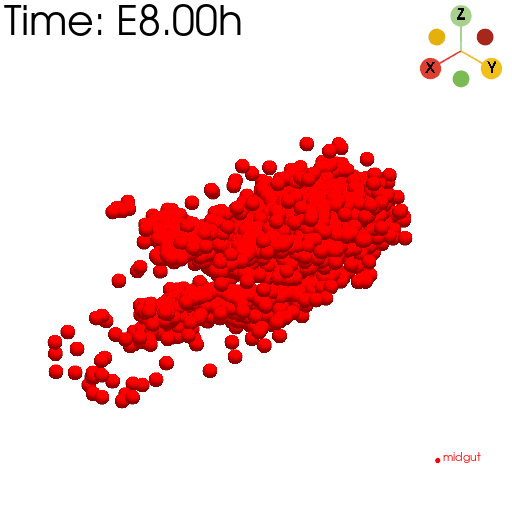
